# Reinforcement learning with evolutionary priors unlocks multi-site enzyme optimization from sparse experimental feedback

**DOI:** 10.64898/2026.10.19.760152

**Authors:** Xiaopeng Xu, Hao Ye, Ziqi Gao, Ruochi Zhang, Xinhai Zhou, Guiting Zhang, Tao Guo, Wenjia He, Chenjie Feng, Bin Xiao, Feifei Hu, Qiangcheng Zeng, Aixi Bai, Xiao Zhang, Jie Wang, Xin Gao

## Abstract

Industrial enzymes underpin sustainable manufacturing, yet natural proteins rarely combine the activity, robustness and application compatibility required for practical use. Engineering such properties requires identifying beneficial combinations of interacting mutations within vast sequence spaces that are too large to search exhaustively and too sparsely sampled to model reliably. Here we introduce AlphaVariant, a reinforcement-learning (RL) framework that couples a generative evolutionary prior with iteratively updated low-N fitness models to enable feedback-driven search of large combinatorial protein sequence spaces. Across six experimentally measured protein fitness landscapes, AlphaVariant ranked first on the three large, non-enumerable multi-site landscapes. In two successive rounds of savinase activity optimization, AlphaVariant-designed libraries yielded 75.4% and 83.8% activity-improved variants among experimentally characterized candidates even as the accessible mutation space expanded from 25 to 67 positions. The search identified beneficial mutation combinations involving up to 17 structurally dispersed sites and produced a variant with 7.1-fold higher activity than wild type after only two design-test cycles. We then extended AlphaVariant to jointly optimize activity, thermostability and a patent-informed constraint for intellectual-property freedom. This multi-objective search yielded variants that outperformed the commercial protease Uno 100 L in both biochemical assays and application-relevant detergent washing tests. These results demonstrate that generative RL can efficiently navigate vast protein fitness landscapes, progressively incorporate multiple engineering objectives and convert sparse experimental feedback into enzyme variants with industrially relevant performance, effectively resolving the limits of biocatalyst engineering.

## Introduction

Industrial enzymes are indispensable catalysts for sustainable manufacturing, underpinning sectors from pharmaceuticals and agriculture to food processing and material science. Their exquisite specificity and catalytic efficiency enable cleaner, lower-energy processes that reduce waste and diminish reliance on harsh chemical reagents. In practice, however, the performance demands of industrial applications — high catalytic activity, thermal resilience, detergent compatibility, and freedom from intellectual property (IP) conflicts — rarely converge in a single natural enzyme^1^. Meeting these demands requires engineering proteins across multiple residue positions simultaneously, because the functional improvements that matter most in industrial settings frequently arise from coordinated, higher-order combinations of substitutions rather than from isolated single-site changes^2,3^. Identifying such beneficial multi-site variants efficiently is therefore a central challenge for industrial biotechnology.

Traditional enzyme engineering strategies have long struggled with this challenge. Structure-based rational design is constrained by the scarcity of high-resolution structural data, the difficulty of predicting the effects of distal mutations, and pervasive epistatic interactions that render additive models unreliable, limiting its applicability to well-characterized systems. Directed evolution circumvents these structural requirements by mimicking natural selection through iterative mutagenesis and screening, but it faces a fundamental combinatorial barrier: the number of possible multi-site variants grows exponentially with the number of mutable positions, yet experimental throughput is capped by transformation efficiency and screening capacity. Conventional directed evolution therefore samples the protein landscape with extreme sparsity, and the resulting optimization cycles, resource-intensive as they are, suffer from an exceptionally low fraction of improved variants and are frequently trapped by rugged fitness landscapes^4,5^. The deeper the multi-site exploration required, the more acutely these limitations bite.

Machine learning has substantially advanced the field by learning sequence-fitness relationships from experimental data, enabling *in silico* ranking of candidate variants before costly synthesis and screening^6,7^. Active learning further improves data efficiency by iteratively selecting the informative variants for experimental testing, progressively refining the predictive model across rounds^8^. More recently, generative design methods have moved beyond ranking predefined libraries to proposing entirely new sequences^9,10^. Yet a persistent limitation remains: most current approaches struggle to design coordinated multi-site mutations and to accurately capture the epistatic interactions that govern high-performance regions of sequence space. Supervised predictors degrade when extrapolating beyond the measured sequence distribution, and generative models trained on natural protein sequences, while effective at producing foldable, biologically plausible variants, lack any mechanism to direct generation toward specific industrial objectives such as substrate activity, thermostability, or detergent compatibility. The evolutionary knowledge encoded in protein language models is rich, but it is agnostic to the task-specific reward signals that industrial engineering demands.

This convergence of challenges raises a fundamental question: can we formulate multi-site enzyme design as a feedback-driven, data-efficient process, one that simultaneously exploits the evolutionary plausibility encoded in generative protein priors and aggressively steers sequence generation toward specific, measurable industrial properties, even when labeled experimental data are scarce? Reinforcement learning (RL) offers a principled answer to this question. By framing sequence design as a sequential decision-making problem, RL enables an agent to propose variants and receive reward signals derived from predicted or experimentally measured properties, iteratively improving its policy without requiring large upfront datasets^11,12^. Crucially, when the RL agent is initialized from a generative protein prior and constrained to a user-defined mutational space, the search remains anchored to biologically plausible sequence neighborhoods while being actively directed toward high-reward regions that encode desired enzymatic properties^13,14^.

Here, we introduce AlphaVariant, a RL framework designed specifically for low-N multi-site enzyme optimization (**Figure 1**). AlphaVariant integrates three components: a dynamically defined mutation space that focuses the search on experimentally tractable positions; a VariantGPT generative prior trained on homologous sequences to bias generation toward evolutionary feasible variants; and an ensemble of task-specific fitness predictors that supply reward signals during policy optimization. Together, these components allow AlphaVariant to explore higher-order mutation combinations, including long-range epistatic interactions, that are inaccessible to single-site or low-order library designs, while remaining robust to the data scarcity that characterizes real industrial campaigns.

**Figure 1.**
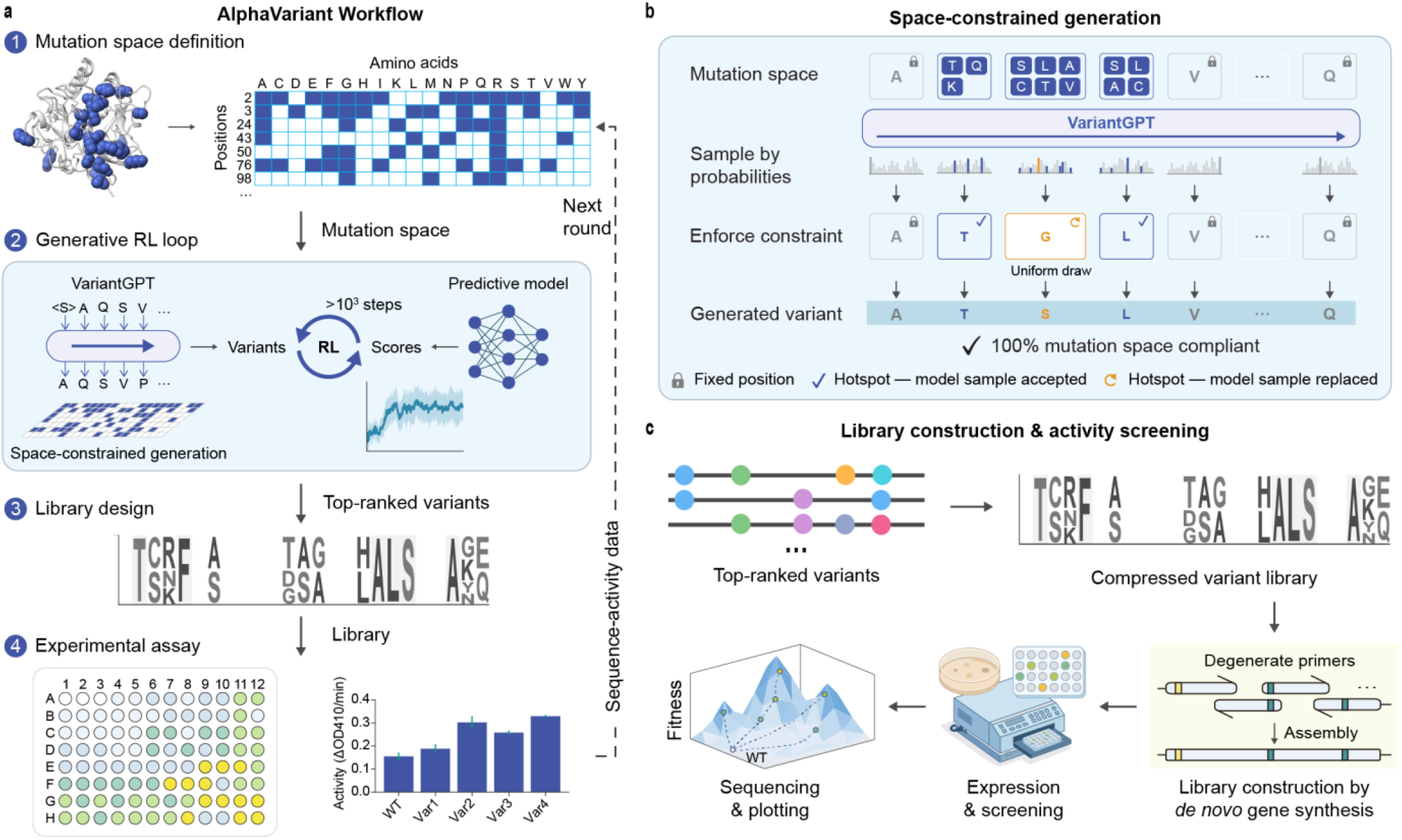
Overview of the AlphaVariant workflow for protein engineering. **a**, The four-step AlphaVariant workflow. First, the mutation space was defined based on structural insights and existing knowledge. Then, VariantGPT was trained through generative reinforcement learning (RL) loops to generate variants within the mutation space via space-constrained generation. The variants are scored by predictive models, and the generation policy was optimized over >10³ steps to generate variants of improved scores. After that, computational libraries were designed based on the top-ranked generated variants. Finally, experimental assays were conducted to obtain the activities of variants in the designed library. Sequence-activity data is fed back for the next round of optimization. **b**, Space-constrained generation mechanism. VariantGPT samples amino acids based on probabilities. The mutation space constraint was enforced to ensures that mutations only occur within the predefined mutation space: the fixed positions are locked, mutations in hotspots were allowed, while any amino acids sampled out of the predefined mutation space were replaced by uniform drawing from permitted amino acids in a site. This results in generated variants that are 100% compliant with the mutation space. **c**, Library construction and activity screening pipeline. Top-ranked variants are compressed into a variant library, which is then constructed via de novo gene synthesis using degenerate primers. The library is expressed and screened, and selected variants are assayed and sequenced to map the fitness landscape and identify improved variants.

We evaluated AlphaVariant in two complementary settings. First, we benchmarked the framework across six published protein fitness landscapes spanning diverse protein families and functional classes, comparing it against nine established sequence optimization methods. AlphaVariant performed competitively across all landscapes and ranked strongest on the three large multi-site landscapes, precisely the regime where the combinatorial space is too vast for enumeration-based methods. Second, we applied AlphaVariant to savinase, an industrial alkaline protease central to detergent formulations, where simultaneous improvements in activity, thermostability, and IP-conflict avoidance are required for commercial deployment^15^. Across two rounds of activity optimization, AlphaVariant-designed libraries achieved improvement rates of 75.4% and 83.8%, and the best second-round variant exhibited more than four-fold higher activity than wild-type savinase. In a third round of multi-objective optimization, three selected variants outperformed Uno 100 L, a leading commercial enzyme standard refined over decades of industrial development, in both laboratory assays and realistic detergent washing tests. These results demonstrate that reinforcement learning-guided sequence generation provides a powerful, data-efficient paradigm for navigating the complex, multi-objective fitness landscapes that define modern industrial enzyme engineering.

### Benchmarking AlphaVariant across protein fitness landscapes

A central question for any new sequence optimization method is whether its advantages are general or confined to specific problem settings. Existing methods span a wide range of strategies, from exhaustive supervised surrogates to generative and evolutionary search, and their relative performance is known to depend strongly on the size and topology of the fitness landscape. To characterize AlphaVariant’s performance across this spectrum, we benchmarked it against nine published methods on six experimentally measured fitness landscapes, revealing that AlphaVariant performs competitively on small, densely labeled landscapes and achieves the strongest overall ranking on large, non-enumerable multi-site landscapes, precisely the regime most relevant to industrial enzyme engineering.

To evaluate AlphaVariant against established sequence-optimization approaches, we assembled a benchmark of six experimentally measured fitness landscapes spanning diverse protein families and six functional classes (**Figure 2a**): a protein-binding protein (GB1^16^), a signaling kinase (PhoQ^17^), an RNA-binding protein (PAB1^18^), an enzyme (TrpB^19^), a fluorescent protein (CreiLOV^20^), and a viral-capsid assembly protein (AAV^21^). The landscapes form two complementary regimes designed to probe distinct optimization challenges. Three four-site combinatorial landscapes (GB1, PhoQ, and TrpB; 140,527–159,129 measured variants) are near-complete 20^4^ libraries in which every candidate has a measured fitness label, enabling exact lookup and providing a stringent test of a method’s ability to identify rare high-fitness combinations within a fully enumerable space. Three multi-site landscapes (AAV, CreiLOV, PAB1; 15–74 variable positions) define combinatorial spaces of approximately 10^19^ to >10^96^ sequences, far beyond any possibility of exhaustive enumeration, and were queried through learned surrogates of the landscapes^22^, testing a fundamentally different capability: the ability to navigate vast, non-enumerable sequence spaces where experimental coverage is necessarily sparse.

**Figure 2.**
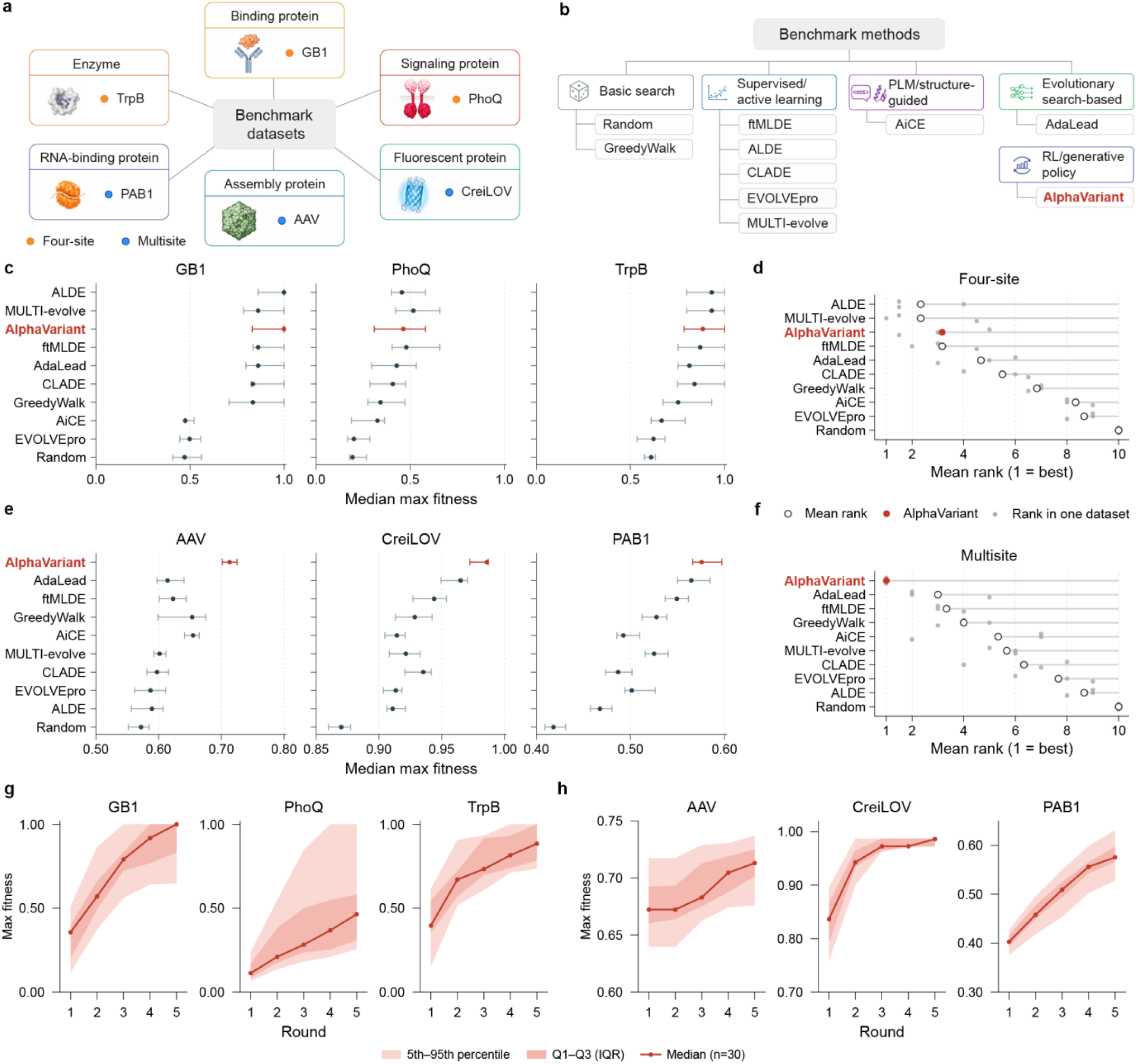
Benchmark of AlphaVariant against state-of-the-art computational methods. **a**, Diverse benchmark datasets encompassing six protein types: enzyme (TrpB), binding protein (GB1), signaling protein (PhoQ), RNA-binding protein (PAB1), assembly protein (AAV), and fluorescent protein (CreiLOV). Orange and blue dots indicate four-site and multi-site problems, respectively. **b**, Categorization of benchmarked methods, including basic search (Random, GreedyWalk), supervised learning or active learning methods (ftMLDE, ALDE, CLADE, EVOLVEpro and MULTI-evolve), protein language model (PLM) or structure-guided method (AiCE), evolutionary search-based method (AdaLead), and AlphaVariant as an RL/generative policy method. **c**, Comparison of median maximum fitness achieved by different methods on four-site datasets. AlphaVariant (highlighted in red) consistently achieves highly competitive performance. Dots show medians and error bars denote the Q1–Q3 interquartile range. **d**, Mean rank comparison across all benchmarked methods for four-site problems. Each grey dot represents the rank of a method on a single dataset; the open circle represents the mean rank across datasets; and the filled red dot highlights AlphaVariant’s mean rank. **e**, Comparison of median maximum fitness achieved by different methods on multi-site datasets. . AlphaVariant (highlighted in red) consistently achieves superior performance. **f**, Mean rank comparison across all benchmarked methods for multi-site problems. AlphaVariant achieves the best on these problems. **g**,**h**, Progression of maximum fitness over five optimization rounds for AlphaVariant on four-site (**g**) and multi-site (**h**) datasets. The solid red line represents the median of 30 independent runs, with the darker shaded region indicating the interquartile range (Q1–Q3) and the lighter shaded region indicating the 5th–95th percentile range.

We compared AlphaVariant with nine published methods spanning four families (**Figure 2b, Supplementary Note 1**): basic search (Random, GreedyWalk); supervised and active-learning methods (ftMLDE^23^, ALDE^8^, CLADE^24^, EVOLVEpro^25^, MULTI-evolve^26^); structure-guided method (AiCE^10^); and evolutionary search-based method (AdaLead^22^). All methods used an identical campaign budget of 480 queries (96 sequences per round × 5 rounds), were run for 30 independent seeds, and scored with two metrics: the normalized maximum fitness discovered, the primary target in directed evolution compaign, and the the mean fitness of the top 128 variants discovered (top-128 fitness), a batch-quality metric that rewards recovering many high-fitness variants rather than a single outlier.

We first benchmarked the methods on four-site landscapes. The four-site landscapes contain very few high-fitness variants: the fraction of sequences with normalized fitness ≥ 0.5 ranged from 0.007% (PhoQ) to 0.35% (TrpB) (**Extended Data Fig. 1a**). For PhoQ, a uniformly random campaign of 480 draws is not expected to recover any high-fitness variant (**Extended Data Fig. 1b**), so success requires guided exploration.

On maximum fitness (**Figure 2c**), AlphaVariant was competitive with the strongest baselines on all three landscapes. It reached the global optimum on GB1 (median maximum fitness 1.00, tied with ALDE) and obtained 0.83 on TrpB, and 0.46 on PhoQ. ALDE was the strongest four-site method overall (medians of 1.00 on GB1 and 0.93 on TrpB). Aggregating ranks across the three landscapes (**Figure 2d**), AlphaVariant placed third on maximum fitness, behind ALDE, and MULTI-evolve. On the batch-quality metric (top-128 mean fitness; **Extended Data Fig. 1c**), AlphaVariant was stronger (0.46 on GB1, 0.58 on TrpB and 0.12 on PhoQ), ranking second overall, and behind only ALDE (**Extended Data Fig. 1d**). In this regime, where exhaustive labels favour supervised surrogates, AlphaVariant performed comparably to the leading methods without any landscape-specific tuning (**Extended Data Fig. 1e,f**)..

The multi-site landscapes test optimization over sequence spaces too large to enumerate. We replaced the lookup table with a convolutional oracle trained on all measured variants of each protein^22^; the oracles were accurate on held-out data (test Spearman ρ = 0.89 on AAV, 0.98 on CreiLOV, and 0.90 on PAB1; n_test = 3,652–16,542; **Extended Data Fig. 2a,b**), and every queried sequence, including sequences distant from the measured set, was scored by the oracle.

In this regime, AlphaVariant achieved the highest median maximum fitness on all three landscapes, AAV (0.71), CreiLOV (0.99) and PAB1 (0.58) (**Figure 2e,f**). On top-128 fitness (**Extended Data Fig. 2c,d**), AlphaVariant was highest on AAV (0.65) and PAB1 (0.50) and close to the leaders on CreiLOV (0.90 versus 0.94 for AdaLead). Two-sided Wilcoxon signed-rank tests on paired per-seed values, Bonferroni-corrected for multiple comparisons (α = 0.05; **Extended Data Fig. 2e,f**), supported these differences: on AAV, AlphaVariant exceeded all nine baselines on both metrics (9/9 comparisons significant), and on PAB1 it was significant against 8/9 baselines for maximum fitness and 7/9 for top-128 fitness. Across the three multi-site landscapes, AlphaVariant had the lowest mean rank (**Figure 2f**; **Extended Data Fig. 2d**), ahead of AdaLead, ftMLDE, and GreedyWalk.

To identify which components underpin this multi-site performance, we ablated each in turn from the multi-site configuration (**Extended Data Table 5**; **Supplementary Note 2**). The dominant component was the mutation cap that restricts proposals to within two mutations of the reference: removing it degraded top-128 fitness increasingly with the size of the variable region, from a negligible change on CreiLOV (∼15 variable positions) to −0.25 on AAV (28 positions) and −0.39 on PAB1 (74 positions). The EV^27^-augmented surrogate features contributed on the larger, sparser landscapes (PAB1) but were approximately neutral on the smaller ones, whereas SHAP^28^-based alphabet pruning and the homolog-pretrained prior were approximately neutral throughout. These results indicate that constraining generation to the neighborhood of the measured data, where the learned oracle is reliable, is what makes policy-gradient search effective on non-enumerable landscapes, and that the value of this constraint grows with the size of the search space, precisely the regime in which AlphaVariant outperforms the baselines.

Across all six benchmark landscapes, the median maximum fitness rises monotonically from round one to five (**Figure 2g,h**). This sustained upward trajectory suggests that AlphaVariant has not yet reached the ceiling of its optimization capabilities within five rounds for PhoQ, TrpB, AAV and PAB1, and that continued iterative optimization would be expected to yield further improvements.

Considered together, these results demonstrate that AlphaVariant’s generative prior and policy-gradient optimization achieve performance comparable to the strongest supervised approaches on densely labeled four-site landscapes while ranking first among ten benchmarked methods on larger multi-site landscapes, using unified configurations without landscape-specific tuning. These findings suggest that AlphaVariant can effectively navigate diverse protein fitness landscapes and that its generative priors and policy-guided exploration become particularly advantageous as the search space expands.

However, benchmark datasets represent controlled experimental settings and do not fully capture the practical challenges of enzyme engineering, including limited assay data, library construction errors, expression variability, and application-specific optimization objectives. To assess whether these advantages translate to real-world protein engineering, we next evaluated AlphaVariant in a wet-lab optimization campaign targeting savinase, an industrial alkaline protease.

### AlphaVariant prioritizes multi-site savinase libraries from a constrained mutational space

Savinase (*Bacillus lentus* alkaline protease) is one of the most commercially significant industrial proteases, deployed at scale in laundry detergents worldwide for over three decades to remove proteinaceous stains under alkaline, low-temperature washing conditions. Despite this long history of industrial use, savinase remains a stringent and unsolved engineering target: its optimal performance demands the simultaneous improvement of catalytic activity, thermostability, and detergent compatibility, three properties that are often in tension and cannot be addressed by single-site substitutions alone. Critically, the dense patent landscape surrounding savinase and its close homologs means that any commercially viable engineering campaign must also guarantee IP freedom, adding a fourth, non-biochemical constraint that is entirely orthogonal to fitness optimization. The structural complexity of the active site and the pervasive long-range epistasis among functional residues further limit the utility of rational design, which has historically been restricted to well-characterized, near-active-site positions and has rarely achieved the coordinated multi-site improvements required for commercial deployment. Savinase therefore represents precisely the class of problem, multi-objective, multi-site, IP-constrained, and resistant to rational design, for which a data-efficient generative optimization framework is most needed.

Because publicly available activity data for savinase variants are scarce, we initialized the first round of design using experimentally characterized variants of subtilisin BPN′, the most extensively studied member of the subtilisin family and the canonical model system for serine protease engineering. Subtilisin BPN′ and savinase share approximately 60% sequence identity across the mature protease domain and adopt a nearly identical three-dimensional fold, with conserved catalytic triad geometry (Asp32–His64–Ser221 in BPN′ numbering) and a structurally equivalent substrate-binding cleft (**Extended Data Fig. 3**). This high structural homology means that mutation effects characterized in BPN′, including well-documented stabilizing substitutions, activity-modulating active-site changes, and epistatic interactions, provide a biologically grounded prior for savinase engineering. Mutation positions and amino acid substitutions reported for BPN′ were mapped to the savinase sequence by pairwise sequence alignment; positions with unambiguous one-to-one correspondence were retained for mutation space definition. This transfer step defined the initial constrained mutation space for the first round, after which experimentally measured savinase variants were used to update both the fitness models and the mutational space in subsequent rounds.

Our RL agent, guided by a specialized generative model (VariantGPT) and a predictive fitness model, was tasked with generating improved variants within a defined mutational space (**Figure 3a, Extended Data Fig. 4a,b**). The VariantGPT model, pretrained on a large corpus of homology sequences and fine-tuned on BPN’ variants, learned to generate focused, multi-site variants with a mean of seven mutations, a significant deviation from the high-mutation variants generated without fine-tuning (**Extended Data Fig. 4c**). This agent was guided by an ev+onehot fitness model^29^ that achieved a Spearman correlation of 0.72 on a held-out test set, providing reliable predictions of variant activity (**Extended Data Fig. 4d**).

**Figure 3.**
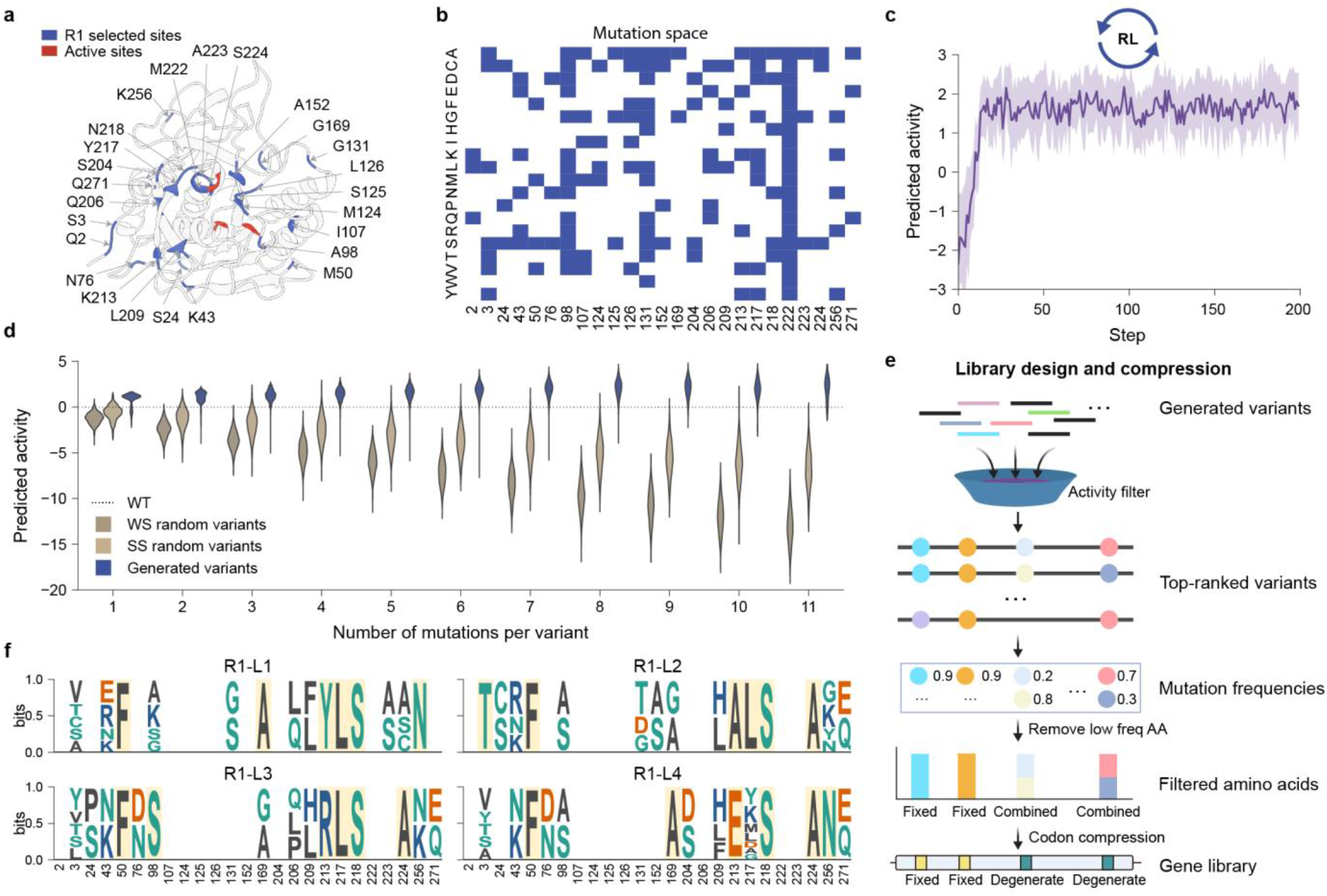
Round one savinase library design using AlphaVariant. **a**, The mutation sites selected in the first round (R1) of optimization. 25 sites were selected for mutagenesis, as highlighted in blue. The active sites are highlighted in red. **b**, The mutation space for R1. The allowed amino acid substitutions at each selected position across the full amino acid alphabet are predefined before the RL optimization. **c**, Optimization of predicted activity during the RL optimization process, showing rapid convergence toward high-fitness sequences within ∼50 steps. **d**, Comparison of the predicted activities between the AlphaVariant-generated variants and the random variants. Two types of random variants were analyzed, including variants with random mutations in the whole sequence (WS) and in the selected sites (SS). Generated variants are consistently enriched for high-predicted-activity sequences relative to random baselines. **e**, Workflow used to design computational libraries from the generated variants. **f**, The four libraries in the first round.

The RL optimization achieved a remarkable breakthrough in multi-site activity optimization. Within just 30 steps, the agent generated novel variants with enhanced predicted activities (**Figure 3c**) and multiple mutations (**Extended Data Fig. 4e,f**). A total of 42,117 unique improving variants were generated containing one to eleven coordinated mutations during the RL process. Critically, the agent learned to identify synergistic mutations, leading to a reversal of the typical relationship between mutation number and activity. While randomly generated variants tend to lose activity with an increasing number of mutations, the variants generated by AlphaVariant showed a positive correlation between mutation count and predicted activity (**Figure 3d**), which indicates our model’s ability to effectively explore the combinatorial effects of individual mutations. This *in silico* result provides strong evidence that our RL framework can effectively navigate the complexities of multi-site optimization and identify beneficial high-order mutation combinations, a capability that has been a long-standing challenge for protein design.

On the basis of the strength of these computational results, we proceeded with experimental validation. We designed four libraries of variants on the basis of computationally generated sequences, balancing predicted activity with sequence diversity to maximize the potential for discovering highly active enzymes during experimental testing (**Figure 3e,f**).

### First-round experimental screening validates activity-enriched savinase libraries

Having established that AlphaVariant can efficiently navigate multi-site fitness landscapes *in silico*, the critical test was whether these computational predictions would translate into experimentally improved enzyme variants. Conventional directed evolution approaches such as error-prone PCR are inherently low-yield: deleterious mutations dominate random libraries (30–50% of variants), while truly beneficial mutations are exceedingly rare^30^. Beyond this general challenge, a question specific to our campaign was whether the transfer learning strategy, initializing the mutation space and fitness predictor from BPN′ data and mapping it to savinase, would retain sufficient predictive fidelity to enrich improved variants in the wet lab.To address both, the four AlphaVariant-designed libraries were cloned into the expression plasmid pWB980, which incorporates the sacB signal peptide and prodomain upstream of the mature protease sequence to enable constitutive extracellular secretion in *Bacillus subtilis* (*B. subtilis)* WB600 under the p43 promoter (**Figure 4a**). This secretion-based format allows direct activity readout from culture supernatants, enabling high-throughput screening at scale. DNA libraries were constructed by amplifying five overlapping gene fragments with degenerate primers and assembling them by overlap-extension PCR, a strategy that introduces multiple mutations in a single synthesis step while preserving the reading frame (**Extended Data Fig. 11**).

**Figure 4.**
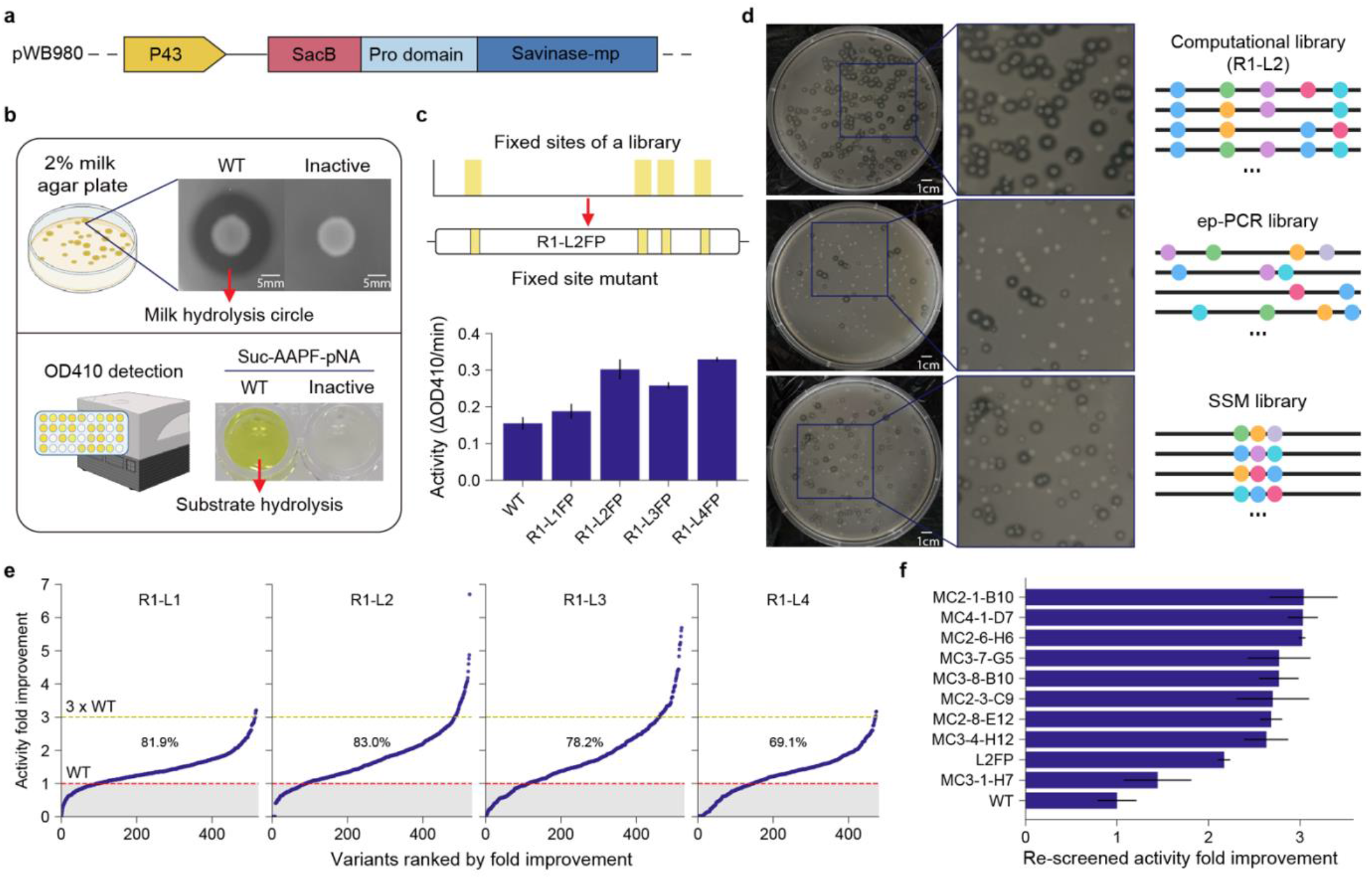
AlphaVariant-designed libraries achieve high experimental activity-improvement rates in savinase. **a**, Illustration of the plasmid used to express the variants. **b**, Examples of 2% milk agar plates and OD 410 used to select and determine the variant activities. On agar plates, active variants, such as the wild type (WT) displayed here, are characterized by colonies surrounded by clear hydrolysis halos, whereas inactive variants do not exhibit this clear halo. During OD410 assays, active variants catalyze the cleavage of suc-AAPF-pNA, generating a colored product with a distinct absorption at 410 nm, whereas inactive variants show no color change. **c**, The catalytic activities of the fixed mutants in the four libraries and WT. **d**, Comparison of the computational library (R1-L2) with the error-prone PCR (ep-PCR) and site saturation mutagenesis (SSM) libraries. **e**, The catalytic activities of the variants in the four constructed libraries (R1-L1 to R1-L4). The success rates of obtaining variants with activities better than that of WT are shown on the curves. **f**, Re-screened activity fold improvements of the top-performing variants from round one. Each variant was repeatedly assayed three times.

Screening proceeded in two stages. Active variants were first identified by the formation of clear milk-hydrolysis halos on 2% agar plates, a rapid, colony-level selection that eliminates inactive clones before quantitative assay (**Figure 4b**). Variants passing this selection were then expressed in liquid culture and their catalytic activity quantified using the chromogenic substrate suc-AAPF-pNA, which produces a colored product absorbing at 410 nm. As an early validation of the computational design, all four fixed-mutation variants derived from the computational libraries showed improved activity relative to wild-type savinase (**Figure 4c**). providing initial evidence that the RL agent had learned to identify beneficial multi-site mutation combinations prior to any library-level screening.

To benchmark the enrichment achieved by AlphaVariant, we ran parallel controls using two conventional library generation strategies, error-prone PCR and site-saturation mutagenesis at positions 211, 212, and 216. AlphaVariant-designed libraries produced substantially higher fractions of active variants on selection plates than either control (**Figure 4d**). A small fraction of library clones were inactive. Sequencing revealed that the majority of these failures arose from frameshift mutations, insertions and deletions, introduced during PCR assembly rather than from incorrect computational predictions, highlighting library synthesis fidelity as the primary experimental challenge.

Among the 2,406 variants that passed the two-stage screening workflow, 1,813 showed activity exceeding wild-type savinase, corresponding to an improvement rate of 75.4% (**Figure 4e**). Each of the four libraries contributed variants with more than three-fold activity improvements, and the top-performing variants after rescreening are shown in **Figure 4f**. Sequence analysis of the highest-performing variants revealed coordinated mutations spanning more than seven sites distributed across the protein structure, confirming that the RL agent had learned to identify synergistic mutation networks involving long-range epistatic interactions, precisely the class of multi-site combinations that conventional single-site or low-order library designs cannot access. Notably, 20–30% of the sequenced library clones carried mutations introduced outside the original computational designs during PCR assembly (**Extended Data Fig. 13**), effectively expanding the explored sequence space and providing an additional source of sequence– activity information for subsequent rounds.

Taken together, the first-round results established two key points. First, the transfer learning strategy, initializing the mutation space and fitness predictor from BPN′ data and mapping it to savinase, was experimentally validated: the 75.4% improvement rate demonstrates that evolutionary knowledge from a homologous system can be effectively leveraged to enrich beneficial variants in a data-scarce industrial target. Second, the first-round sequence–activity dataset, comprising 2,406 experimentally characterized savinase variants spanning up to 11 coordinated mutations, constituted a substantially richer training resource than the BPN′-derived initialization data. This raised a direct question: could AlphaVariant exploit this first-round experimental feedback to further improve its search efficiency and identify variants with even greater activity gains in a second round?

### Experimental feedback improves second-round activity optimization

The first round generated not only 1,813 improved savinase variants but also a sequence– activity dataset of 2,406 experimentally characterized multi-site variants, a resource far richer than the BPN′-derived initialization data used to launch the campaign. The second round was designed to test whether feeding this experimental feedback back into AlphaVariant could simultaneously expand the accessible mutation space and further increase the fraction of improved variants, directly probing the iterative learning hypothesis at the heart of the framework.

Analysis of the first-round sequence–activity data using statistical methods identified 42 additional mutation sites that contributed positively to enzyme performance (**Figure 5a,b, Extended Data Fig. 14a**). These sites were added to the initial 25-site space, with new mutations from R1 and five most frequent amino acids from homologous sequences incorporated at each new position. The resulting second-round mutation space spanned 67 sites and a combinatorial sequence space of 1.71 × 10^50^, an expansion of nearly 33 orders of magnitude relative to the first round, while remaining anchored to experimentally validated beneficial regions (**Figure 5a**).

**Figure 5.**
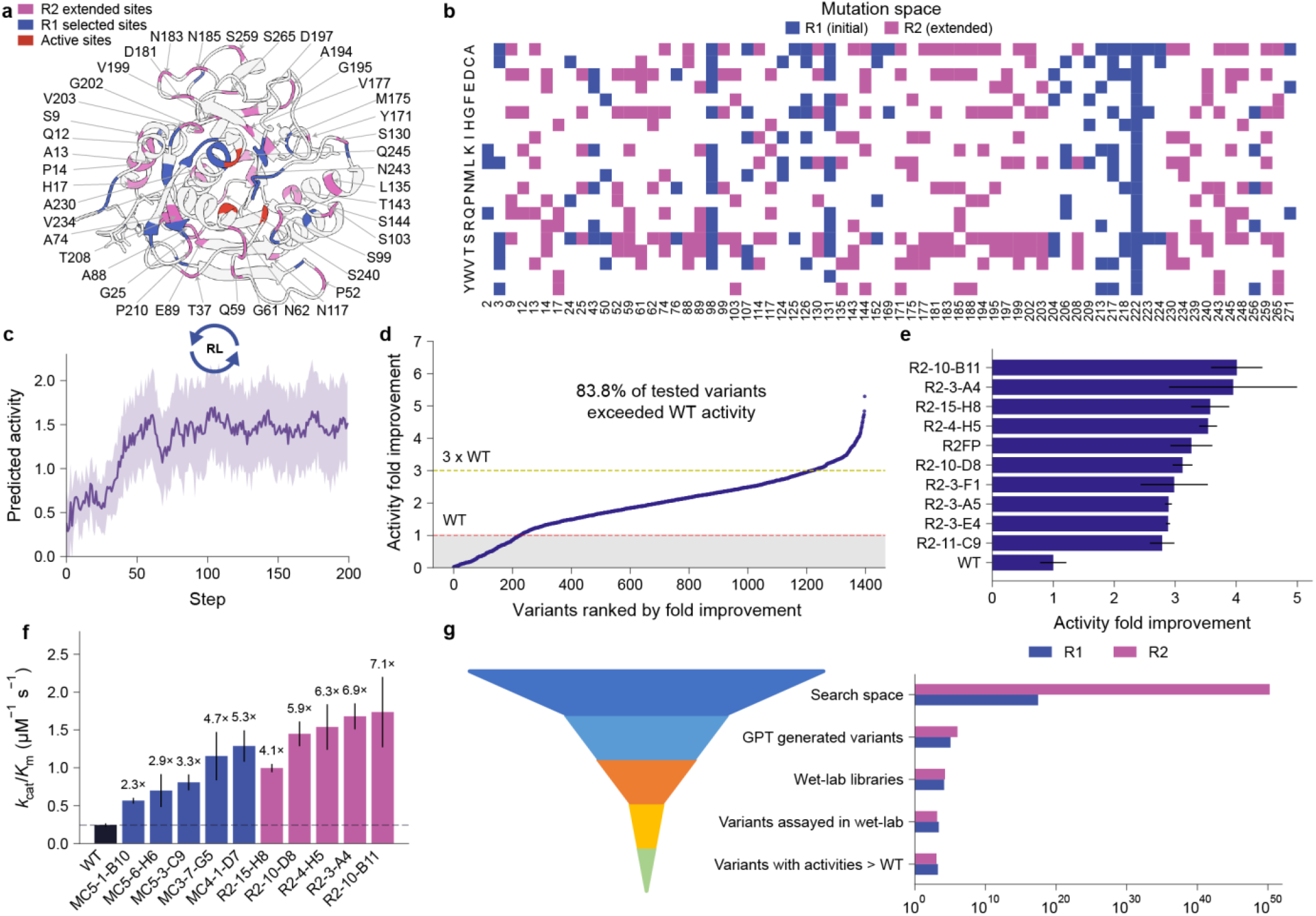
Experimental feedback improves AlphaVariant’s round two search efficiency and activity gains. **a**, The mutation sites selected in round two (R2). 42 sites were extended for mutagenesis in round two (purple). The active sites (red) and R1 sites (blue) are also highlighted. **b**, Expanded mutation space in R2. **c**, Optimization of activity in the RL process. **d**, Experimental catalytic activities of the variants in the R2 library. A total of 83.8% of the variants presented better activities than wild-type savinase. **e**, Activity improvements of the R2 top-ranked variants. Each variant was repeated three times. **f**, Kinetic parameters of wild-type savinase and the top five mutants from the two rounds. Each variant was assayed with three replicates. The best variant R2-10-B11 achieved a 7.1-fold improvement. **g**, The funnel of variant spaces explored in the first two rounds of experiments.

The fitness predictor was retrained on the first-round experimental data. We benchmarked multiple protein language model embeddings (ESM2-3B^31^, ProT5-XL^32^, and ProtAlbert^32^) alongside the proven ev+onehot^29^ approach and selected the ev+onehot model, which achieved the best Spearman correlation of 0.60 on a held-out test set (**Extended Data Fig. 14c,d**), a modest but expected decrease from the first-round correlation of 0.72, reflecting the greater diversity of the expanded sequence space. To explore multiple high-performance regions simultaneously, we initialized six parallel RL optimization runs from the six best first-round variants rather than from the wild-type sequence. This multi-start strategy allowed the agent to exploit experimentally validated beneficial starting points while independently exploring the expanded landscape, with each run converging to improved solutions within 60 steps (**Figure 5c**).

The computational outputs from the six parallel runs were consolidated into a single experimental library using the DGC-libgen method, which identifies shared amino acid preferences across optimization runs and encodes them using degenerate codons, enabling a compact library that captures the diversity of multiple RL trajectories without sacrificing coverage of high-predicted-fitness variants (**Extended Data Fig. 14e**, **Extended Data Fig. 16**). Experimental screening of approximately 1,500 colonies from the second-round library yielded an improvement rate of 83.8%, an increase of 8.4 percentage points over the first round (**Figure 5d**). The top-performing variant, R2-10-B11, exhibited more than four-fold higher activity than wild-type savinase (**Figure 5e**). Kinetic characterization of the top five variants from both rounds confirmed consistent improvements in k_cat_/K_M_ ratios, with the activity gains attributable primarily to increases in k_cat_ rather than decreases in K_M_, suggesting that the RL agent had learned to optimize the chemical step of catalysis rather than substrate binding affinity (**Figure 5f, Extended Data Fig. 14f,g**).

A direct comparison of the two rounds illustrates the compounding benefit of iterative experimental feedback (**Figure 5g, Extended Data Fig. 14h**): the second round achieved a higher improvement rate while exploring a sequence space nearly 33 orders of magnitude larger, and the top variant’s activity gain exceeded that of the best first-round variant by more than 30%. Together, the two rounds produced a combined sequence–activity landscape of over 3,900 experimentally characterized savinase variants spanning up to 17 coordinated mutations, providing a detailed map of the sequence–function relationships in this enzyme system (**Extended Data Fig. 19**).

However, high catalytic activity alone is insufficient for commercial deployment. Industrial detergent applications additionally require thermostability under wash conditions and freedom from IP conflicts, two objectives that the activity-focused RL reward function used in rounds one and two did not address. The third round therefore extended AlphaVariant to a fully multi-objective setting, asking whether the framework could simultaneously optimize activity, thermostability, and IP freedom to produce variants suitable for direct industrial application.

### Multi-objective optimization balances activity, thermostability, and IP-conflict avoidance

To address these industrial constraints, we extended AlphaVariant’s reward function from a single activity objective to a composite score that jointly reward predicted activity, thermostability, and IP-conflict avoidance (**Figure 6a**). Three corresponding predictive models were required: one for activity, one for thermostability, and one for IP-conflict avoidance. We systematically benchmarked multiple protein language model embeddings, ProtT5-XL^32^, ProtAlbert^32^, ESM-2^31^, and ESM-1v^33^, alongside the ev+onehot^29^ approach used in earlier rounds to identify the best-performing model for each property (**Extended Data Fig. 20c,d,**). ProtT5-XL embeddings achieved the highest accuracy for both activity and thermostability predictions and were selected for the RL process (**Extended Data Fig. 20f, g**).

**Figure 6.**
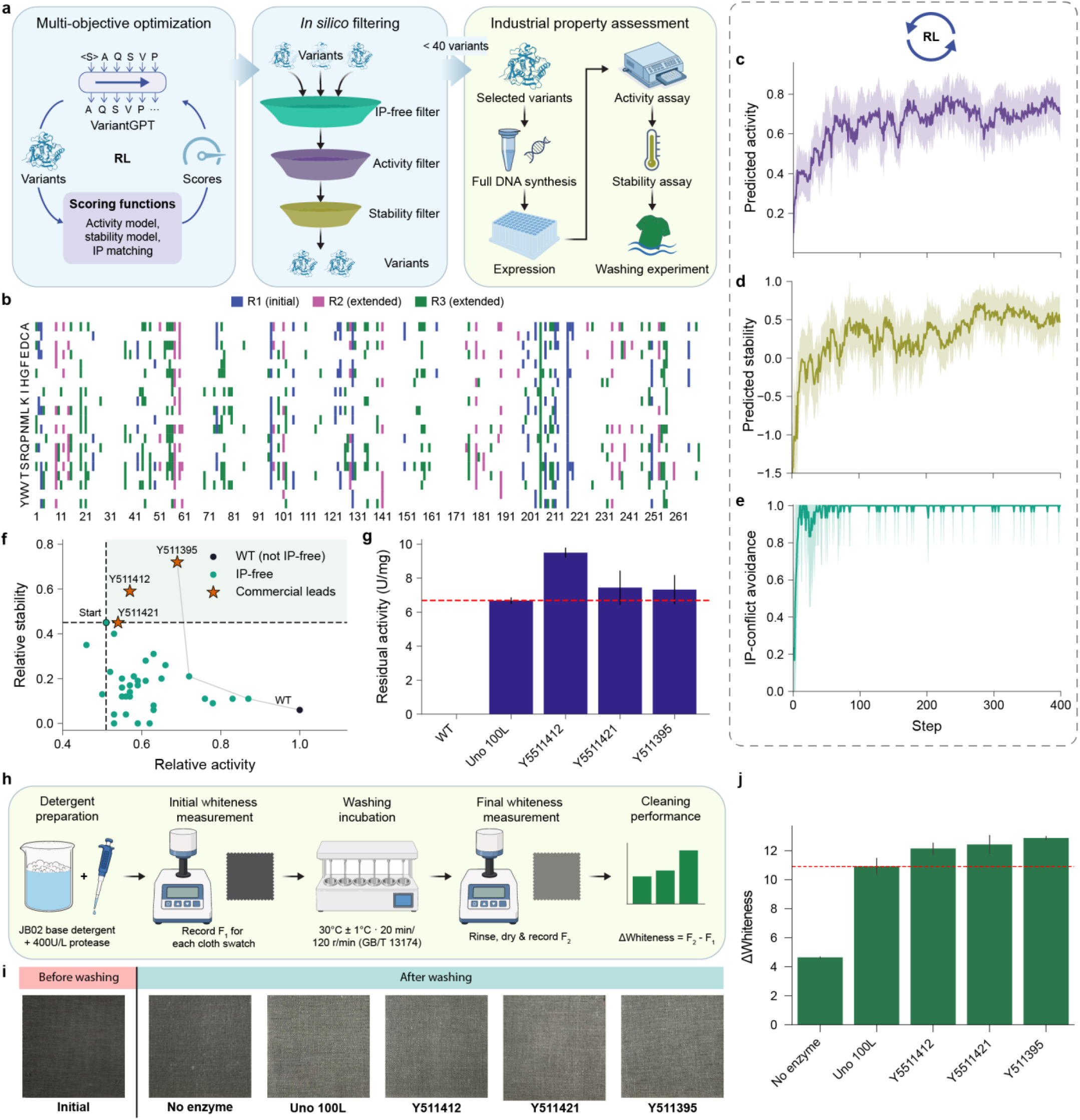
AlphaVariant identifies industrially relevant savinase variants through multi-objective reinforcement learning. **a**, Multi-objective optimization workflow for industrial application in the third round (R3). In this round, activity, stability, and intellectual property (IP) conflict avoidance were simultaneously co-optimized. Multiple predictors were added as feedback to the agent in the RL process. The generated variants were filtered via IP-free, activity, and stability filters to obtain 32 variants for extensive industrial tests. **b**, The mutation space in the third round. **c**-**e**, Improvements in activity (**c**), stability (**d**) and IP-conflict avoidance (**e**) properties in the RL process. **f**, Scatter plot of relative stability versus relative activity for WT, generated IP-free variants (green dots), and selected commercial leads (orange stars). The three top candidates (Y511412, Y511421, Y511395) are labeled, all exceeding the start variant in both activity and stability. **g**, Residual activity of WT, commercial benchmark Uno 100L, and the top three selected variants after thermo-incubation. **h**, Schematic of the washing experiment workflow to assess cleaning performance. The detergent containing the variants were incubated during washing (30°C ± 1°C, 20 min, 120 r/min, GB/T 13174). The change of whiteness of JB-02 soiled cloth was calculated for comparison by measuring the whiteness of the cloth before and after washing. **i**, Representative images of cloth swatches before washing and after washing with no enzyme, Uno 100L, and three variants. **j**, Quantitative whiteness score changes for all treatment groups. The three variants outperform both the no-enzyme control and the commercial benchmark Uno 100 L (dashed red line), confirming their superior cleaning performance.

The mutation space was expanded to 116 sites by incorporating high-perplexity positions, residues where the protein language model assigns low confidence to the wild-type amino acid, indicating evolutionary tolerance for substitution, alongside frequently observed amino acids from homologous sequences (**Figure 6b, Extended Data Fig. 20a,b**). This results in a R3 sequence space of 8.8×10^77^, more than 27 orders of magnitude larger than R2.

The key methodological innovation of this round was the integration of IP-conflict scoring directly into the RL reward signal (**Supplementary Note 3**). We developed an IP conflict detection method that quantifies potential patent overlaps for each generated variant and assigns a continuous IP-freedom score, rewarding the agent for proposing sequences that are both high-performing and commercially deployable.

Within R3 expanded search space, the RL agent simultaneously improved predicted activity, thermostability, and IP freedom within the first 100 steps, demonstrating its ability to navigate the competing trade-offs among all three industrial objectives (**Figure 6c–e**). The optimization process generated 513,048 variants, of which 235,098 passed the IP-freedom filter. Ensemble scoring with the top three activity and thermostability predictors identified the 200 most promising candidates, which were further refined through molecular dynamics simulations under the high-pH, high-temperature conditions. This multi-stage funnel reduced the candidate pool to 32 variants selected for experimental validation.

The 32 selected variants were expressed in *B. subtilis*, and their activities were quantified using casein as substrate to reflect industrial wash conditions more closely than the suc-AAPF-pNA assay used in earlier rounds. Thermostability was assessed by incubating variants at 85 °C for three minutes and measuring residual activities. Three of the 32 variants showed improved activities and stabilities in comparison to the start variant (**Figure 6f**). Further assays of the three variants together with Uno 100 L and WT showed that they outperformed Uno 100 L, a leading commercial enzyme standard that represents decades of industrial protein engineering, in both activity and thermostability assays (**Figure 6g, Extended Data Fig. 20h**).

To evaluate the performance of the three variants in real-world applications, we conducted washing tests following the standardized detergency test (JB02) (**Figure 6h**). Critically, washing performance tests under realistic detergent conditions confirmed that all three variants delivered superior cleaning efficacy compared with Uno 100 L (**Figure 6i,j**), demonstrating that the computational gains translated directly into application-relevant performance.

These results demonstrate that a single AlphaVariant campaign, requiring only 32 experimental measurements in the final round, was sufficient to identify IP-free savinase variants that exceed a commercial standard refined over decades of industrial development. The ability to simultaneously optimize three competing industrial objectives, overcoming trade-offs that typically constrain multi-trait optimization, represents a qualitative advance over conventional enzyme engineering workflows that optimize traits sequentially or in isolation.

## Discussion

The central question posed by this work was whether RL could bridge the longstanding gap between the evolutionary knowledge encoded in generative protein models and the task-specific reward signals demanded by industrial enzyme engineering, all under the severe constraint of scarce experimental data. The three-round savinase campaign provides a clear affirmative answer. Starting from a cold-start transfer of BPN′ data, AlphaVariant iteratively enriched experimentally improved multi-site variants across successive rounds, each time exploiting accumulated experimental feedback to expand the accessible sequence space and sharpen the fitness predictors. The culmination, three novel, IP-free savinase variants that outperformed Uno 100 L in both laboratory assays and realistic detergent washing tests, demonstrates that reinforcement-learning-guided sequence generation is not merely a computational curiosity but a practically deployable strategy for industrial enzyme engineering. A central and perhaps most striking finding of this work is the RL agent’s capacity to discover synergistic, long-range epistatic interactions that are inaccessible to conventional single-site or low-order library designs. The highest-performing R1 and R2 variants validated *in vitro* harbored between eight and 16 mutations distributed across structurally distal regions of the protein, yet these mutations acted cooperatively to produce activity gains far exceeding what any individual substitution could achieve. This observation reinforces a fundamental principle of protein function: the fitness landscape is not a smooth, additive surface but a rugged terrain in which peak performance emerges from specific networks of coordinated changes. Critically, AlphaVariant’s policy-gradient formulation, which optimizes the joint probability of entire sequences rather than scoring individual positions independently, is architecturally suited to discovering precisely these high-order dependencies, a capability that supervised predictors and single-site saturation libraries cannot replicate.

The benchmark results clarify when generative RL search adds the most value relative to discriminative approaches. On densely labeled, fully enumerable four-site landscapes, supervised and active-learning methods such as ALDE and ftMLDE performed exceptionally well, leveraging exhaustive fitness labels to build highly accurate surrogates. AlphaVariant was competitive in this regime but did not dominate it, a result that is mechanistically expected, because the primary advantage of generative search lies not in re-ranking a known finite library but in proposing novel sequences within a space that is too large to enumerate. On the three non-enumerable multi-site landscapes, where combinatorial spaces ranged from 10^19^ to >10^96^ sequences, AlphaVariant achieved the lowest mean rank across all ten benchmarked methods, with statistically significant advantages on AAV. This regime-dependent performance profile provides a principled guide for practitioners: when a near-complete fitness landscape is available, supervised surrogates remain the method of choice; when the search space is vast and data are sparse, the defining condition of real industrial enzyme campaigns, generative RL offers a decisive advantage.

The R3 optimization extended this advantage into a genuinely multi-objective industrial setting, simultaneously targeting catalytic activity, thermostability, and freedom from IP conflicts. The ability to encode patent-rule-based IP constraints directly as a reward term, and to have the RL agent navigate these constraints without sacrificing enzymatic performance, represents a qualitative expansion of what computational protein design can deliver. The three variants that outperformed Uno 100 L were not marginal improvements; they demonstrated superior performance in both controlled laboratory assays and realistic washing tests under detergent conditions. The fact that a computationally driven, data-efficient framework could identify variants superior to a product refined through decades of industrial enzyme engineering suggests that even highly optimized commercial enzymes occupy local optima in a fitness landscape that harbors substantially better solutions, solutions that require a global, multi-objective search strategy to reach.

Interpreting these results requires acknowledging the boundaries within which AlphaVariant operates. Three limitations are particularly important for future applications. First, like all RL methods that optimize against surrogate reward models, AlphaVariant is susceptible to reward hacking: the policy-gradient agent can learn to exploit artifacts in the fitness predictor rather than true biological fitness, generating sequences that score well *in silico* but fail experimentally^34^. In this work, the mutation cap and the constrained mutation space served as practical safeguards against this failure mode, but they do so by limiting the search radius rather than by principled uncertainty quantification. Second, the framework’s effectiveness depends on the availability of homologous sequence data and related experimental datasets to initialize the generative prior and the low-N surrogate. The successful transfer from subtilisin BPN′ to savinase demonstrates that this “warm start” strategy can be powerful, but it also defines the current scope of the method: applying AlphaVariant to entirely novel enzyme families without rich homologous data or existing mutation datasets remains an open challenge. A third, more practical limitation is that a fraction of library variants in each round were rendered inactive by frameshift mutations introduced during PCR assembly, highlighting that the fidelity of library synthesis remains a bottleneck that limits the full realization of computational designs. These limitations point directly toward the next generation of improvements. Replacing the hard mutation cap with principled uncertainty quantification, for example, through ensemble disagreement scores^8^ or trust-region policy optimization^35^, would allow the agent to distinguish between genuinely unexplored high-fitness regions and regions where the surrogate is simply overconfident, enabling more principled exploration–exploitation trade-offs. For mutation site identification, integrating structure-based approaches would move beyond the sequence-statistical signals currently used, enabling the framework to reason about steric compatibility and active-site geometry. Most ambitiously, structural generative models such as ESM3^36^ and ProteinMPNN^37^ could serve as secondary physical reward functions, penalizing mutations that cause severe steric clashes or destabilize the fold, acting as biophysical constraints that prevent the RL agent from exploiting artifacts in the sequence-based activity surrogate. Together, these advances would transform AlphaVariant from a framework that works well within a constrained neighborhood of known sequences into one capable of genuine *de novo* exploration.

In conclusion, this work establishes RL as a viable, data-efficient, and industrially deployable strategy for multi-objective enzyme optimization. The key insight is architectural: by formulating sequence design as a feedback-driven decision-making problem and anchoring the search in a generative evolutionary prior, AlphaVariant can navigate the vast, rugged fitness landscapes that define real industrial engineering challenges, landscapes that are simultaneously too large for exhaustive enumeration and too data-sparse for conventional supervised learning. The savinase campaign demonstrates that this approach can deliver variants competitive with the best commercially available enzymes within a small number of experimental rounds, without requiring the years of iterative directed evolution that traditional industrial development demands. As uncertainty-aware models and structural constraints are integrated into the RL objective, reinforcement-learning-guided protein design is poised to evolve from a promising research tool into a robust, general platform for the next generation of industrial biocatalysts.

## Methods

### The AlphaVariant framework

AlphaVariant couples a generative protein-sequence prior (VariantGPT) with reinforcement learning (RL) to propose high-value variants within a defined mutation space. The same framework underlies both the benchmark evaluation and the savinase campaigns; the components are described here, and application-specific instantiations (prior architecture, reward, and constraints) are given in the corresponding sections below.

#### Generative prior (VariantGPT)

The prior is an generative pre-trained transformer (GPT)^38^ model trained by next-token prediction on protein sequences, pretrained on a homologous family and optionally fine-tuned on task-specific variants. The agent used in RL is initialized from this prior; the prior remains fixed and serves as the anchor in the RL objective. The specific architecture and training corpus are given per application (4-layer GPT for the benchmark; the larger VariantGPT for savinase).

#### RL objective

Variants are optimized with an augmented-likelihood policy gradient adapted from REINVENT^39^. For a sampled sequence s = (s_1_, … , s_L_), the agent and prior log-likelihoods are summed autoregressive token log-probabilities:

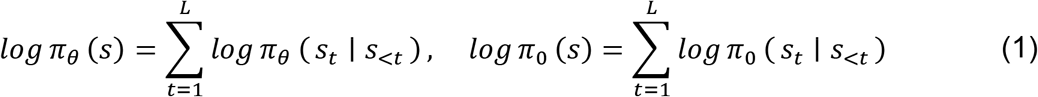

The augmented likelihood combines the prior with the reward:

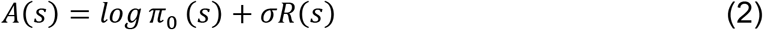

where π_0_ is the fixed prior model, *π_θ_* is the trainable agent, *R*(*s*) is a task-specific reward in a bounded range, a surrogate-ensemble upper-confidence bound for the benchmarks, and a transformed activity/multi-property score for the savinase campaigns, and reward scale *σ* controls the strength of reward-driven displacement from the prior. The agent is trained to minimize:

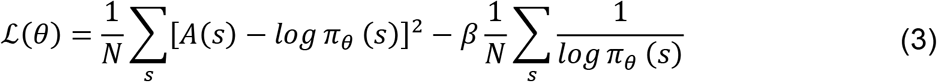

With regularization coefficient β = 5 × 10^3^ . The squared-deviation term increases the likelihood of high-reward sequences while anchoring the agent to the prior distribution; the second term is an exploration regularizer that discourages premature collapse of the agent distribution.

#### Space-constrained generation and library design

A space-constrained generation strategy is implemented to enforce the mutation space constraints. Generation is restricted to a predefined mutation space: the agent samples permitted amino acids at the designated mutable positions while wild-type residues are retained elsewhere. If a non-permitted amino acid is sampled, it will be replaced by uniform drawing from the permitted amino acids in a site. Optimization proceeds over iterative rounds in which the reward model is refit on accumulated data and the agent is re-optimized. Variants produced during RL are aggregated and prioritized to construct experimental libraries, as detailed per application.

### Benchmarking AlphaVariant on fitness landscape

#### Benchmark datasets

We evaluated all methods on six protein-fitness landscapes (**Figure 2a**). Assay values were min–max normalized to [0, 1] within each landscape, so the global maximum maps to 1.0 and metrics are comparable across proteins. The landscapes were grouped into two regimes.

Three four-site combinatorial landscapes were used as exact lookup tables: GB1 (IgG-binding B1 domain, positions 39/40/41/54; 149,361 variants^16^), PhoQ (signaling histidine kinase; 140,517 variants^17^), and TrpB (tryptophan synthase β-subunit; 159,129 variants^19^) (**Extended Data Table 1**) . Each is a near-complete 20^4^ library in which every candidate has a measured fitness. The landscapes are sparse in high-fitness variants: the fraction of sequences with normalized fitness ≥ 0.5 was 0.13% (GB1), 0.007% (PhoQ), and 0.35% (TrpB) (**Extended Data Fig. 1a**).

Three multi-site landscapes were used with a learned oracle: AAV (capsid VP1 28-residue diversification region^21^), CreiLOV (LOV-domain fluorescent protein, 119 aa, ∼15 variable positions^20^), and PAB1 (RNA-recognition motif, 75 aa^18^) (**Extended Data Table 2**). These span combinatorial spaces of approximately 10^36^ (AAV) to >10^96^ (PAB1) sequences. A lookup table returns zero for any unmeasured sequence, which collapses guided search; we therefore scored candidates with a trained oracle.

#### Learned oracle for multi-site landscapes

For each multi-site landscape we trained a sequence-to-fitness oracle using a length-agnostic convolutional architecture (1D convolution, 20→256 channels, kernel size 5; length-wise global max-pooling; linear regression head), matching the predictor used in AdaLead^22^. Oracles were trained on all measured variants by minimizing mean-squared error (Adam, learning rate 1×10^−4^, weight decay 1×10^−4^, batch size 1,024, ≤100 epochs; PyTorch 2.1.1) with fitness-weighted sampling (weight ∝ 1/(target − min + 1)) to emphasize the low-fitness range and avoid over-scoring and letting a method “win” by finding blind spots. The fitted min/max scaler was stored with each checkpoint and used to de-normalize predictions.

Oracles were accurate on held-out test variants (test Spearman ρ = 0.89/0.98/0.90 and R^2^ = 0.78/0.95/0.78 for AAV/CreiLOV/PAB1; n_test = 4,412/16,542/3,652; **Extended Data Fig. 2a**). During optimization we used a pure-oracle policy: every queried sequence, including sequences far from the measured set, was scored by the oracle, and measured labels were used only to train the oracle. To detect oracle-exploitation artifacts we recorded the mutational novelty of selected sequences relative to the measured set (**Extended Data Fig. 2b**).

#### Optimization campaign

All methods ran an identical campaign: an initial random batch of 96 sequences followed by four model-guided rounds of 96, for 480 queries in total (96 × 5), the budget commonly used in machine learning directed-evolution benchmarks^8^. On four-site landscapes a query was resolved by exact lookup; on multi-site landscapes it was scored by the trained oracle. For methods that select from a fixed enumerated library (ALDE, CLADE, and ftMLDE), the multi-site candidate source was replaced by a mutate-from-elites generator so that all methods operated over the same non-enumerable space while retaining each method’s own surrogate model and acquisition rule.

#### Compared methods

We benchmarked nine published methods alongside AlphaVariant, grouped by family (**Figure 2b**, **Extended Data Table 3**). Two basic search methods were tested, including Random (uniform sampling each round) and GreedyWalk (evaluation of single-mutation neighbors of the current best); five supervised and active-learning methods compose of ftMLDE^23^ (cross-validated ensemble of pretrained predictors), ALDE^8^ (ensemble regressor with Thompson sampling), CLADE^24^ (ensemble with cluster-stratified selection), EVOLVEpro^25^ (ESM-2 embeddings with a random-forest head; facebook/esm2_t12_35M_UR50D) and MULTI-evolve^26^ (bootstrap neural-network ensemble); one structure-guided design method, AiCE^10^, using ProteinMPNN structure-conditioned per-position amino-acid frequencies (1,000 samples, T = 0.5) blended with observed high-fitness frequencies; one evolutionary search-based method, AdaLead^22^ (recombination with surrogate-guided mutation), and our generative policy-based search methods, AlphaVariant. Each method was run in the configuration recommended by its authors and used the unified metric pipeline.

#### AlphaVariant settings

Within the benchmark, AlphaVariant followed the same framework, VariantGPT generator, augmented-likelihood reinforce^39^ objective (**Equation (3)**), surrogate-derived reward, and cluster-based selection. The generator was a four-layer GPT with four attention heads and embedding dimension 128, its context length matched to each sequence. The reward *R*(*s*) was the surrogate-ensemble upper-confidence bound (**Equation (4)**),

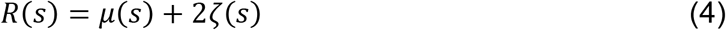

over a five-model ensemble (two ridge, one Bayesian ridge, one random-forest, one gradient-boosting regressor); surrogate features were per-position one-hot amino-acid encodings plus four physicochemical descriptors (side-chain volume, hydropathy, solvent-accessible area, and polarity), where μ(s) and ζ(s) denote the mean and standard deviation, respectively, of predictions from the five-model surrogate ensemble.

All runs used five rounds, batch size 96, 500 reinforce updates per round and *σ* = 60 (480 queries per campaign). Each campaign began with one cluster-based initialization round (k-means on sequence features, 96 sequences across 10 clusters, no surrogate/reward); in rounds 2–5 the surrogate was refit on all data, the agent optimized, a proposal pool sampled, and the next 96 sequences chosen by cluster-stratified selection (top-1,000 by predicted fitness, then best-per-cluster).

The two regimes differed only in landscape source and a few components (**Extended Data Table 4**). Four-site configs are: exact lookup-table fitness; an in-run GPT prior finetuned on the collected sequences in each round (10 epochs, lr 1 × 10^4^); no reward shaping, alphabet pruning, or mutation cap. While the multi-site configs are: learned CNN-oracle fitness; a homolog-pretrained GPT prior (no per-round finetuning); surrogate features augmented with a standardized EVmutation^27^ statistical-energy score; SHAP^28^-based alphabet pruning (enabled at ≥ 50 samples; SHAP > 0 retained; min 3, ma× 10 amino acids per position) propagated into the generator with violating proposals filtered; and a cap of two mutations from the reference variant.

#### Evaluation metrics

We report two complementary metrics per campaign. Maximum fitness is the highest normalized fitness among the 480 queried variants, the canonical directed-evolution objective. Top-128 fitness is the mean fitness of the 128 highest-fitness discovered variants, a batch-quality metric that rewards recovering many strong variants. Method rankings (**Figure 2c–f**) were computed per landscape (rank 1 = highest median for max fitness or highest mean for top-128 fitness; ties assigned the average rank) and averaged within each regime.

#### Statistics and reproducibility

Each method × landscape combination was run for n = 30 independent random seeds (the first 30 entries of a fixed seed list). Distributions across seeds are heavy-tailed and multimodal; we therefore summarized each combination by its median and interquartile range (Q1–Q3) rather than by the mean (**Figure 2c,e**; **Extended Data Fig. 1c, Extended Data Fig. 2c**). Pairwise differences between methods were assessed with two-sided Wilcoxon signed-rank tests on the paired per-seed values, with Bonferroni correction for multiple comparisons at α = 0.05 (**Extended Data Fig. 1e,f, Extended Data Fig. 2e,f**). No data were excluded and seeds were fixed for reproducibility.

### Implementation of AlphaVariant on savinase optimization

In the savinase optimization project, the VariantGPT prior model was trained on homologous protein sequences with six million parameters. The pretraining was to establish a strong evolutionary prior, followed by fine-tuning on experimentally characterized enzyme variants to adapt the model for specific optimization objectives. The per-round datasets and mutation space configurations can be found in **Extended Data Table 6** and **Extended Data Table 7**.

#### The first round

##### Datasets

The datasets used in the first round include homology sequences, publication and patent data. The homologous sequences of subtilisin BPN’ were collected from the Pfam subtilase family sequences (Pfam id: PF00082)^40^. The variant activity data were manually collected from publications to train an activity model. Only variant data with suc-AAPF-pNA as the substrate and subtilisin BPN’ as the wild type were selected to build the model, which resulted in 150 records. The patent data were also manually collected. These patent and publication datasets were used to define the mutation space.

##### Mutation space

The subtilisin BPN’ sequence consists of 275 positions, resulting in a vast mutation space to explore. To efficiently generate active variants, we utilized existing knowledge from the literature and patents to narrow down the search space by selecting specific positions for optimization, which we refer to as mutation sites (**Extended Data Fig. 4a**). We identified 25 mutation sites on the basis of the positions reported to be present in variants with improved activity. Within each mutation site, both the wild-type amino acid and amino acids known to have improving effects were permitted. This narrows the sequence space to explore to a size of 2.80 × 10^17^.

##### Predictive model

A low-N fitness model was built to predict the activity of subtilisin BPN’ variants following the protocol of Hsu *et al*.^29^. The activity data of subtilisin BPN’ variants with suc-AAPF-pNA as the substrate at room temperature (20 to 25°C) were obtained from publications. Variants with deletions or insertions were removed, resulting in 150 sequence-activity paired data. The activity scores were normalized by dividing them by the wild-type subtilisin BPN’ activity and then transformed to a logarithmic scale. The ev+onehot model was used to build the activity model, with 80% of the variants used for training and 20% (30 variants) used for testing.

##### Prior model

For savinase, the prior was VariantGPT, a GPT-2-style autoregressive model (∼6 million parameters; input and positional embeddings, eight decoder blocks with masked multi-head self-attention and feed-forward layers, residual connections with layer normalization, and embedding dimension of 256). It was pretrained on the Pfam subtilase family (PF00082^40^; 56,623 sequences of length 127–478, 90/10 train/test, 20 epochs, batch size 128) and fine-tuned on 150 published Subtilisin BPN’ variants (10 epochs), with cross-entropy loss and AdamW (1 × 10^−4^).

##### RL optimization

The agent (initialized from VariantGPT) was optimized with the augmented-likelihood objective (**Equation (3)**), constrained to permitted amino acids at the mutation sites with wild-type residues elsewhere. The reward *R*(*s*) was the predicted activity transformed to [0,1] by a sigmoid with lower/upper bounds *l* = −1, ℎ = 2 (**Equation** (5)).

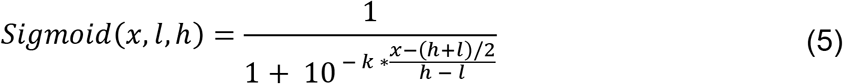

##### Library design

To design libraries for wet-lab experiments, we enumerated high-frequency mutation combinations (MC) within activity-improving variants (**Extended Data Fig. 6b**). An MC was defined as a set of mutations that were present in at least 10 improving variants. We iteratively examined the generated improving variants to identify all possible MCs. First, 36,406 MCs with one to eight mutations among the improving variants were obtained via a dynamic programming algorithm. The mutation sites for each MC were subsequently identified by enumerating the positions other than the fixed sites. If an MC involved more than ten mutation sites, the top ten mutation sites with the highest occurrence rates were retained. Finally, the permitted amino acids for each position other than the fixed position were determined by analyzing the covered variants and calculating their frequencies.

Once MCs were obtained, we ranked them to identify those with the highest potential to cover variants with high activities. A weighted score for each MC was calculated by ensembling the activity of the variant with fixed mutations, the mean activity of the covered variants, and the maximum activity of the covered variants. The scores were scaled from zero to one via the empirical cumulative distribution, with the average calculated value being the weighted score. The top 10 MCs were selected to construct preliminary libraries. To reduce cost, we merged these top 10 MCs into four libraries by expanding MC-1, MC-2, MC-7, and MC-9 to cover the remaining MCs while introducing as few mutagenesis sites as possible. The final four expanded libraries were used as the libraries for wet-lab experiments (**Figure 3f**).

#### The second round

##### Datasets

Extending from the first round, the homologous sequences of savinase were retrieved, and R1 sequence-activity datasets were utilized in the second round. The homologous sequences of savinase were obtained by querying with the wild-type sequences in the Uniref100 database via JackHmmer ^41,42^. 2,414 sequences were collected from the missense variants in the first round.

##### Mutational space

We extended the mutation sites by including new improving mutations and positions in the previous round. Statistical analysis and SHAP^28^ analysis were performed to obtain mutations with positive contributions in both analyses. Additionally, the mean activity of variants that contained a mutation was calculated, and mutations with mean activities lower than those of wild-type savinase were excluded. This resulted in 42 new positions, and the amino acids in these positions were extended to include the top five most frequent amino acids in savinase homology sequences. The new positions were merged with the sites in the previous round, resulting in 67 mutation sites covering a sequence space of 1.71 × 10^50^.

##### Predictive model

We evaluated nine protein language models (pLMs) alongside the ev+onehot model on the first-round activity data. For each pLM, PCA preprocessing of the embeddings followed by a support vector regression head gave the best performance; the data were split into train/test sets across 20 seeds and models ranked by mean Spearman correlation. Ev+onehot, ESM-3B, ProtT5-XL and ProtAlbert performed best; ev+onehot achieved the highest correlation (Spearman 0.60; **Extended Data Fig. 14c,d**) and, being fast at inference, was used as the predictive model in the RL process.

##### Prior model

Unlike the first round, which used subtilisin BPN′ as the template, the second round used savinase as the template. A VariantGPT prior was pretrained on savinase homologs, retrieved from UniRef100 with JackHmmer^41,42^ (80/20 train/test, 20 epochs), and fine-tuned on the 2,414 R1 variants; the fine-tuned model served as the prior in the RL process. **RL optimization.** The RL process of the second round is similar to that of the first round. One main difference is that the top six variants with the best catalytic activities in the first round were selected as the starting sequences to train the agents. That is, we launched six RL tasks with the top six variants in the first round as the starting variant. Only mutations within the mutation sites were allowed for the agent model. The ev+onehot model was used as the activity model. Each agent model was trained for 3,000 steps with a batch size of 30. The variants generated through the RL process were collected for downstream analysis.

##### Library design

For each run, the top 10,000 variants by ev+onehot score were re-scored by three pLMs (ESM-2b, ProtT5-XL, ProtAlbert) and ensemble-ranked by mean rank. Libraries were then generated with DGC-libgen: among the top variants, a position was fixed when one amino acid exceeded 0.75 frequency, amino acids below 0.10 frequency were dropped, and DYNAMCC0^43^ converted the retained amino acids at each position into degenerate codons; a sequence logo was produced per library. Because the six per-run libraries were highly similar, they were merged, pooling unique positions and permitted amino acids and recomputing degenerate codons with DYNAMCC0, into a single library of size 17,280 (**Extended Data Fig. 14e, Extended Data Fig. 16**).

##### New mutation identification

We conducted an analysis to obtain new mutations that contribute to the improvement of activities in savinase variants. The mutations that occurred in the variants whose activity improved by more than 2.5-fold were obtained. Mutations that were reported in publications and patents were removed. This resulted in 41 mutations that occurred at 32 positions, with 14 positions presented at least twice among the experimentally improved variants (**Extended Data Fig. 14i,j**).

#### The third round

##### Datasets

Extending from the second round, we collected in-house activity and stability datasets and patent claim data in the third round. In this round, casein was used as the substrate. A sequence-activity variant dataset of 459 variants was collected from previous works where casein was used as the substrate. Among them, 358 variants were assayed for stability.

The matching rules from 71 unique patents were retrieved. Their claims were extracted to define a patent rule to identify intellectual property (IP) conflict variants (as detailed in **Supplementary Note 3**). The rules were manually defined for each patent and verified by an IP expert.

##### Mutation space

We extended the mutation sites by including new positions in the third round. All mutations that were explored in the in-house activity and stability datasets were included in the mutation sites. Mutations (21A, 56T, 82A, 129T, 195S, 251Q, 256 N, and 271Y) and positions (140, 165, 251, and 257) that showed negative effects from attribution analysis were removed from the mutation space. This results in 116 mutation sites covering a sequence space of 8.8 × 10^77^.

##### Predictive models

Similar to the second round, we evaluated nine pLMs and the ev+onehot model using the activity and stability data from the third round. Principal component analysis (PCA) preprocessing and support vector regressor (SVR) were selected as the head models for the pLM models. The datasets were split into training and test sets with 20 different seeds, and the Spearman correlation coefficient was calculated as the metric, with the mean and variance calculated to prioritize the models.

We found that ProT5-XL had the best performance for activity prediction and the third best performance for stability prediction. It was selected as the predictive model of the two properties in the RL process because it performed well but much faster than ESM-3B.

##### Prior model

The AlphaVariant prior model in the third round is similar to that of the second round. The difference is that this prior model was fine-tuned on the sequence-activity data in the third round.

##### IP score calculation

The IP conflict checker returns a binary outcome per variant indicating whether any matching patents were identified (as detailed in **Supplementary Note 3**). For the multi-objective RL process, this binary outcome was inverted to yield the final IP score. A score of one indicates IP conflict avoidance for IP-free variants, while a score of zero indicates an IP conflict.

##### Multi-objective RL optimization

In the third round, three properties, activity, stability, and IP-free, were set as the optimization goals in the RL process. These three scores were combined as a single score to be used in the reinvent loss (**Equation (6)**). The activity and stability scores were predicted by the ProtT5-XL models and transformed into the range of zero to one. The IP score calculated above is added by one and then multiplied with the mean of the activity and stability scores to obtain the merged score, so that IP-free variants get additional bonus over the ones with IP conflicts.

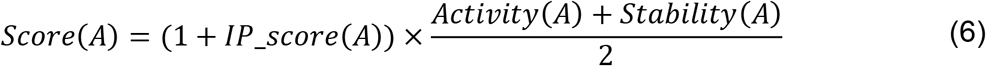

##### Library design

In the third round, we collected all the variants generated by the agent and filtered them by IP-free, activity and stability models to obtain variants for experimental validation. A step-by-step filtering procedure was implemented with the IP-free, ProtT5-XL, and ev+onehot models utilized in the first step and an ensemble of three pLMs and ev+onehot in the second step. Finally, the MD simulations were run at pH 10.5 and 85°C for 100 ns, and 32 variants were selected as the experimental library.

### Experimental protocol in the first two rounds of savinase optimization

#### Strains and plasmids

The gene encoding the predomain and mature forms of savinase, with the SacB signal peptide, was cloned and inserted into the pWB980 vector under the control of the p43 promoter. The full plasmid sequence is provided in the supplementary data. The *E. coli JM101* strain was used for cloning and library construction. The recombinant plasmid was then transformed into the *B. subtilis WB600* strain for protein expression.

#### Transformation of Bacillus subtilis

The WB600 strain was streaked onto Luria–Bertani (LB) agar plates without antibiotics and incubated at 37°C for 12 hours. Each single colony was picked and inoculated into 2 mL of SPI medium, followed by incubation at 37°C in a shaking incubator for 12 hours. 100 µL of the overnight culture was transferred into 5 mL of fresh SPI medium and incubated at 37°C for 4– 5 hours. Afterward, 200 µL of the culture was transferred into 2 mL of SPII medium and further incubated at 37°C with shaking at 220 rpm for 1.5 hours. Then, 0.1 mM EGTA was added to the culture, and the mixture was incubated at 37°C with shaking at 220 rpm for 10 minutes. The culture was then aliquoted into 1.5 mL microcentrifuge tubes, with 500 µL per tube. To each tube, 5–10 µL of a plasmid (verified by sequencing) was added. The tubes were incubated at 37°C with shaking at 220 rpm for 2 hours. After incubation, the cells were collected via centrifugation at 5000 rpm, and the resulting pellet was resuspended in 100 µL of the remaining liquid and plated onto 2% milk agar plates containing kanamycin. The plates were incubated at 37°C overnight. SPI medium (10 mL): 200 µL of CAYE solution, 200 µL of 50% glucose solution, 4.8 mL of SPA salt solution, and 4.8 mL of SPB salt solution. SPII medium (6 mL): 60 µL of 50 mM CaCl₂ solution, 60 µL of 250 mM MgCl₂ solution, and 5.88 mL of SPI medium. SPA Salt Solution: 0.4% (NH₄)₂SO₄, 2.8% K₂HPO₄·3H₂O, 1.2% KH₂PO₄, 0.2% trisodium citrate dihydrate. SPB Salt solution: 0.04% MgSO₄·7H₂O. CAYE Solution: 2% casein acid hydrolysate and 10% yeast extract.

#### The expression and detection of savinase activity

Each single colony exhibiting a hydrolysis cycle was picked from a 2% defatted milk agar plate and inoculated into 4 mL of LB medium (Kan+). The cultures were then grown in a shaking incubator at 37°C and 220 rpm for 24 h to allow protein expression in a constitutive manner. After expression, the culture was centrifuged at 4°C and 4000 rpm for 15 minutes to collect the supernatant. The supernatant was diluted 100-fold with borax-NaOH buffer (4.77 g/L Na_2_B_4_O_7_·10H_2_O, 1.72 g/L NaOH, 0.444 g/L CaCl_2_, pH adjusted to 10.0). The reaction mixture was prepared in a transparent 384-well plate as follows: 5 µL of 40 mM AAPF, 35 µL of borax-NaOH buffer, and 10 µL of the diluted supernatant, for a total volume of 50 µL. The absorbance at 410 nm was immediately monitored via a microplate reader (Tecan) set to 40°C. Measurements were taken every 30 s for 40 cycles. The ΔOD410/min value of each sample was calculated, and the ratio of the sample value to the value of the wild-type enzyme was used as the experimental activity.

#### Construction of savinase gene libraries

The primers used for library construction were ordered from GeneWiz, and 2× Fanta Flash Master Mix (Vazyme) was used for PCR assembly and library construction. The sequences of primers used are provided in the **Supplementary Table 3–6**. The computational library construction process (using the R1-L2 library as an example) was as follows: the primers with degenerate codons were mixed in groups of five in the following combinations: p1–p6, p5–p10, p9–p14, p13–p18, and p17–p22, and the proportion was adjusted according to the diversity (6×2 µL). The total reaction volume was 25 µL, and the PCR program consisted of the following conditions: 95°C for 30 s, followed by 35 cycles of 95°C for 10 s, 55°C for 10 s, and 72°C for 5 s. The PCR products were recovered via agarose gel electrophoresis to obtain fragments (F1–F6). The recovered fragments were divided into two groups (F1–F3, F3–F5) and mixed at an equimolar concentration of 10 µM. The total reaction volume was 50 µL, and the PCR program was as follows: 95°C for 30 s, followed by 35 cycles of 95°C for 10 s, 57°C for 10 s, and 72°C for 10 s. The PCR products were again recovered via agarose gel electrophoresis to obtain two long fragments. The two long fragments were mixed in equal amounts of 10 µM. Primers at both ends of the full-length gene were added at 2 µL each to the reaction mixture, and the total reaction volume was 50 µL. The PCR program was as follows: 95°C for 30 s, followed by 35 cycles of 95°C for 10 s, 57°C for 10 s, and 72°C for 15 s. The full-length gene library fragments were then recombined with the vector fragments via the ClonExpress Ultra One Step Cloning Kit V2 (Vazyme). The recombinant products were transformed into *E. coli* JM101 and plated on agar plates, and colonies were collected to extract plasmids, generating the library plasmid. Error-prone PCR was performed via the Controlled Error-prone PCR Kit (BTN160903) according to the manufacturer’s instructions. Wild-type savinase was used as the template for random mutagenesis of the full-length gene. The error-prone PCR products were purified and then recombined with the vector, followed by transformation into *E. coli* JM101. The ep-PCR library plasmids were then extracted from the transformed colonies. The site saturation mutagenesis library (sites 211, 212, 216) was constructed using primers containing NNK codons for PCR amplification. A full-length plasmid fragment with homologous arms was obtained and subjected to homologous recombination. The recombinant plasmid was then transformed into *E. coli* JM101, and colonies were collected for plasmid extraction.

#### High-throughput screening of savinase variants

The constructed library was transformed into *B. subtilis WB600* competent cells as previously described. The transformants were plated on 2% skim milk agar plates supplemented with kanamycin to select for successful transformants, which were subsequently screened for halo formation. Single colonies exhibiting a clear halo were picked and inoculated into 500 μL of LB medium (Kan+) in 96-deep-well plates. The cultures were incubated at 37°C for 24 h on a high-speed shaker at 500 rpm. Following incubation, the plates were centrifuged at 4000 rpm for 10 min at 4°C, and the supernatants were collected. The supernatants were diluted 100-fold with borax-NaOH buffer in new 96-well plates. Protease activity was then determined by measuring the ΔOD410/min via the method described above. The activity of each variant was compared to that of the wild-type enzyme. For the first round of directed evolution, approximately 1300 single colonies from each library were screened, and all single colonies were sent for Sanger sequencing by Genewiz. For the second round of directed evolution, approximately 1500 single colonies were screened, and the 150 variants with the highest activity were selected for Sanger sequencing. Samples with an activity value greater than three were selected for rescreening (with triplicate measurements) to confirm their activity.

#### Savinase catalytic kinetics assay

To determine the K_m_ and k_cat_ values of high-activity samples, plasmids encoding the top five active samples (from both rounds of evolution, a total of 10 samples) with a six-His-tag were constructed and transformed into the *B. subtilis WB600* strain. After 48 hours of expression at 37°C, the culture was centrifuged at 4000 rpm, and the supernatant was collected. The alkaline protease content in the supernatant was quantified via western blotting. For kinetic parameter estimation, the reaction rates were measured at a designated temperature (40°C) using a series of substrate concentrations (0.25, 0.5, 1, 2, and 4 mM). The initial velocity values at each specific substrate concentration were fitted to the Michaelis‒Menten equation via nonlinear regression. Each sample was measured in triplicate.

### Experimental protocol in the third round of savinase optimization

#### Bacterial strains, plasmids, and growth conditions

The expression of the *wt* Savinase gene and its engineered variants was driven by the amyE gene promoter derived from *B. subtilis ATCC 6051a*. The terminator sequence was from the *Bacillus amyloliquefaciens* α-amylase gene (amyQ). A synthetic DNA fragment encompassing the full-length wt savinase gene and the amyQ terminator was commercially synthesized by GenScript. The pUC57-BsaI-free plasmid, which was created by introducing a point mutation to eliminate the BsaI restriction site in the original pUC57 plasmid, served as the foundational vector for all subsequent constructs.

Unless otherwise specified, all *B. subtilis* strains were cultured in LB media (1% tryptone, 0.5% yeast extract, 1% NaCl) at 37°C with continuous shaking at 220 rpm. For the selection and screening of transformants, LB agar (1.5% agar) supplemented with 2% (w/v) skim milk and 5 µg/mL chloramphenicol was used.

#### Construction of the integration vector

The pUC57-BsaI-free plasmid was first linearized via enzymatic digestion with BamHI and HindIII. The resulting 2674 bp fragment was purified from an agarose gel. The upstream (amyE-5’) and downstream (amyE-3’) homologous arms of the amyE locus were amplified from the genomic DNA of *B. subtilis ATCC 6051a* (**Supplementary Table 7**). The amplification of amyE-5’ with the primers amyE-5’-F and amyE-5’-R yielded a 668 bp product, whereas the amplification of amyE-3’ with the primers amyE-3’-F and amyE-3’-R produced a 671 bp product. These two amplicons, along with the linearized 2674 bp pUC57-BsaI-free fragment, were assembled via a three-fragment recombination reaction via the CloneZ enzyme kit (GenScript), yielding the plasmid pUC57-BsaI-free-amyE. This intermediate vector was designed with two BsaI restriction sites located between the amyE-5’ and amyE-3’ fragments to facilitate subsequent cloning steps.

To construct the final integration vector, pUC57-BsaI-free-amyE was linearized with BsaI, generating a 3876 bp fragment. The coding sequence for the subtilisin variant, including the *Bacillus lentus* subtilisin 309 signal peptide, propeptide, and mature peptide, along with the amyQ terminator, was amplified from the synthetic Sav-wt gene template via the primers Sav-F and Sav-R, resulting in a 1273 bp PCR product. A chloramphenicol resistance cassette (1264 bp) was amplified from the genomic DNA of *Bacillus licheniformis ATCC 14580* via the primers Cm-F and Cm-R. These three DNA fragments (3876 bp, 1273 bp, and 1264 bp) were then assembled via the CloneZ recombination enzyme to generate the final integration plasmid, designated pUC57-Sav-Cm-amyE (Supplementary Information). The sequences of primers used for plasmid construction are detailed in Table 1. The amyE promoter, located at the 3’ end of the amyE-5’ fragment, is directly fused to the subtilisin 309 signal peptide to drive expression.

#### Construction of B. subtilis expression strains

The host strain for protein expression, A164Δ4, was constructed as previously described (Genscript (Nanjing) Co., Ltd. Chinese Patent CN104073458A, 2014). The integration plasmid pUC57-Sav-Cm-amyE and its derivatives were introduced into competent A164Δ4 cells via a standard chemical transformation protocol for *B. subtilis*. Following transformation, the cells were incubated in LB medium for 2 hours at 37°C with shaking at 200 rpm to allow for homologous recombination and expression of the resistance marker. The cell culture was then serially diluted, and 100 µL of a 10^−2^ dilution was plated onto selective LB agar containing 2% skim milk and 5 µg/mL chloramphenicol. The plates were incubated overnight at 37°C. Colonies exhibiting a clear halo, indicative of protease activity, were selected for further analysis. The correct integration of the expression cassette into the amyE locus was confirmed by colony PCR and Sanger sequencing, resulting in the final expression strains.

#### Protein production and purification

For protein production, the *B. subtilis* strains were cultivated in shake flasks. Single colonies were used to inoculate 50 mL of LB seed medium in 250 mL flasks, which were incubated overnight at 37°C and 220 rpm. The overnight cultures were then used to inoculate 50 mL of AKP fermentation medium (10% glucose, 6% soybean meal, and 1% anhydrous Na_2_HPO_4_) in 250 mL flasks to an initial optical density at 600 nm (OD_600_) of 0.1. Fermentation was conducted for 96 hours at 37°C with shaking at 220 rpm. After fermentation, the culture broth was harvested, and the cells were removed via centrifugation. The resulting supernatant, containing the secreted protease, was then assayed for protease activity and subsequent characterization.

#### Industrial performance assays of savinase variants

To evaluate the industrial applicability of the engineered protease variants, a series of standardized assays were performed. The alkaline protease activity was quantified based on the method described in the Chinese National Standard GB 1886.174-2016 for enzyme preparations used in the food industry. The thermal stability of the proteases was assessed by measuring the residual activity after incubation at 85°C for 3 minutes. The cleaning performance was evaluated via a standardized detergency test (JB02) in a de-enzymed base detergent solution. The assay was performed with a protease concentration of 250 ppm for 20 minutes at 30°C. These characterization studies were designed to ensure that the engineered protease variants possess the requisite properties for demanding industrial applications.

## Supporting information

Supplementary information of AlphaVariant

## Data availability

The benchmark datasets and savinase design three round datasets to reproduce the figures are submitted to Figshare (https://doi.org/10.6084/m9.figshare.33307125). All experimental data of this article are available through the corresponding authors upon reasonable request.

## Code availability

Source codes of AlphaVariant framework, the benchmark, and savinase design project will be open sourced on GitHub. The source code of DCG-libgen publicly available on GitHub (https://github.com/charlesxu90/dgc-libgen). The checkpoints, including VariantGPT model and predictive models, and the experimental datasets will be submitted to Figshare.

## Acknowledgements

We acknowledge the computational resources provided by the Supercomputing Laboratory at King Abdullah University of Science & Technology (KAUST). We thank Bestzyme for providing industrial context and validation studies.

## Author Contributions

X.X., H.Y., and Z.G. contributed equally to this work. X.X. developed the AlphaVariant framework, implemented the VariantGPT model and predictive algorithms, and conducted computational design. H.Y. and Z.G. designed and performed the experimental validation studies in the first and second rounds. X.Z., G.Z., M.H., B.X., F.Z., F.H., and A.B. contributed to the experimental design and industrial validation in the third round. T.G., W.H., and C.F. assisted with the computational infrastructure and data analysis. J.W. and X.G. supervised the project and provided strategic guidance. All the authors contributed to manuscript preparation and revision.

## Extended Data Figures

**Extended Data Fig. 1.**
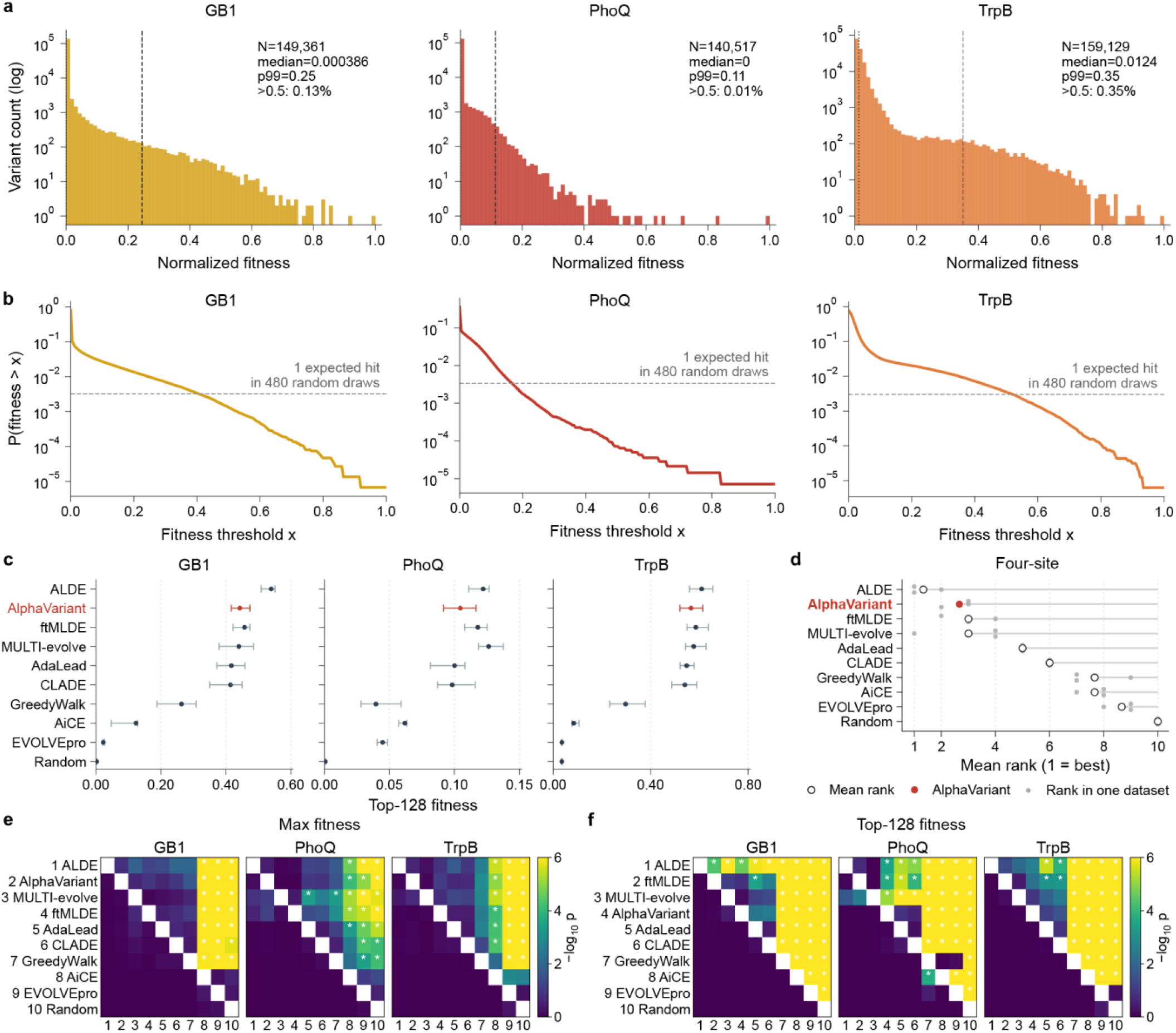
Extended benchmark comparison of AlphaVariant against state-of-the-art methods on four-site datasets. **a**, Distribution of normalized fitness for the GB1, PhoQ, and TrpB benchmark datasets. The median (dotted line) is nearly zero and the p99 is marked as dashed line. The y-axis is on a logarithmic scale. **b**, The probability curves illustrating the likelihood of finding a variant with fitness greater than threshold x, *P*(*fitness* > *x*), across the fitness spectrum for GB1, PhoQ, and TrpB. The dashed grey line represents the threshold for one expected hit in 480 random draws. **c,** Comparison of top-128 fitness (the mean fitness of the top 128 variants discovered) across all benchmarked methods for four-site problems (GB1, PhoQ, and TrpB). Error bars represent the standard deviation across 30 independent runs. **d**, Mean rank comparison across all benchmarked methods for four-site problems, evaluated on top-128 fitness. Each grey dot represents the rank of a method on a single dataset; the open circle represents the mean rank across datasets; and the filled red dot highlights AlphaVariant’s mean rank. AlphaVariant (highlighted in red) achieves highly competitive top-128 fitness across all four-site datasets. **e,f**, Pairwise statistical comparison matrices (Wilcoxon signed-rank test, −log_10_ p-value) for median max fitness (**e**) and top-128 fitness (**f**) across four-site datasets. Methods are ordered by rank (1 = best). Brighter (yellow) cells indicate statistically significant differences; darker (purple) cells indicate non-significant differences.

**Extended Data Fig. 2.**
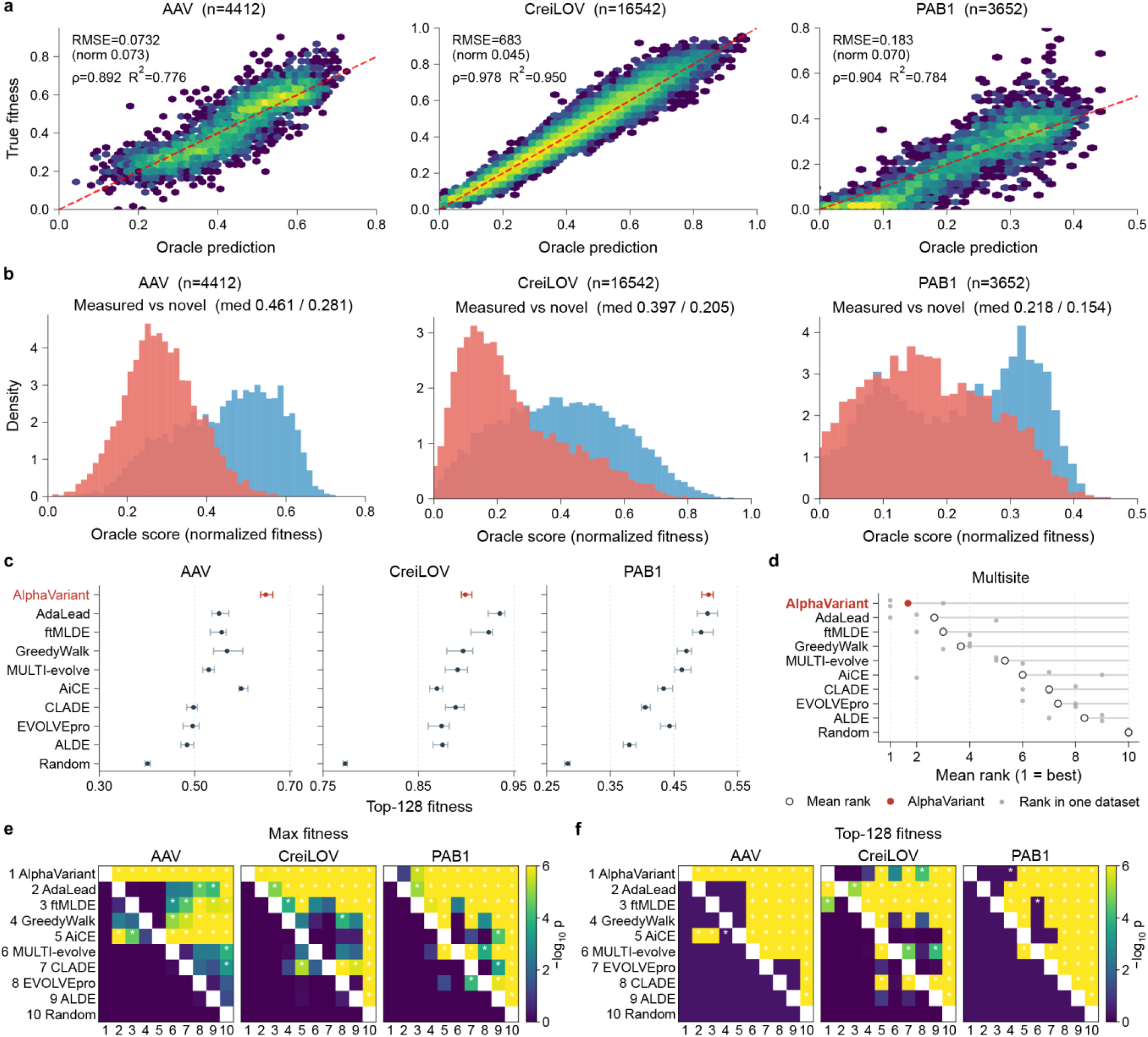
Extended benchmark comparison of AlphaVariant against state-of-the-art methods on multi-site datasets. **a**, Hexbin scatter plots comparing oracle-predicted fitness against true fitness for the AAV, CreiLOV, and PAB1 datasets. Performance metrics including RMSE, normalized RMSE, Spearman correlation (ρ), and R2 are shown. **b**, Density histograms comparing the oracle score (normalized fitness) distributions of the measured test set (blue) versus the novel unmeasured set (red) for AAV, CreiLOV, and PAB1, demonstrating the shift in distribution toward lower fitness for the novel generated variants. **c**, Comparison of top-128 fitness (the mean fitness of the top 128 variants discovered) across all benchmarked methods for multi-site problems (AAV, CreiLOV, and PAB1). Error bars represent the standard deviation across 30 independent runs. **d**, Mean rank comparison across all benchmarked methods for multi-site problems, evaluated on top-128 fitness. Each grey dot represents the rank of a method on a single dataset; the open circle represents the mean rank across datasets; and the filled red dot highlights AlphaVariant’s mean rank. AlphaVariant achieved the best mean rank on multi-site problems. **e,f**, Pairwise statistical comparison matrices (Wilcoxon signed-rank test, −log_10_ p-value) for median max fitness (**e**) and top-128 fitness (**f**) across multi-site datasets. Methods are ordered by rank (1 = best). Brighter (yellow) cells indicate statistically significant differences; darker (purple) cells indicate non-significant differences.

**Extended Data Fig. 3.**
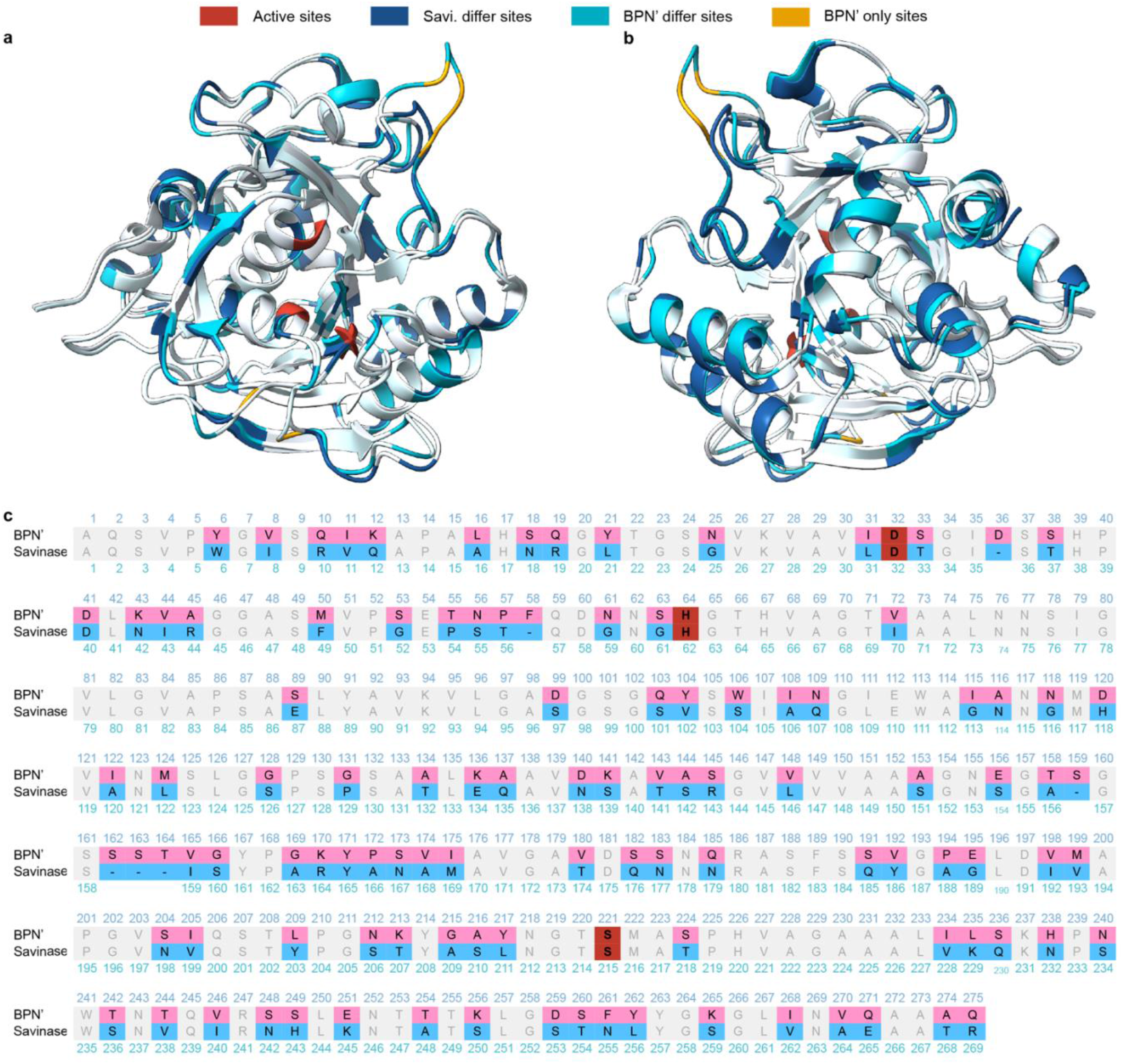
Structure and sequence alignment of BPN’ and savinase. **a,b**, Front (**a**) and back (**b**) view of the structure alignment. Red, the active sites; dark blue, missense mutation sites in savinase; cyan, missense mutation sites in BPN’; gold, BPN’ only sites, which are deleted in savinase. **c**, The sequence alignment. Red, active sites; blue, differ sites in savinase; pink, differ sites in BPN’; grey, identical residues.

**Extended Data Fig. 4.**
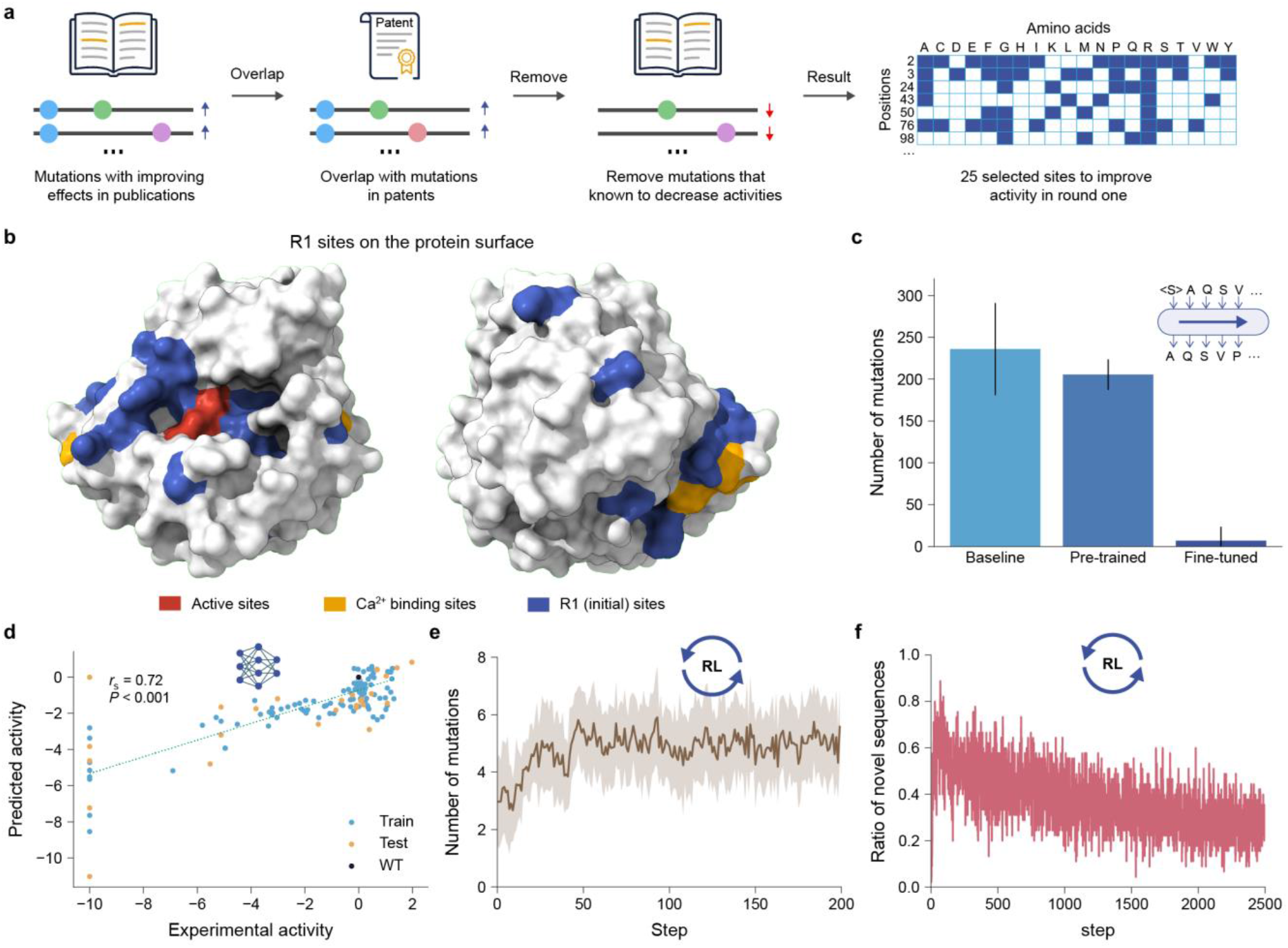
Additional analysis of *in silico* design in round one. **a**, Workflow for defining the R1 mutation space. Mutations with improving effects reported in the literature are first identified, then overlapped with mutations in existing patents. Mutations known to decrease activity are removed, yielding a final set of 25 selected sites to improve activity in round one, represented as a mutation space heatmap. **b**, Surface representation of the R1 selected sites from two orientations. Red, active sites; gold, Ca²⁺ binding sites; blue, R1 mutation sites. **c**, Number of mutations in the variants generated by the prior models. The distribution of the homology sequences is shown as the baseline. **d**, Evaluation of the activity model built from the publication dataset. A Spearman correlation of 0.72 was achieved on the test dataset. **e**, The average number of mutations of variants generated in the RL process. **f**, The ratio of novel sequences in each batch during the RL process.

**Extended Data Fig. 5.**
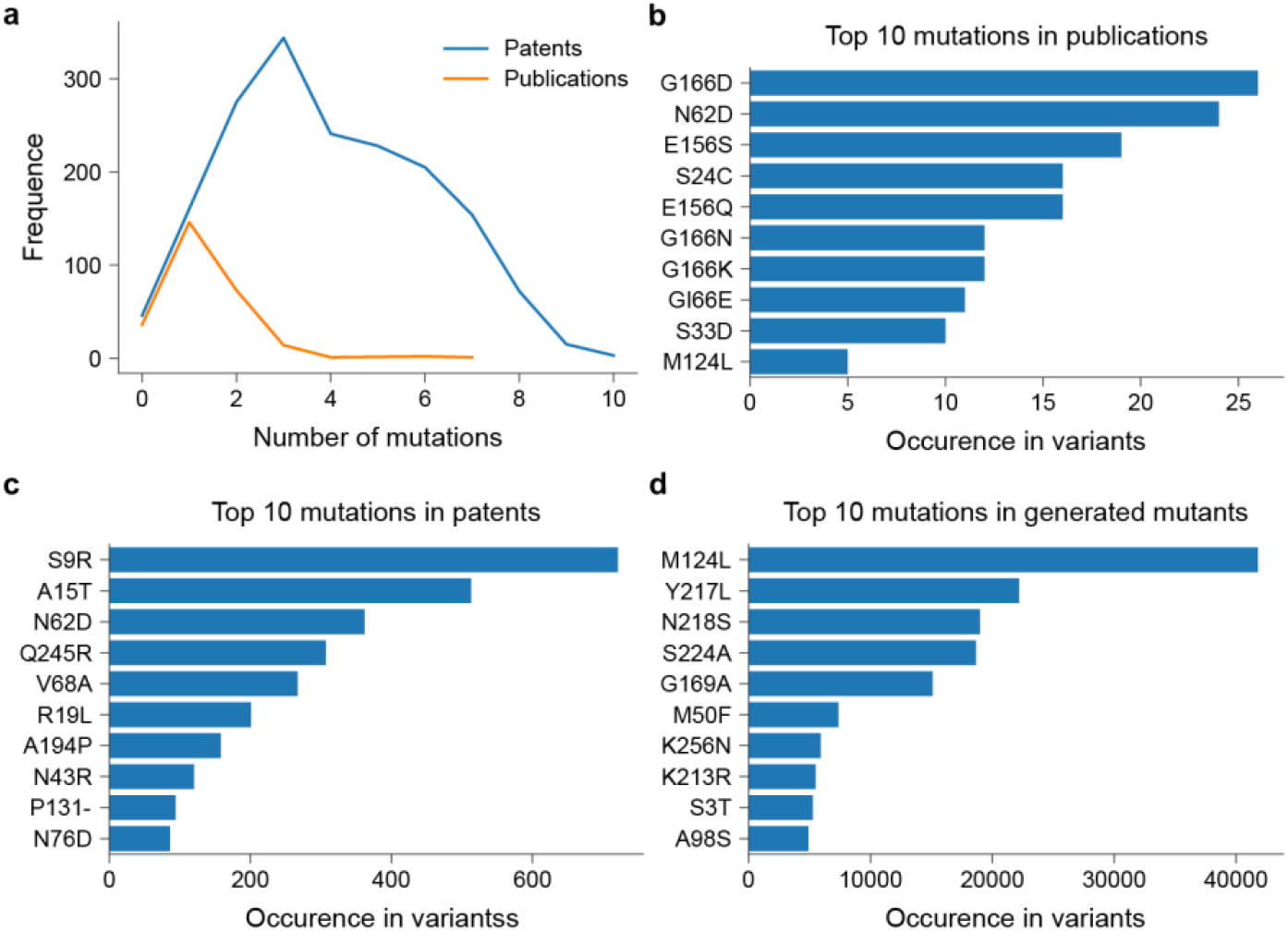
Analysis of mutation sources in round one design. **a**, Number of mutations of variants in patents (blue) and publications (orange). **b-d**, The top 10 most frequent mutations in publications (**b**), patents (**c**), and R1 generated variants (**d**). The difference between the generated variants from publications and patents highlights shift in mutational preferences explored by AlphaVariant.

**Extended Data Fig. 6.**
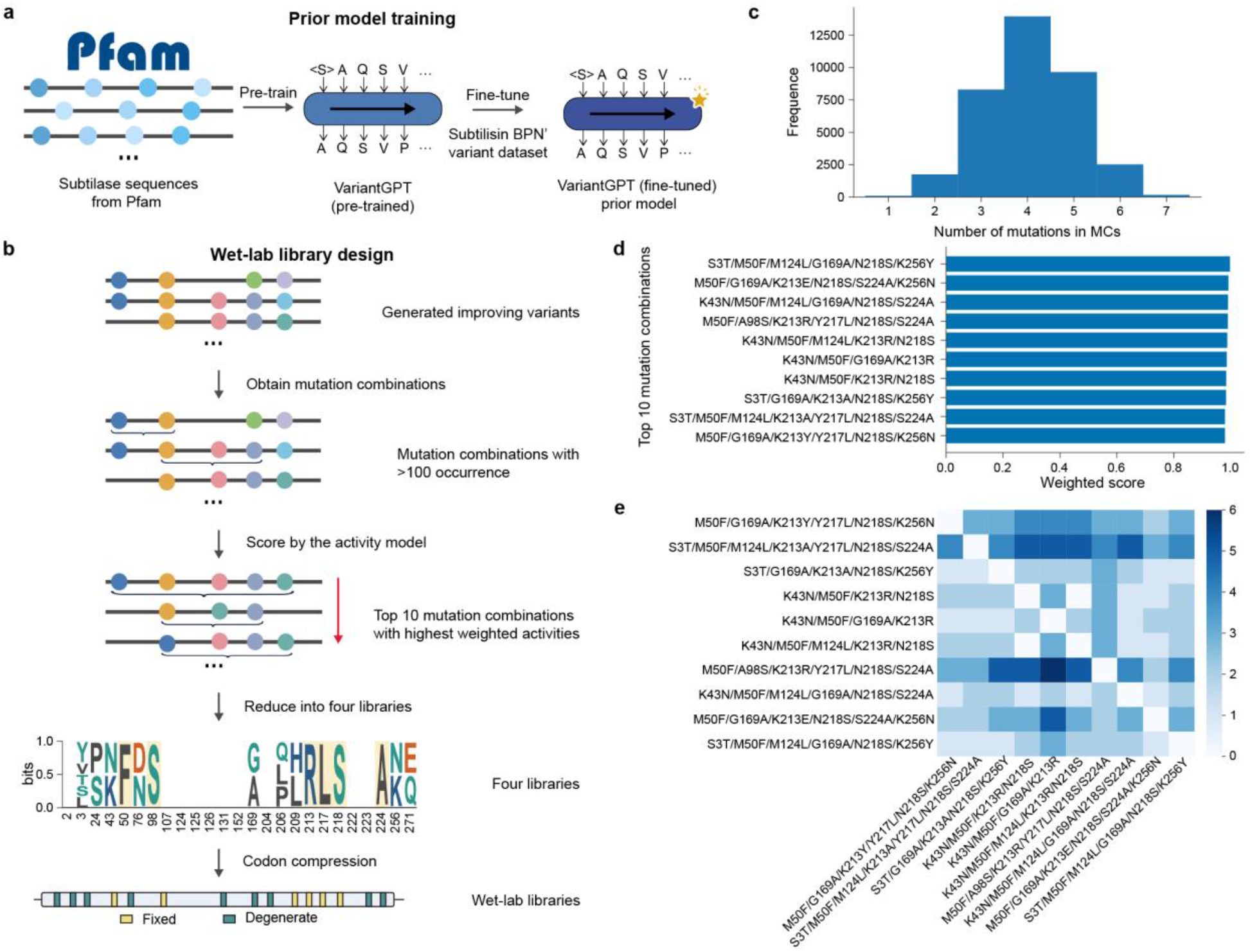
The prior model training and library design pipeline in round one. **a**, Workflow to train a VariantGPT model in the round one (R1). **b**, Workflow for wet-lab library design in R1. AI-generated improving variants are analyzed to extract high-occurrence mutation combinations (MC). These combinations are scored by the activity model, and the top 10 are selected. These are then reduced into four sequence libraries and finally compressed into degenerate codon libraries for wet-lab synthesis. **c**, Distribution of the number of mutations within the extracted MCs, centering around four to five mutations. **d**, The top 10 MCs ranked by their weighted activity scores. **e**, Heatmap visualizing the co-occurrence and overlap of specific mutations among the top 10 MCs.

**Extended Data Fig. 7.**
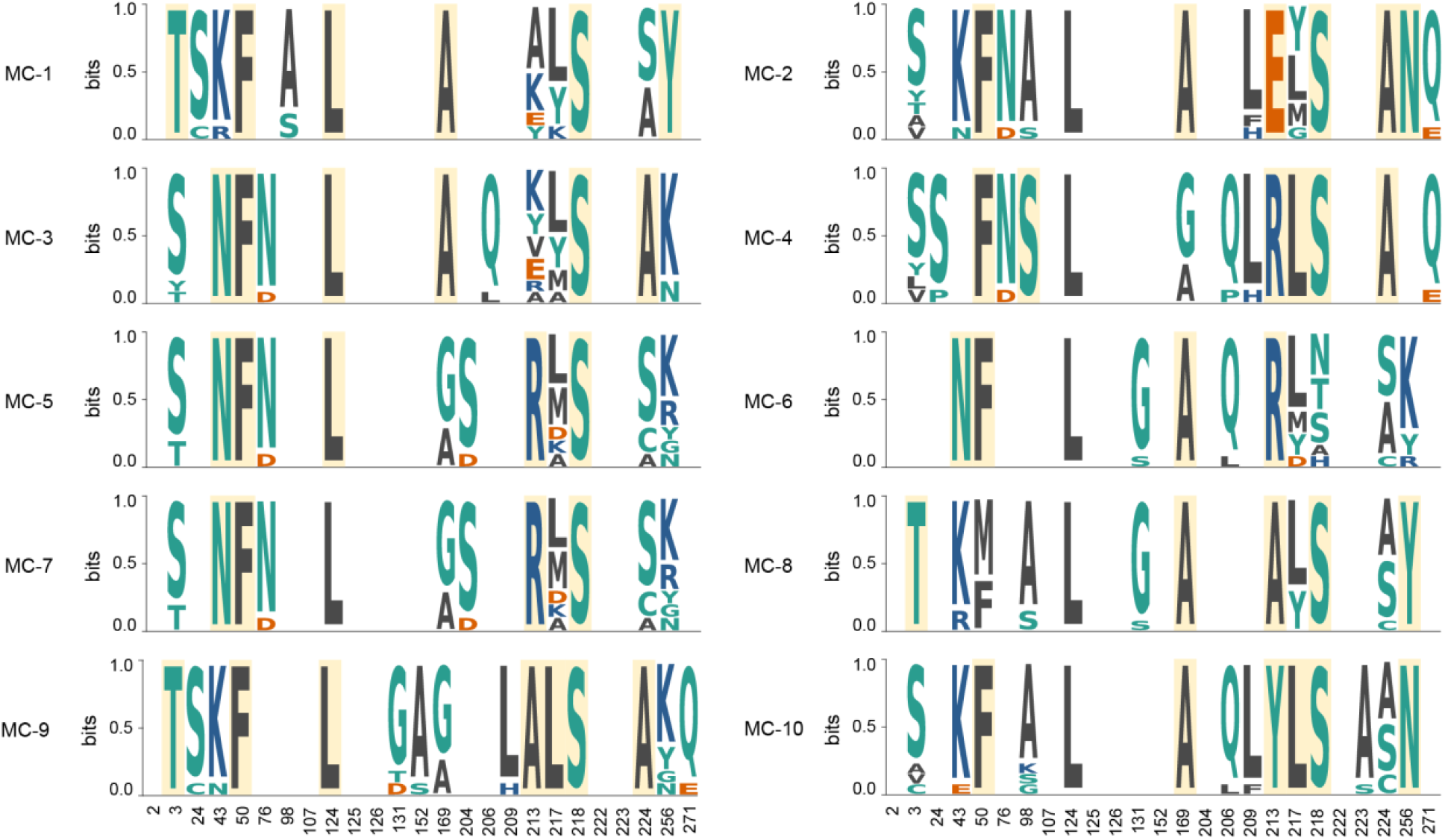
Libraries generated from top 10 mutation combinations in round one. For each of the top 10 MCs, we identified the concurrent mutations among the generated variants as associated sites. The corresponding amino acid frequencies were calculated to draw the sequence logos. The sequence logos representing the amino acid preferences at the selected mutation sites for the top 10 MCs (MC-1 to MC-10).

**Extended Data Fig. 8.**
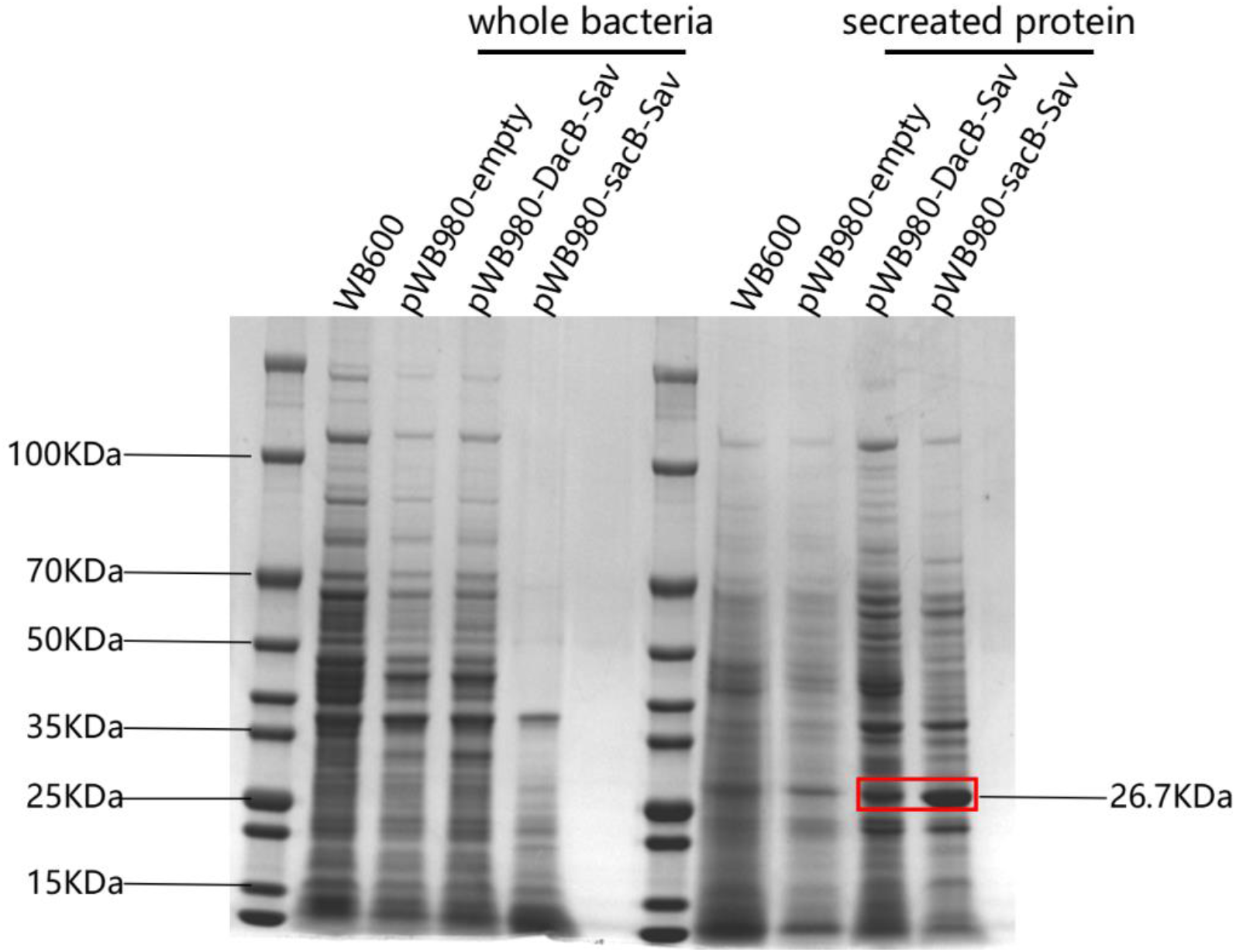
Validation of savinase expression and secretion. The blank WB600 strain and strains transformed with different plasmids were cultured in parallel for 24 hours. Whole cells and secreted protein (total protein in the medium) samples from the supernatant were collected. The supernatant was acidified by adding trichloroacetic acid (final concentration 13%) and incubated at 4°C for 12 hours. The precipitate was collected by centrifugation and washed three times with acetone. SDS-PAGE was performed to detect protein expression. Lanes include the WB600 host control, empty vector (pWB980-empty), and savinase expressed with two different signal peptides (pWB980-DacB-Sav and pWB980-sacB-Sav). The red box highlights the successful secretion of the mature savinase protein (∼26.7 kDa) in the culture supernatant.

**Extended Data Fig. 9.**
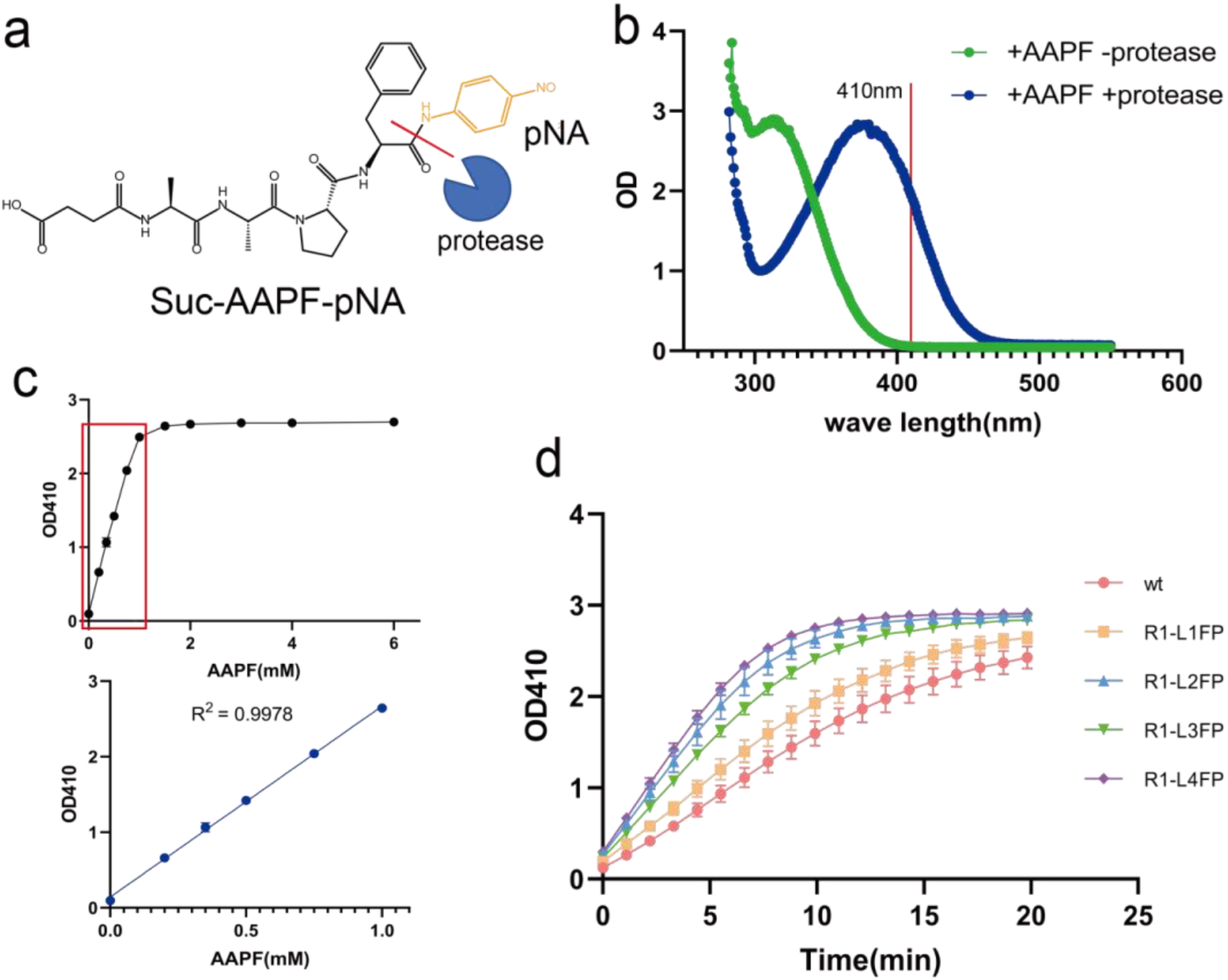
Validation of the high-throughput protease activity assay. **a**, Structure of suc-AAPF-pNA, with the protease cleavage site indicated by a red line. The yellow structure represents the colored product, pNA, generated upon cleavage. **b**, Absorbance spectra of the substrate (+suc-AAPF-pNA -protease, green) and the cleavaged product (+suc-AAPF-pNA +protease, blue). The vertical red line marks the 410 nm wavelength used for detection. **c**, Standard curve demonstrating the linear relationship between substrate concentration (and resulting pNA release) and absorbance at OD410 (R^2^ =0.9976). **d**, Kinetic absorbance curves (OD410 over time) for WT and fixed-position variants (R1-L1FP to R1-L4FP), validating the dynamic range and temporal resolution of the assay.

**Extended Data Fig. 10.**
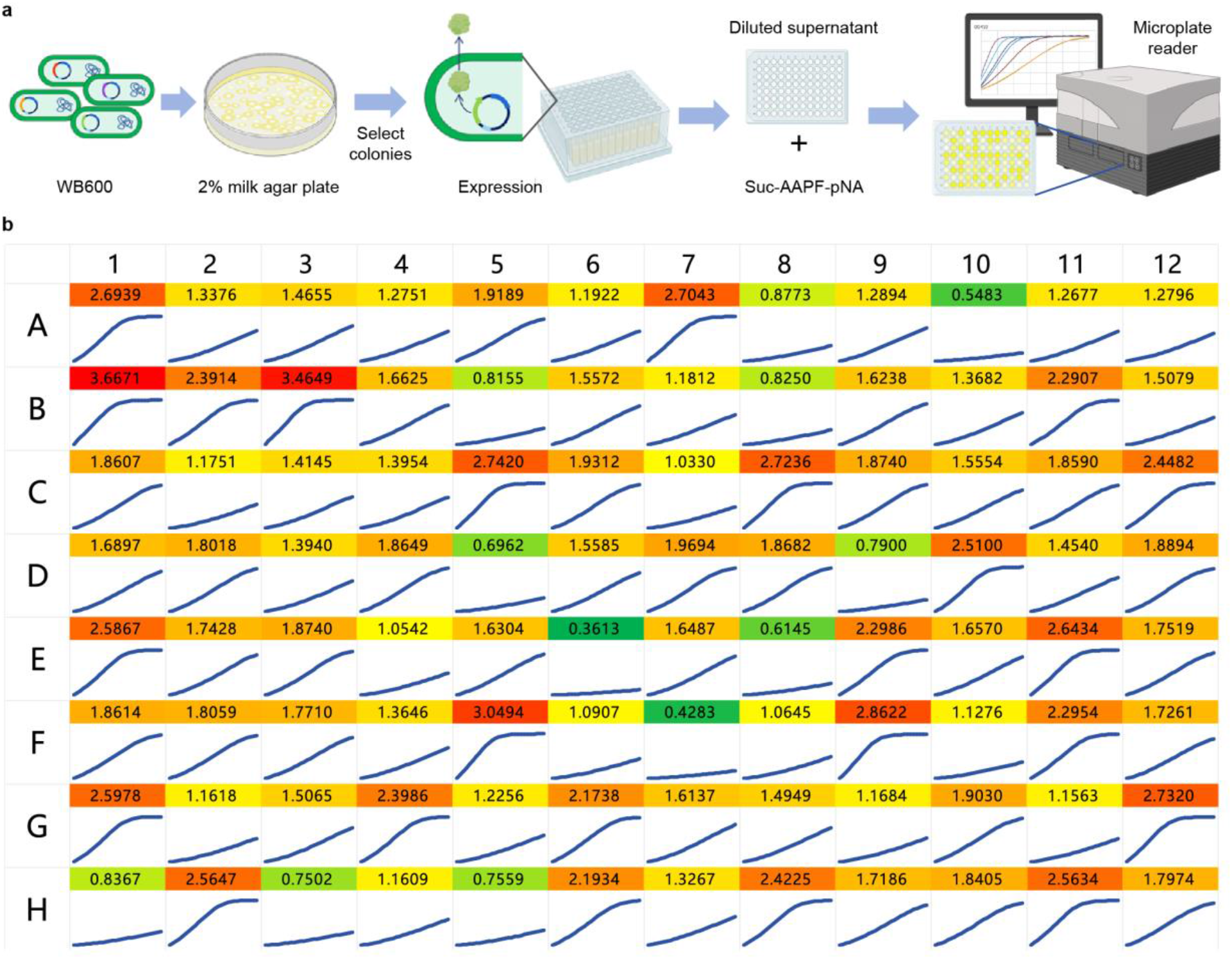
High-throughput protease activity assay and time-OD curve monitoring diagram. **a**, High-throughput protease activity assay workflow, including colony selection on 2% milk agar plates, expression in deep-well plates, incubation with Suc-AAPF-pNA, and microplate reader measurement. **b**, High-throughput OD-time curve for samples in 384-well plate (96 samples selected). The wild-type sample’s activity value (the average of the three samples in wells A1-A3), ΔOD410/min, is calculated based on the curve, and the enzyme activity fold increase is determined by dividing the sample activity by averaged wild-type activity. Values are color-coded by activity levels (red = high, green = low).

**Extended Data Fig. 11.**
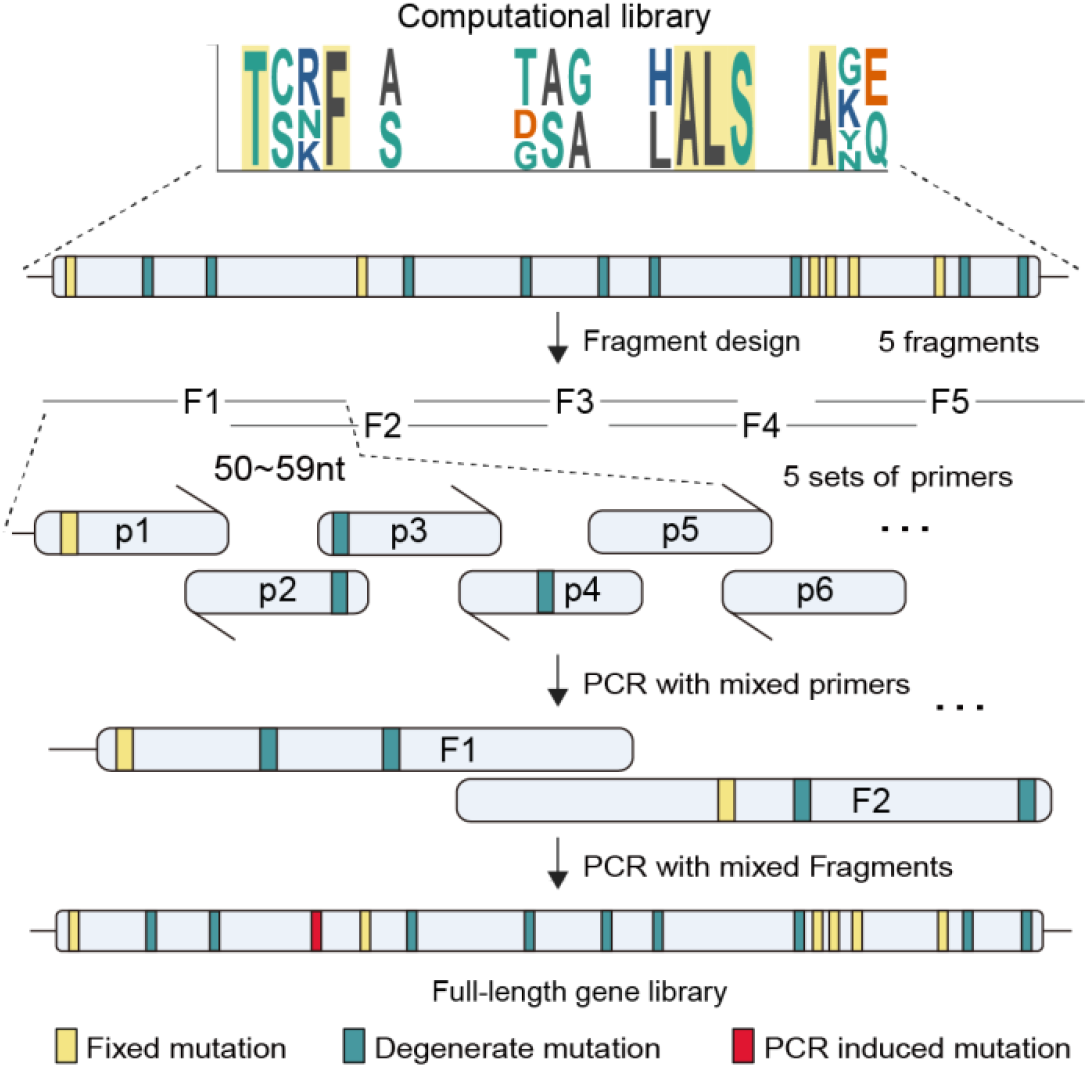
Workflow used to construct the DNA libraries from the computational libraries. The gene was split into five fragments, each amplified by PCR with degenerate primers, and then assembled through overlap-PCR, allowing new mutations for diversity.

**Extended Data Fig. 12.**
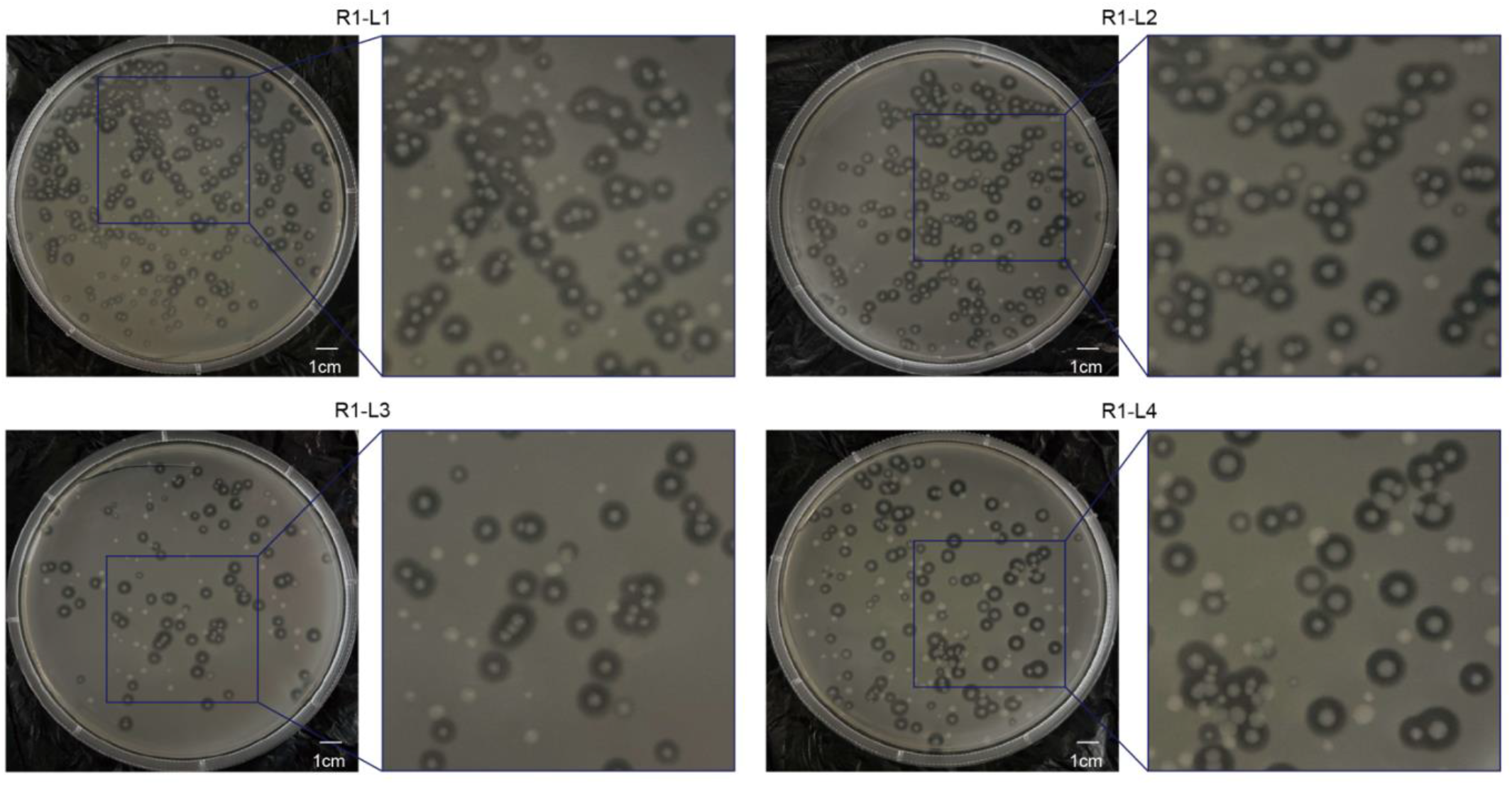
Agar plate display of computational libraries in round one. Representative colony plates from the first-round AlphaVariant-designed libraries, R1-L1–R1-L4. The computationally designed variant libraries were synthesized and assembled into experimental libraries, with zoomed regions showing individual transformant colonies for each condition. The resulting colony distributions indicate successful library construction and transformation for downstream screening.

**Extended Data Fig. 13.**
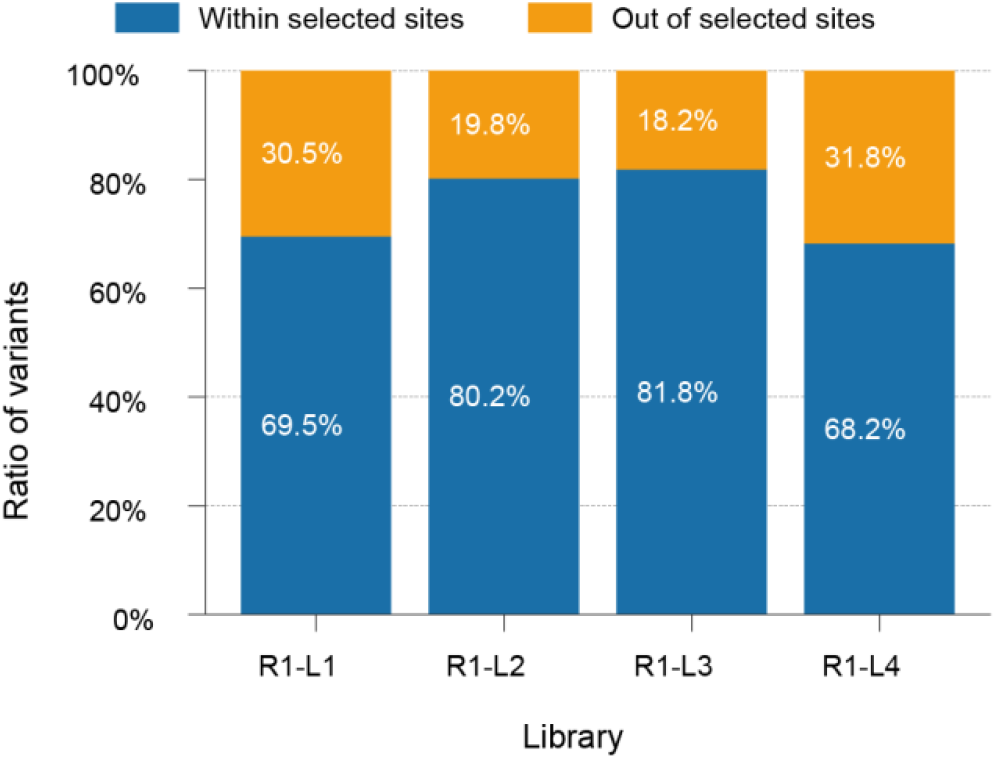
Mutation localization in R1 synthesized libraries. Stacked bar chart quantifying the proportion of variants containing mutations strictly within the R1 selected sites (blue) versus those containing unintended mutations outside the selected sites (orange, e.g., due to PCR errors) for libraries R1-L1 to R1-L4. The high percentage of on-target mutations (∼70-82%) confirms the fidelity of the library synthesis process.

**Extended Data Fig. 14.**
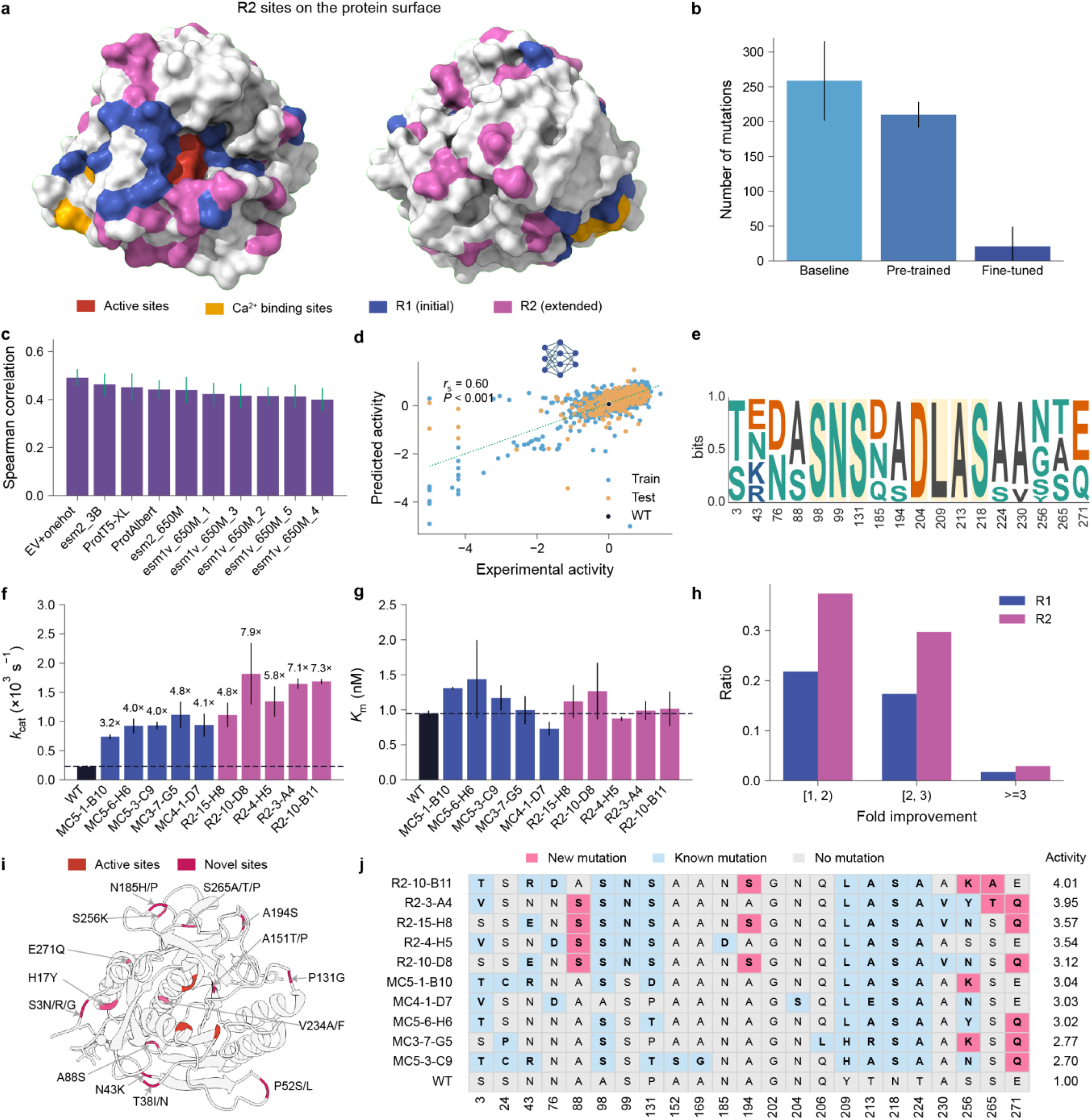
Additional results of the two rounds of design and experiments. **a**, Surface representation of explored sites in round two. Red, active sites; gold, Ca²⁺ binding sites; blue, R1 mutation sites; purple, R2 extended sites. **b**, Number of mutations in the variants generated by the R2 prior models. The distribution of the homology sequences is shown as the baseline. **c**, Benchmark protein fitness models on the activity data collected from R1. **d**, Evaluation of the ev+onehot activity model built from R1 activity dataset. A Spearman correlation of 0.60 was achieved on the test dataset. **e**, The library designed in the second round. **f**, Turnover number (*k_cat_*, ×10^−3^ s^−1^) of WT, top R1 variants (blue), and top R2 variants (purple). Numbers above bars indicate fold improvement over WT. **g**, Michaelis constant (*K_m_*, nM) of WT and top R1 and R2 variants, showing that improved catalytic efficiency is primarily driven by increased *k_cat_* rather than changes in substrate binding affinity. **h**, Fold improvement ratios of the variants in R1 and R2. **i**, Novel mutations with positive contributions in the two rounds. The novel mutations are not included in patents or publications and presented at least twice among the experimentally improved variants. **j**. Mutations in the top five variants from the two rounds. Pink, new mutations; light blue, known mutations; grey, unmutated positions. Activity fold improvement values are shown on the right (mean of three replicated experimental assays).

**Extended Data Fig. 15.**
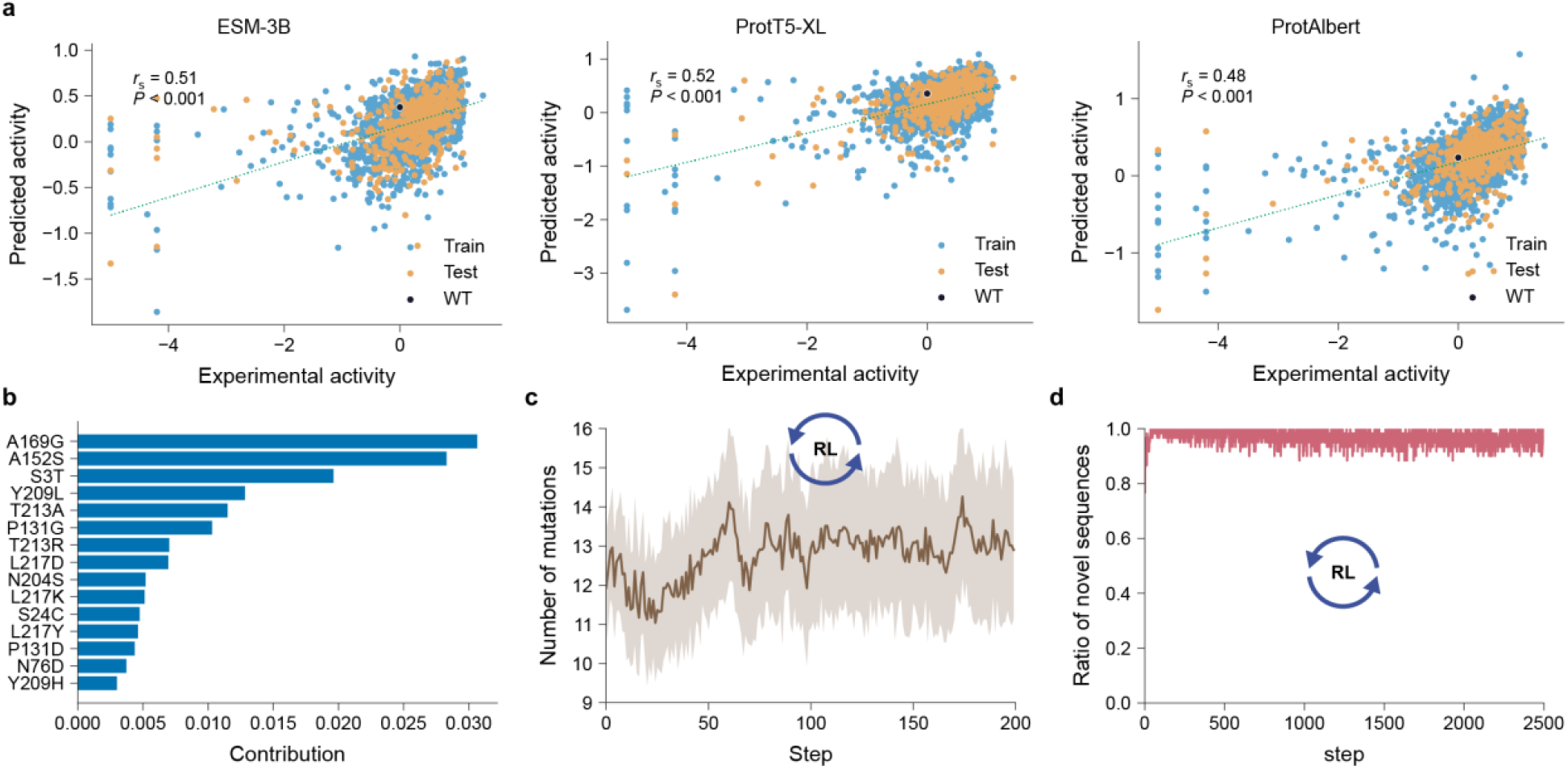
Evaluation of pLM activity models and RL optimizations in round two. **a**, Scatter plots of predicted versus experimental activity for the R2 predictive model using different protein language model embeddings: ESM-3B, ProtT5-XL, and ProtAlbert. Spearman correlation coefficients (r_s_) range from 0.48 to 0.52 (P<0.001). **b**, Shapley contribution analysis of mutations in R1 variants. It identified the specific mutations that most strongly drive the activity predictions, with A169G and A152S showing the highest contributions. **c**, Number of mutations during the RL steps. **d**, Ratio of novel sequences during the RL steps.

**Extended Data Fig. 16.**
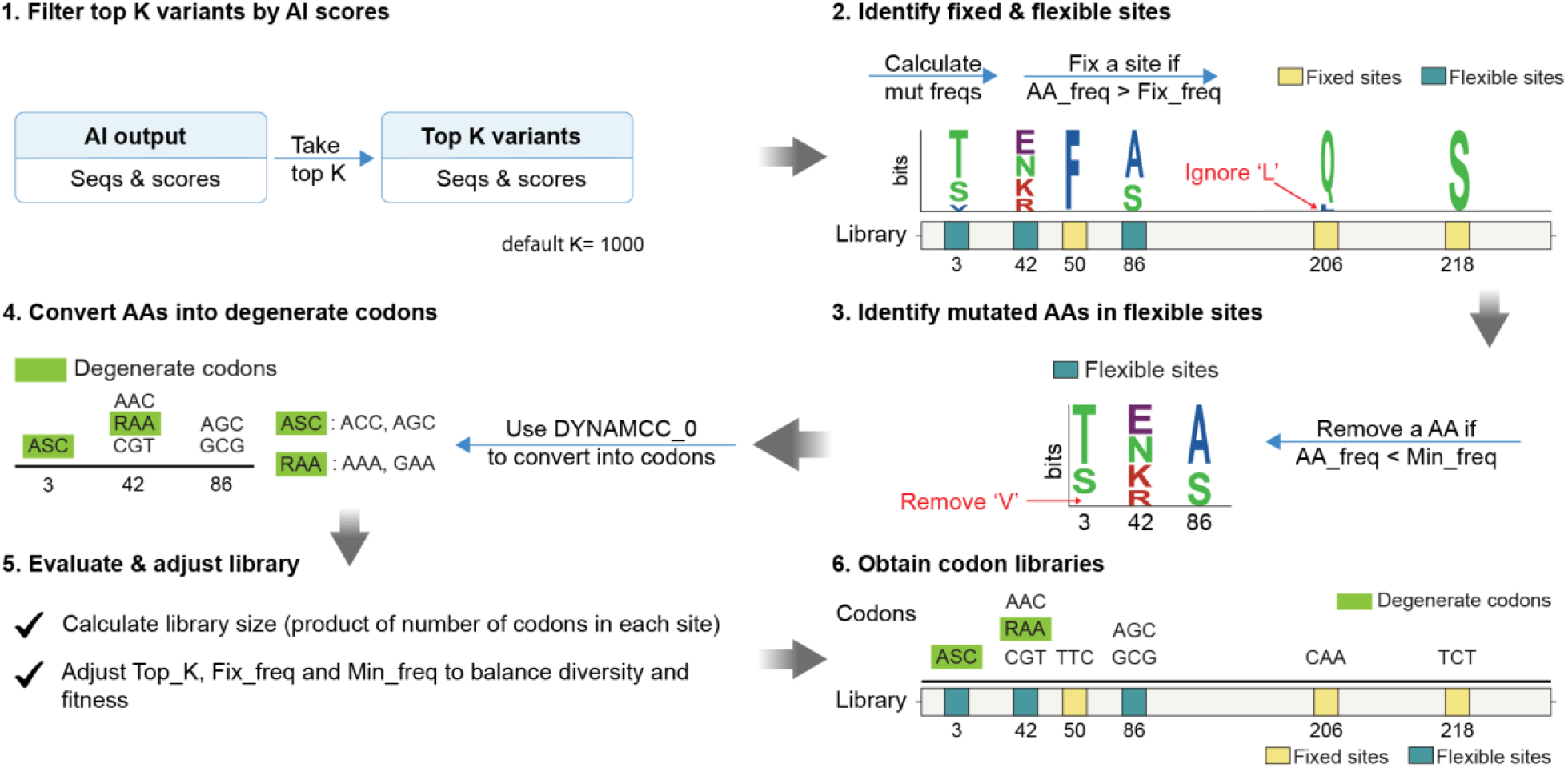
The workflow of DCG-Libgen to design degenerate codon libraries. The workflow to design a degenerate codon (DGC) library from AI outputs includes the f llowing steps: 1) filter the top K variants on the basis of AI scores; 2) obtain fixed and flexible sites by analyzing amino acid (AA) frequencies at each site, with an AA frequency greater than a threshold (Fix_freq) is fixed; 3) mutations with frequencies lower than the minimum frequency (Min_freq) are removed from flexible sites; 4) the amino acids are converted into codons via DYNAMCC_0; 5) the library size is calculated, and the parameters (Top K, Fix_freq, and Min_freq) are adjusted to fit the desired size; and 6) the final mutation sites with respective codons are used to design a wet-lab library.

**Extended Data Fig. 17.**
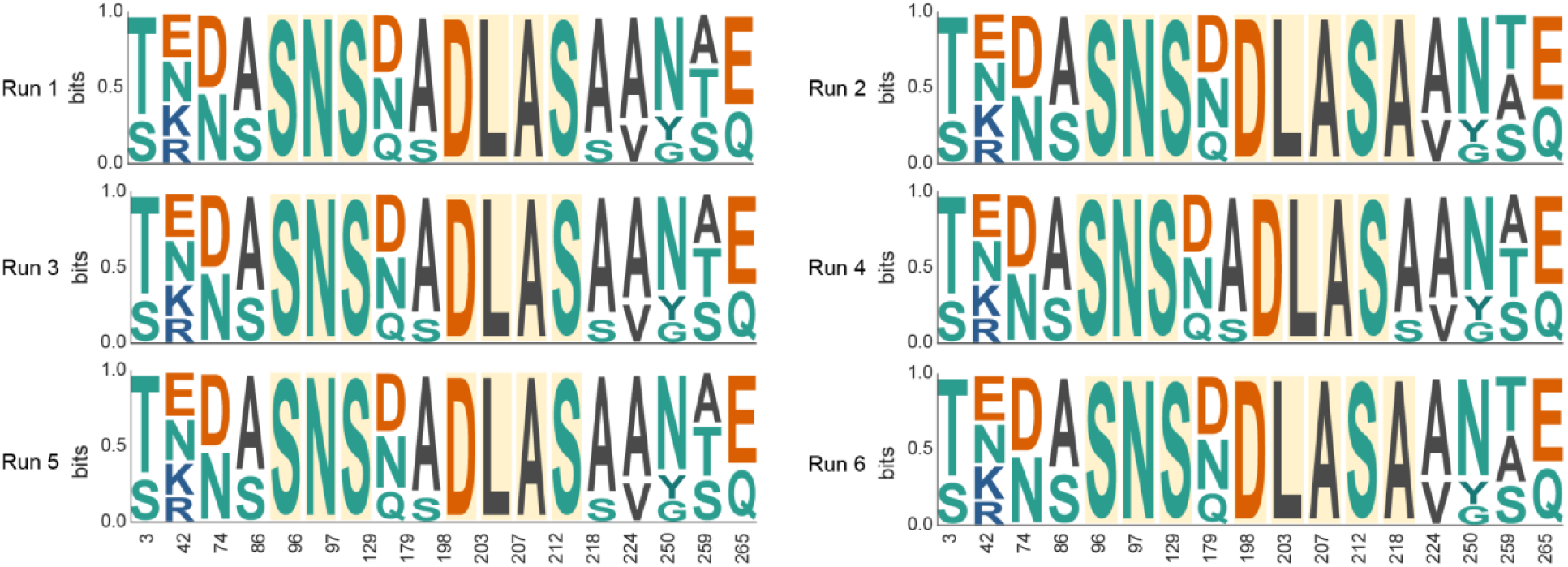
Libraries generated across six independent RL runs in the round two. For each of top six active variants in the first round, we ran RL to generate variants of improved activities. Sequence logos of the generated libraries from six independent RL runs (run 1 to run 6) for R2 are shown. The nearly identical amino acid distributions at the targeted positions across all six runs demonstrate the robustness and convergence of the RL optimization process.

**Extended Data Fig. 18.**
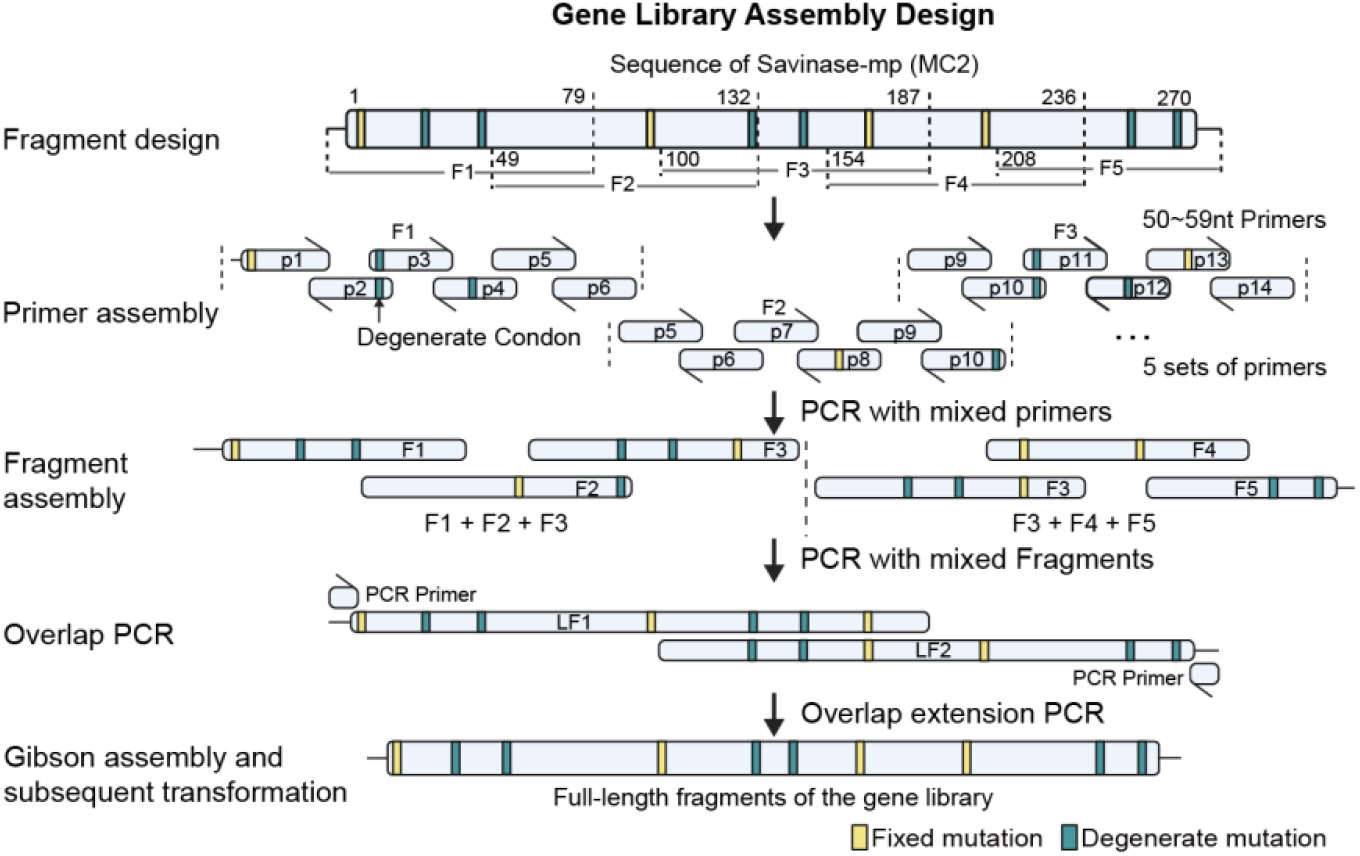
Schematic of the multi-fragment gene library assembly design. The full-length gene is computationally divided into five overlapping fragments (F1–F5). Sets of common and degenerate primers (50–59 nt) are designed to introduce both fixed (yellow) and degenerate (teal) mutations. These primers are assembled into fragments via PCR, and the fragments are subsequently joined using overlap-extension PCR to yield the full-length mutant library for Gibson assembly and transformation. This strategy enables combinatorial incorporation of designed mutations across multiple positions while maintaining compatibility with the computational libraries.

**Extended Data Fig. 19.**
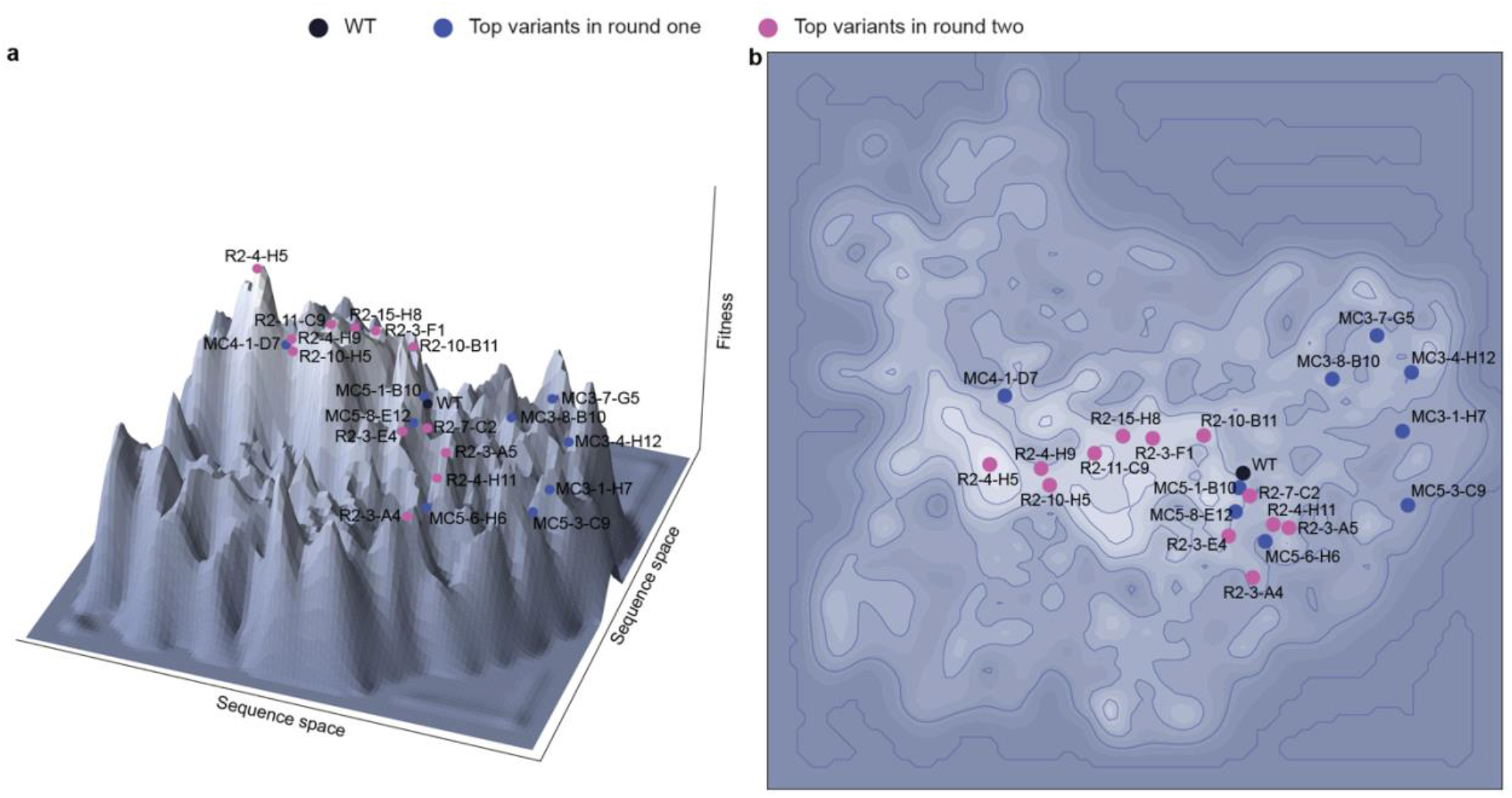
Schematic representation of the savinase’s fitness landscape in AAPF catalysis in the first two rounds. A total of 2,608 unique sequence–activity pairs were used to map savinase’s fitness landscape to catalyze Suc-AAPF-pNA. For each variant, the underlying sequence space is represented by a principal component analysis (PCA) of the embeddings extracted from the ProtT5-XL. **a**, Three-dimensional surface plot representing a projection of the sequence space against fitness. The WT (black dot), R1 top variants (blue dots), and R2 top variants (pink dots) are mapped onto the landscape, showing the progression toward higher fitness peaks in R2. **b**, Two-dimensional contour map (top-down view) of the same fitness landscape, illustrating the clustering and spatial relationship of the engineered variants relative to the WT sequence. Black, the wild-type savinase; blue, R1 top variants; pink, R2 top variants.

**Extended Data Fig. 20.**
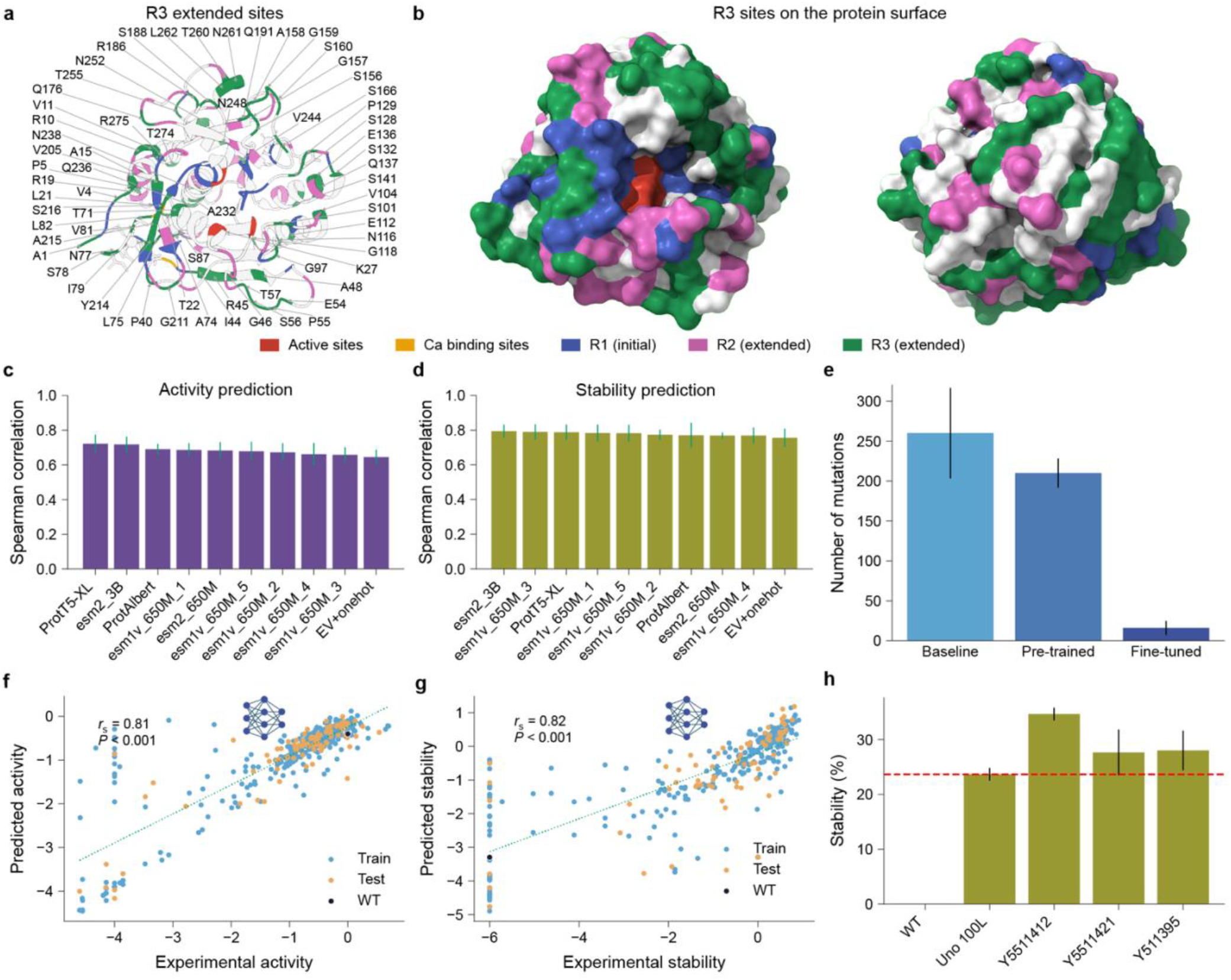
Additional results from the third round. **a**, The extended mutation sites in R3. **b**, Surface representation of R3 explored sites. Red, active sites; gold, Ca²⁺ binding sites; blue, R1 mutation sites; purple, R2 extended sites; green, R3 extended sites. **c**,**d**, Evaluation of protein fitness models on the variant-activity data (**c**) and stability data (**d**). **e**, Number of mutations in the variants generated by the prior models. **f**,**g**, Evaluation of the ProtT5-XL-based activity (**f**) and stability (**g**) models used in the RL process. **h**, Stability of the selected variants.

**Extended Data Fig. 21.**
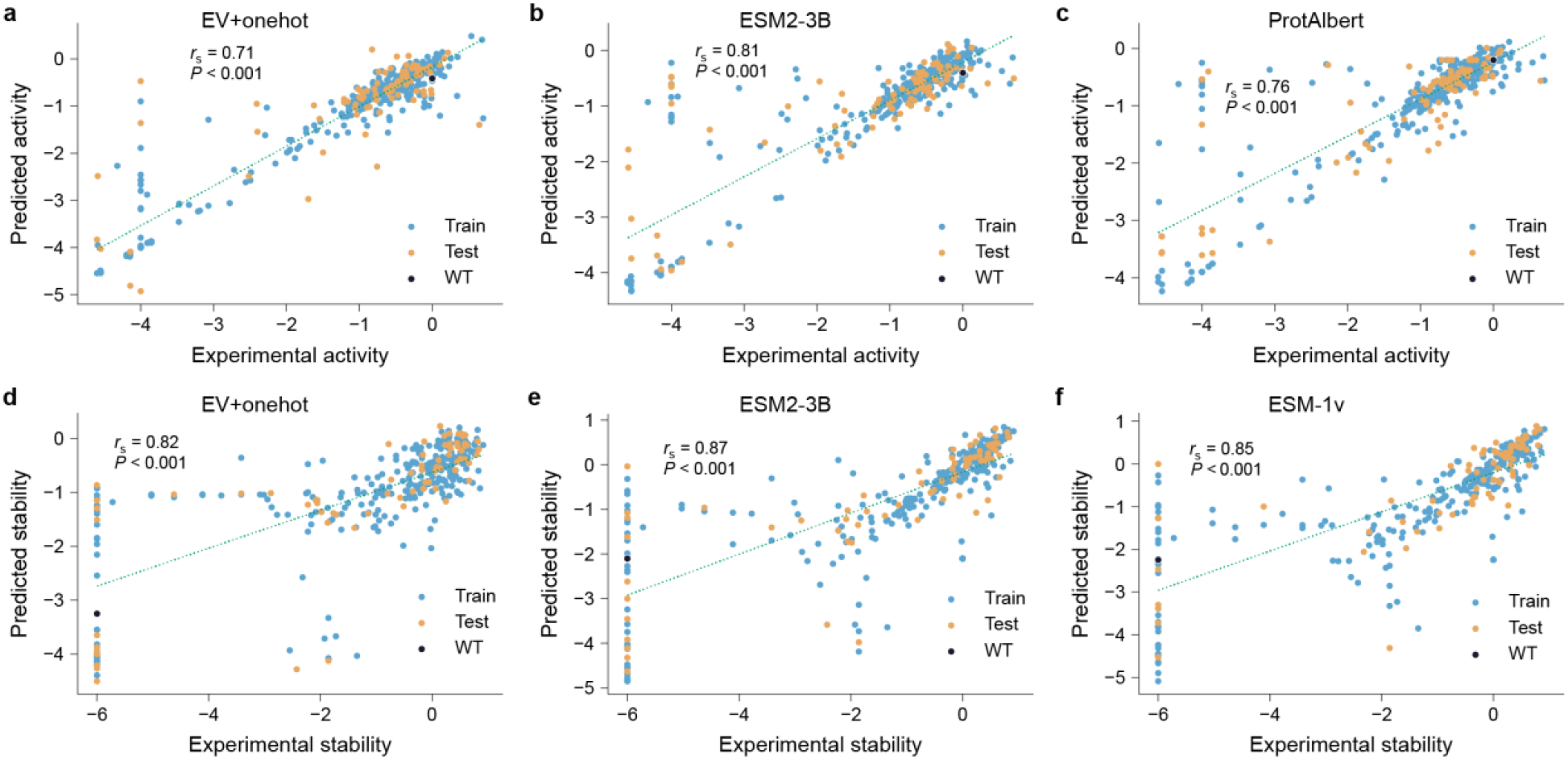
Evaluation of additional models used for variant filtering in round three. **a**–**c**, Evaluation of additional activity models used in variant filtering, Ev+onehot (**a**), ESM-3B (**b**), and ProtAlbert (**c**). **d**–**f**, Evaluation of additional stability models used in variant filtering, Ev+onehot (**d**), ESM-3B (**e**), and ESM-1v(**f**).

**Extended Data Fig. 22.**
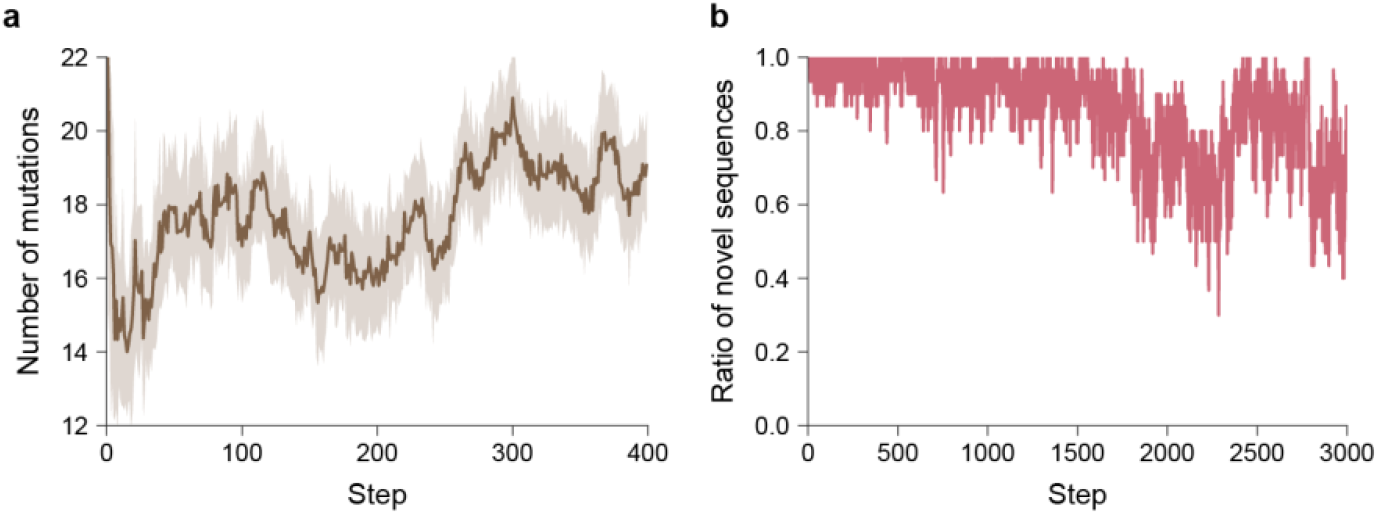
Reinforcement learning dynamics for R3 multi-objective optimization. **a**, Number of mutations during the RL steps. **b**, Ratio of novel sequences in the RL process.

## Extended Data Tables

**Extended Data Table 1.**
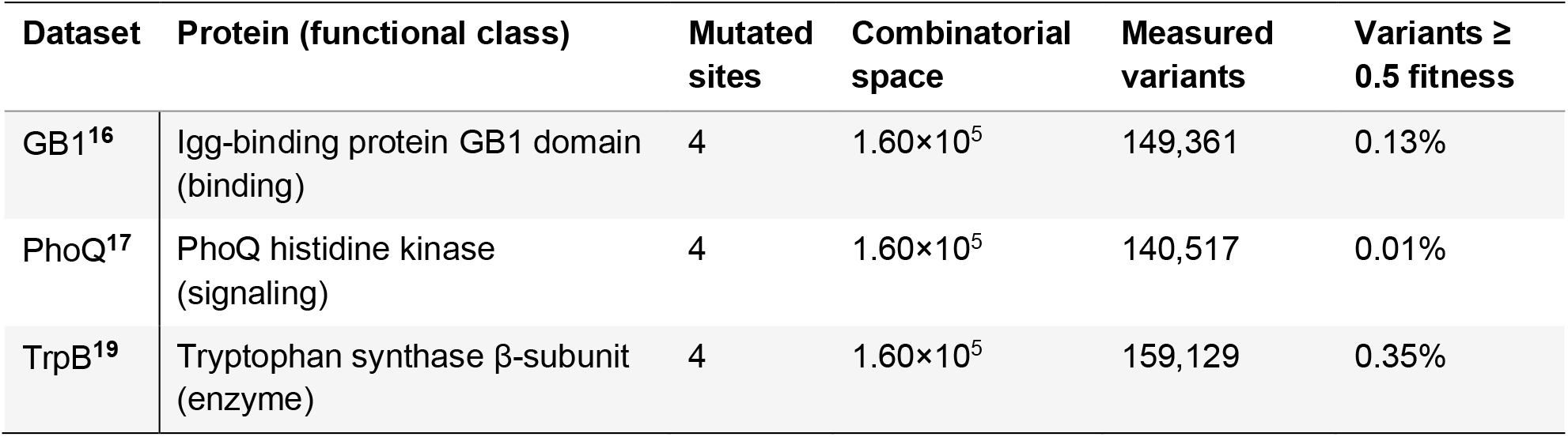
Characteristics of the four-site benchmark datasets.

**Extended Data Table 2.**
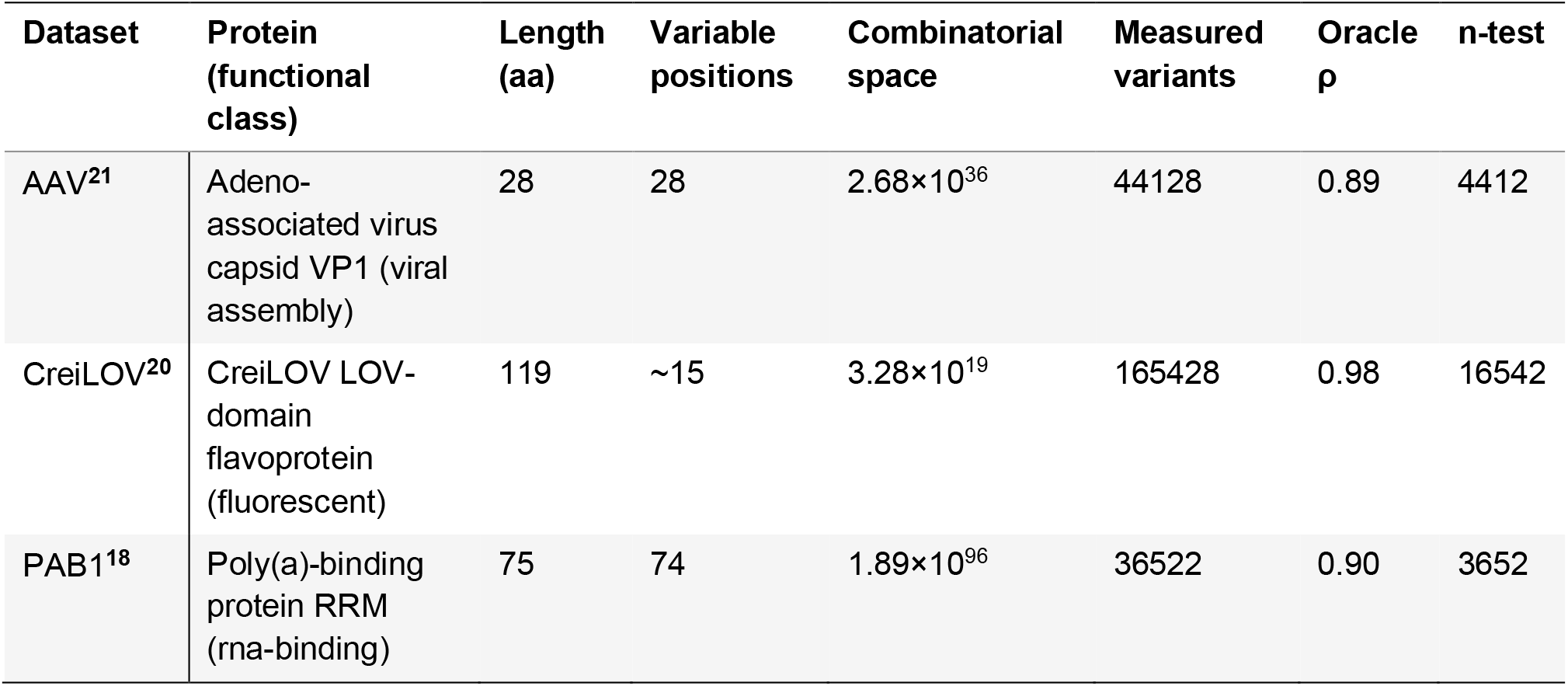
Characteristics of the multi-site benchmark datasets.

**Extended Data Table 3.**
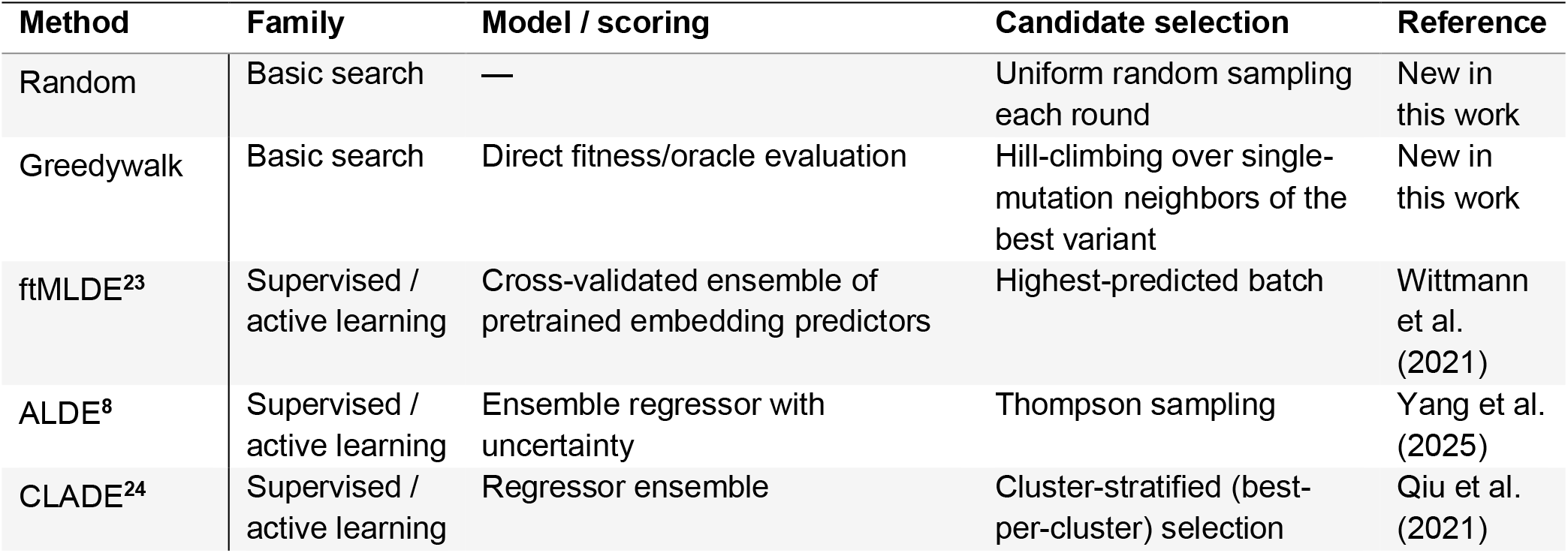
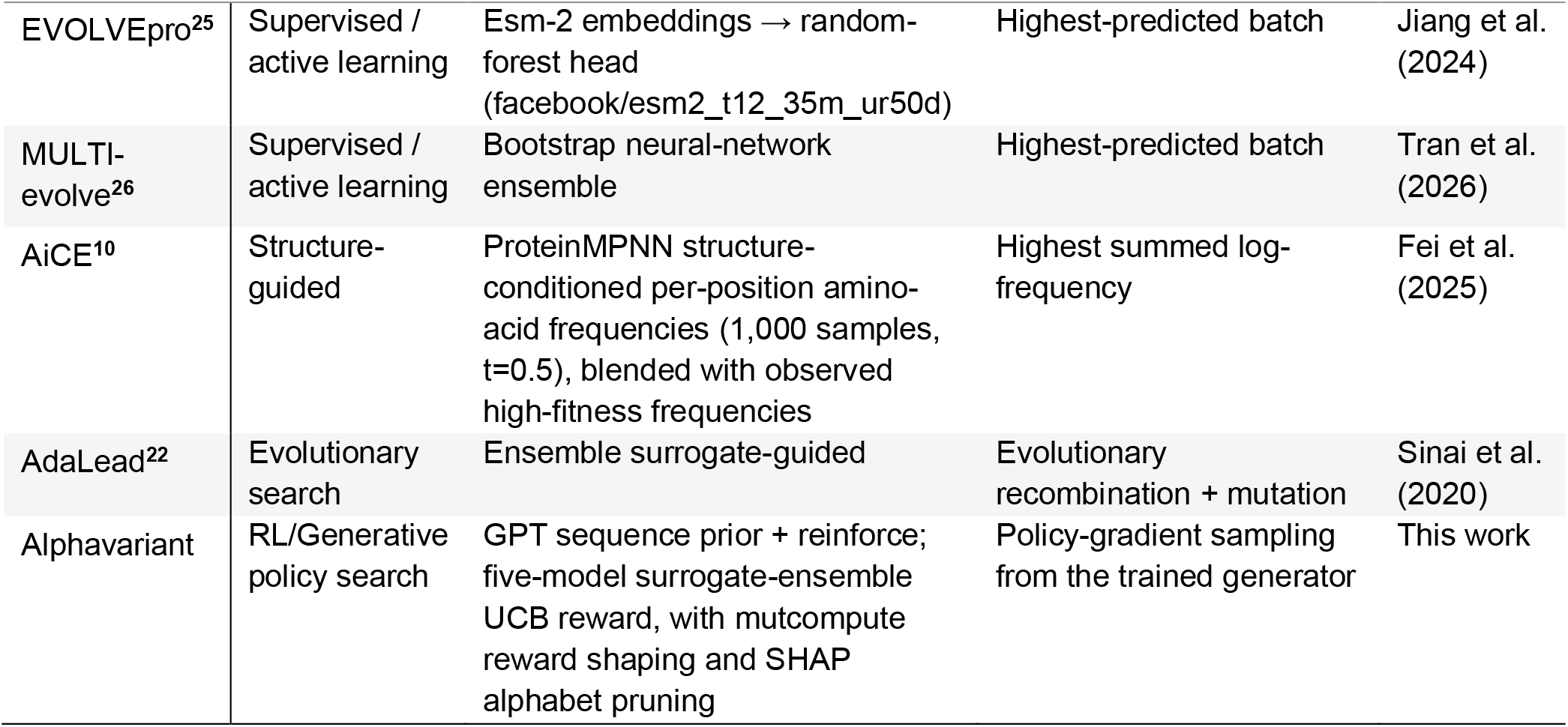
Summary of the ten methods included in the protein fitness optimization benchmark.

**Extended Data Table 4.**
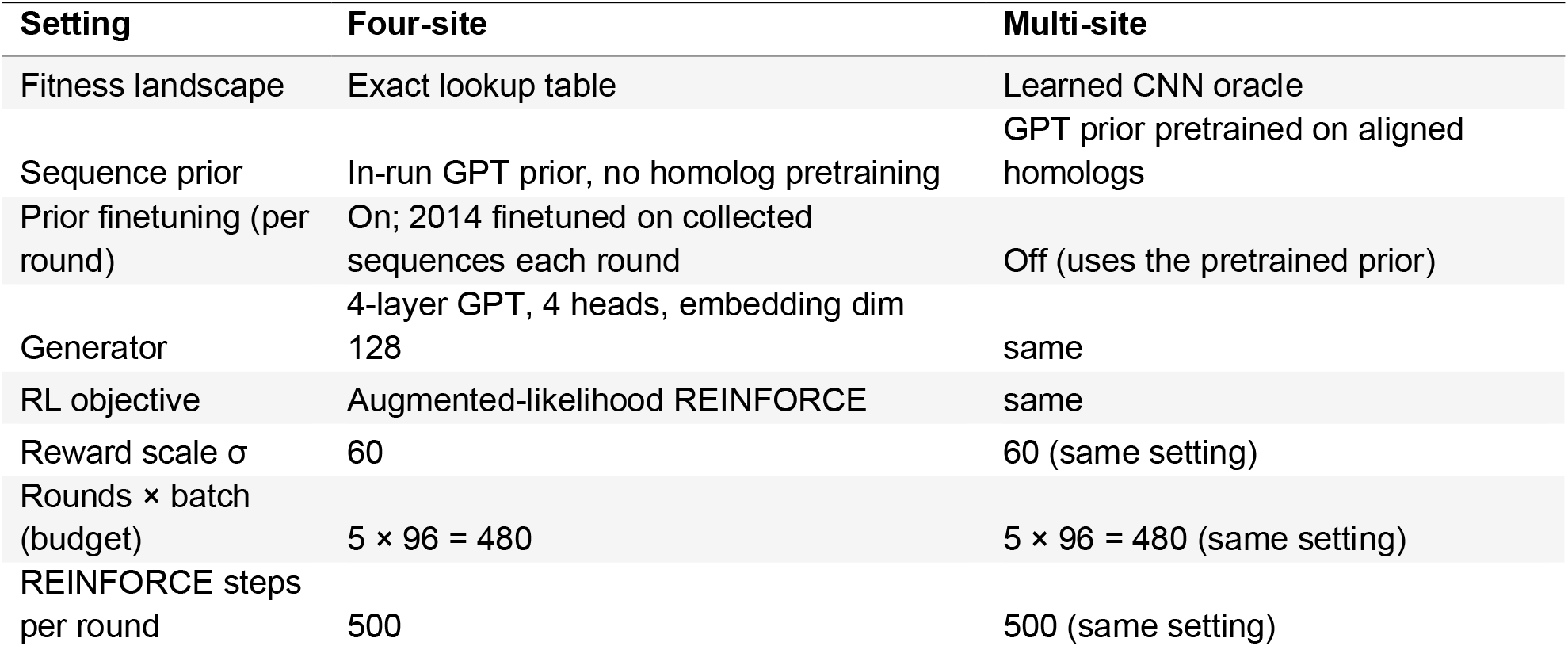
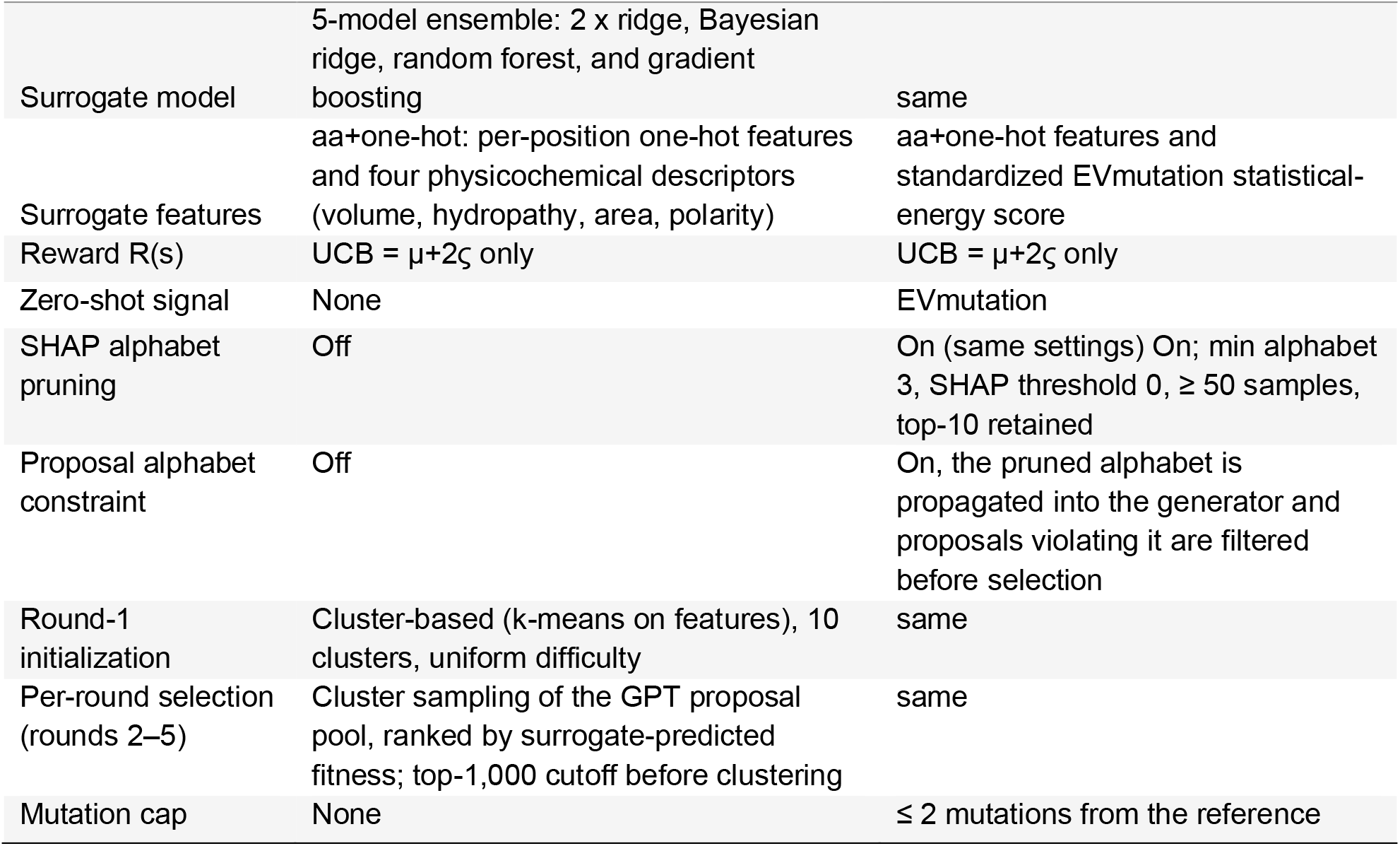
AlphaVariant configurations used in four-site and multi-site benchmark experiments. The table summarizes the benchmark-specific AlphaVariant settings. Shared settings included a four-layer GPT generator with four attention heads and embedding dimension 128, an augmented-likelihood REINFORCE objective, a reward scale of σ = 60, five optimization rounds with 96 sequences per round, 500 REINFORCE steps per round, a five-model surrogate ensemble and SHAP-based alphabet pruning. The four-site and multi-site campaigns differed in the source of fitness values, prior pretraining, surrogate features, reward definition and proposal constraints.

**Extended Data Table 5.**
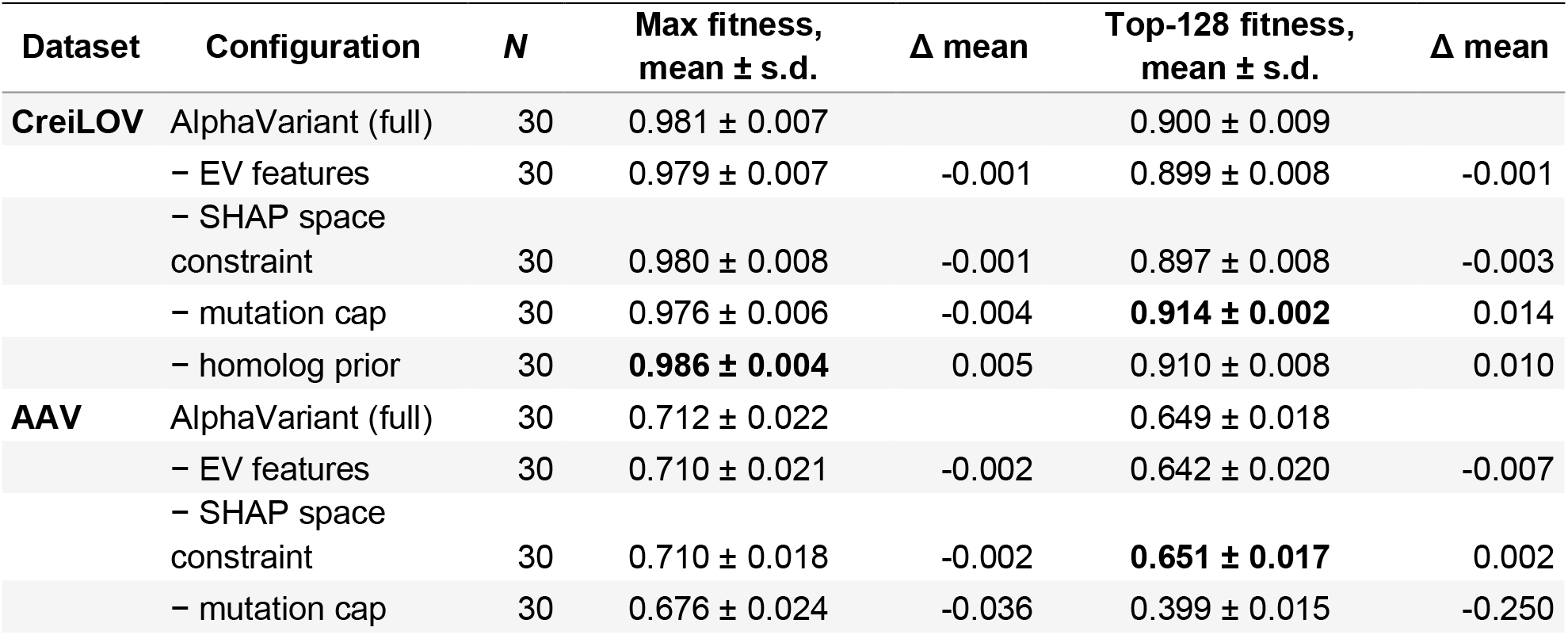
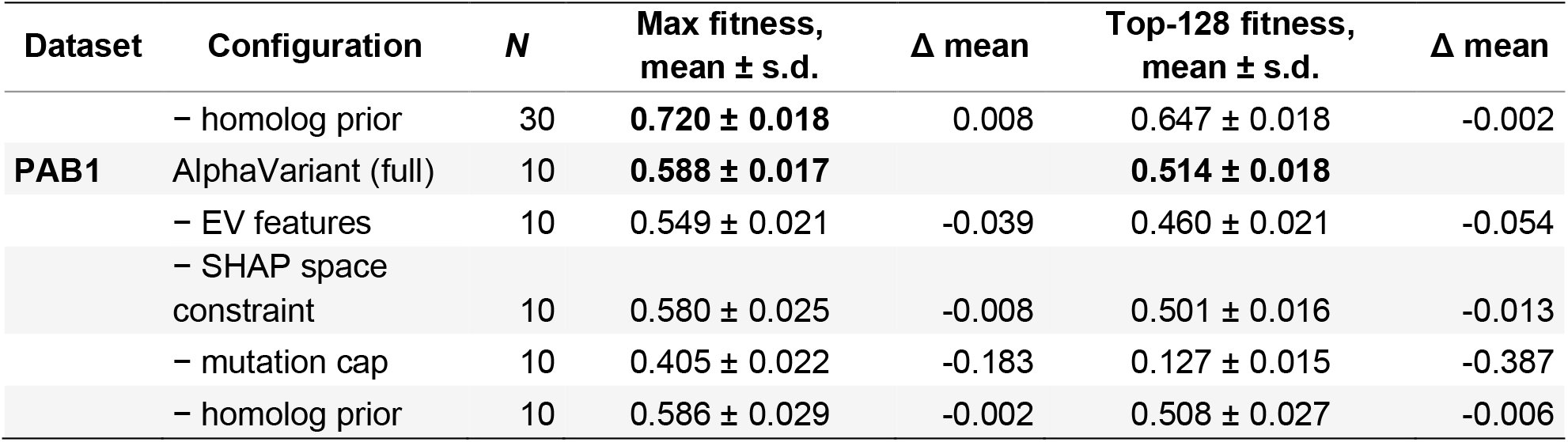
Ablation studies of AlphaVariant on multi-site datasets. Leave-one-out ablation on the three multi-site (learned-oracle) landscapes. Each configuration removes one component from the shipped AlphaVariant configuration and re-runs the full 480-query campaign (96 × 5 rounds). Values are the mean across independent seeds with the standard deviation; Δ is the change in mean versus the full configuration on the same landscape. Fitness is min–max normalized to [0, 1] per landscape (oracle score). Higher is better.

**Extended Data Table 6.**
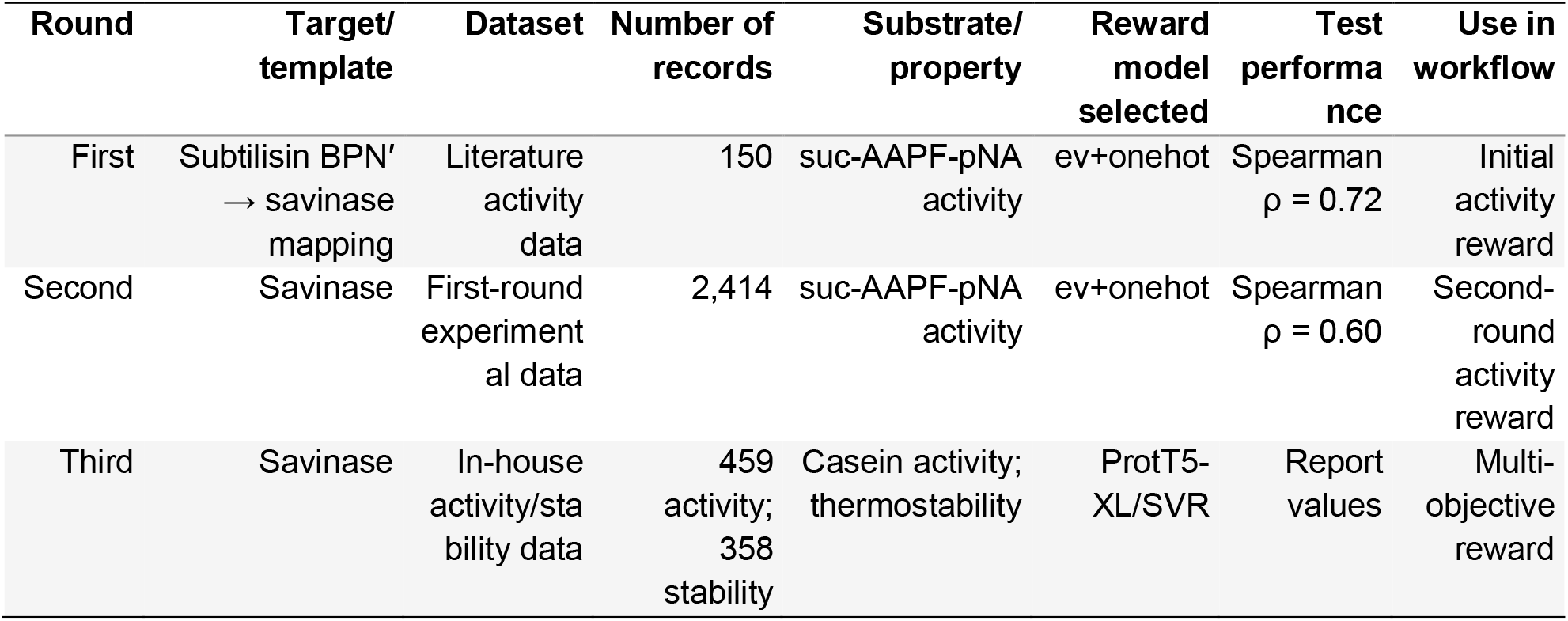
Per-round dataset summary in savinase optimization.

**Extended Data Table 7.**
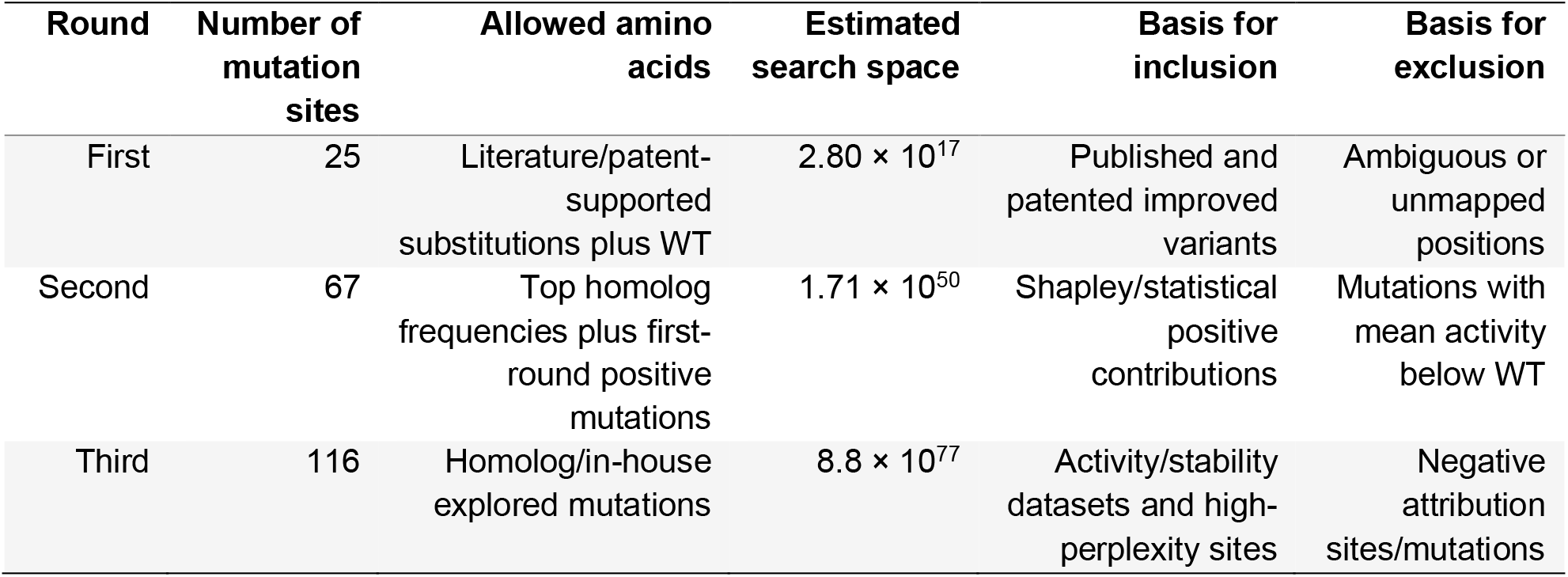
Per-round mutational-space summary in savinase optimization.

