## Supplementary information of AlphaVariant for "Reinforcement learning with evolutionary priors unlocks multi-site enzyme optimization from sparse experimental feedback"

### Supplementary Note 1: Benchmarked methods and configurations

This section details the configurations for the benchmarked methods. The configurations were following the original method implementations. Throughout this study, two distinct dataset types were utilized. For the **four-site** combinatorial landscapes, each method was executed using its native implementation (run_generic.py). Conversely, for the **multi-site** learned-oracle landscapes, all methods were unified under a single generative-proposal script (run_oracle_benchmark.py). In this unified framework, library-enumerating methods were adapted into a mutate-from-elites generator, ensuring that every method operated within an identical, non-enumerable sequence space. Unless otherwise specified, all methods adhered to a standardized protocol consisting of an initial random batch of 96 sequences, followed by four model-guided rounds of 96 sequences each, resulting in a total budget of 480 queries. Each method was evaluated across 30 random seeds. Fitness evaluations for the four-site landscapes relied on exact lookups, whereas the multi-site landscapes utilized trained convolutional neural network (CNN) oracles.

#### Basic search methods

**Random.** This method employed uniform sampling without the use of a predictive model, serving as a lower-bound control. For the four-site landscape, 96 indices were sampled without replacement from the enumerated $20^4$ library in each round. For the multi-site landscape, 96 novel random sequences were generated per round.

**GreedyWalk.** This is a surrogate-free hill-climbing approach. In each round, the highest-fitness sequence identified thus far was selected, and its single-mutation neighbors were directly evaluated. For the four-site landscape, all possible single-mutant neighbors were enumerated from the library, ranked by their measured fitness, and the top 96 were selected, with any remaining slots filled randomly. For the multi-site landscape, single-mutant neighbors were generated and scored directly by the oracle. Both landscapes utilized the standard 20 amino acid alphabet.

### Supervised and active learning methods

All supervised methods encoded sequences using a per-position one-hot representation for the variable positions. Selections were made from either the fully enumerated library (four-site) or a mutate-from-elites candidate pool (multi-site). The multi-site candidate pool was maintained at a size of 8,000, derived from 24 elites, allowing up to two mutations per step.

**ftMLDE**[1]. This method utilized an ensemble regressor coupled with greedy top-N selection. In four-site datasets, a five-model MLDE ensemble was employed, comprising XGBoost (100 trees, max_depth 4, learning rate 0.1), XGBoost with a linear booster (gblinear), random forest (100 trees, max_depth 10), gradient boosting (100 trees, max_depth 3), and kernel ridge regression (alpha = 1.0). Predictions from all models were averaged, and the top 96 variants by predicted fitness over the full library were selected. In multi-site datasets, a "zoo" of four regressors was used: ridge regression (alpha = 1.0), k-nearest neighbors ($k \leq 5$), random forest (100 trees), and extra-trees (100 trees). These models were evaluated using cross-validation ($\leq 3$ folds). The top three models based on cross-validation performance were retained and their predictions averaged. The top 96 variants by predicted fitness from the candidate pool were then selected.
**ALDE**[2]. This is an ensemble regressor incorporating Thompson sampling with an exploration parameter $\xi = 4$. In four-site datasets, a deep neural network (DNN) ensemble (DNN_ENSEMBLE) featuring two hidden layers of 30 units each and leaky-ReLU activations was trained for 300 iterations with a learning rate of $1 \times 10^{-3}$ using one-hot features. Acquisition was performed via Thompson sampling over the full library. In multi-site datasets, a random forest surrogate comprising 100 trees was utilized. Thompson sampling was applied over the candidate pool using the predictive mean and the per-tree standard deviation ($\mu + \sigma \cdot \mathcal{N}(0,1)$).
**CLADE**[3]. This is an ensemble regressor that employs cluster-stratified selection. In four-site datasets, a four-model ensemble consisting of XGBoost, random forest, gradient boosting, and kernel ridge regression was averaged. The library was partitioned using k-means clustering (k = 10), and the variant with the highest predicted fitness from each cluster was selected. In multi-site datasets, a three-model ensemble comprising ridge

regression, random forest, and k-nearest neighbors was averaged. The candidate pool was partitioned using MiniBatch K-means, where $\mathrm{k}$ equaled the batch size. The sequence with the highest predicted fitness within each cluster was selected.

**MULTI-evolve**[4]. This method employs a bootstrap fully-connected neural network (FCN) ensemble with greedy top-N selection. In four-site datasets, an FCN ensemble (2 layers, 100 units, Adam optimizer, batch size 32) was trained on one-hot mutation encodings. The top 96 variants by predicted fitness over the library were selected. In multi-site datasets, a three-network bootstrap FCN ensemble ($256 \rightarrow 64$units, ReLU activations, Adam optimizer with learning rate $1 \times 10^{-3}$, weight decay $1 \times 10^{-4}$, 120 epochs) was used, with each network trained on a bootstrap resample of the data. The top 96 variants by mean prediction over the candidate pool were selected.

**EVOLVEpro**[5]. This approach leveraged frozen protein language model embeddings paired with a random forest top layer. Sequences were embedded using the ESM-2 model (esm2_t12_35M_UR50D, mean-pooled, 480 dimensions). A random forest (100 trees for four-site; 200 trees for multi-site) was trained on the queried labels, and the top 96 variants by predicted fitness were selected. For the multi-site landscape, the candidate pool was restricted to 2,000 sequences per round to manage the computational cost associated with ESM-2 embeddings.

#### Structure-guided design

**AiCE**[6]. This method relied on ProteinMPNN-conditioned per-position amino acid frequencies without a learned regressor. Structure-conditioned frequencies were derived from ProteinMPNN (1,000 sequence samples at a temperature of 0.5) and subsequently blended with the per-position frequencies of the highest-fitness observed variants. The weight assigned to the observed frequencies increased progressively across rounds (from approximately 0.2 to 0.7). Candidate sequences were generated by sampling mutations at variable positions from this blended distribution and were ranked by their summed log-frequency. For the four-site landscape, a linkage-disequilibrium bonus was incorporated for multiple mutations. Dataset-specific structures and precomputed ProteinMPNN frequency tables were utilized.

### Evolutionary search-based method

**AdaLead**[7]. This is a model-guided evolutionary search utilizing recombination and mutation of high-fitness parents. In four-site datasets, the FLEXS AdaLead implementation was used with a five-model CNN ensemble surrogate (32 filters, hidden layer size 100, kernel size 3, trained for 20 epochs each). Mutations were restricted to library-aware positions and amino acids (i.e., only those observed in the library). The top 96 variants by surrogate score were selected, with a maximum of 2,000 model queries permitted per round. In multi-site datasets, a two-wave recombination strategy (20% recombination rate) was combined with mutation from elite parents. Candidates were scored using a CLADE-style three-model ensemble, and the top 96 variants by predicted fitness were selected.

### RL and generative policy-based search

**AlphaVariant (this work).** A GPT sequence prior optimized via augmented-likelihood REINFORCE against a five-model surrogate ensemble. The comprehensive configuration, encompassing the four-site and multi-site oracle settings, is detailed in the main Methods section and the accompanying AlphaVariant configuration table. In brief, the method employed a 4-layer GPT generator and a UCB surrogate reward ($\sigma = 60$). Round 1 initialization was cluster-based, and subsequent rounds utilized a CLADE-style cluster selection for the proposal pool. The four-site configuration utilized an in-run prior that was finetuned each round on the collected sequences (using amino acid and one-hot surrogate features). The multi-site configuration employed a homolog-pretrained prior, EVmutation-augmented surrogate features, a maximum of two mutations per sequence, and a SHAP-pruned proposal alphabet.

### Benchmark evaluation

**Shared settings**. All evaluated methods adhered to a batch size of 96 across five rounds (totaling 480 queries) and utilized a unified metric pipeline. Fitness values were min–max normalized. Performance metrics are reported as medians over 30 independent seeds, with statistical significance determined using paired Bonferroni-corrected Wilcoxon tests (detailed in the main Methods).

**Configuration adherence**. Each method was executed using the configuration recommended by its original authors, with adaptations introduced solely as necessary to accommodate the multi-site generative-proposal framework described above.

### Supplementary Note 2: Ablation of AlphaVariant components

To quantify the contribution of each AlphaVariant component, we performed leave-one-out ablations on the benchmark landscapes, re-running the full optimization campaign (480 queries; 96 × 5 rounds) for each configuration. Both the four-site ablations and the multi-site ablations used the shipped full configuration as the baseline. Each configuration was run for 30 independent seeds, except multi-site PAB1 (10 seeds), where per-seed cost on the longer sequences made 30 seeds impractical. Δ denotes the change in median normalized maximum fitness or top-128 fitness relative to the baseline; per-component values are in the **Extended Data Table 5** and **Supplementary Table 1**.

#### Four-site landscapes

Removing per-round prior finetuning reduced top-128 fitness by 0.004–0.044 on PhoQ, GB1 and TrpB datasets, and GB1 maximum fitness by 0.008, while not drastically improving any landscape, so finetuning was retained. The reinforce step was contributing is contributing to top-128 fitness (removing it lowers top-128 fitness by 0.005–0.041 on PhoQ/GB1/TrpB and GB1 max by 0.02). Replacing the five-model surrogate ensemble with a single random forest mainly reduced top-128 fitness (PhoQ, TrpB) with a landscape-dependent effect. The structure-based reward shaping (MutCompute) and SHAP-based alphabet pruning did not improve over the current configuration on any four-site landscape and were therefore excluded from the settings.

#### Multi-site landscapes

The component ablation for the multi-site configuration — in which the mutation cap is the dominant component and its effect on batch quality scales with the size of the variable region, the EV-augmented features contribute on the larger landscapes, and SHAP pruning and the homolog-pretrained prior are approximately neutral — is presented in the main-text results.

***Per-round finetuning on multi-site***

Because per-round prior finetuning helped on four-site, we separately tested whether it also helped on the multi-site landscapes, where the prior is already homolog-pretrained. Adding finetuning (paired against the shipped configuration on the same seeds) was marginally positive on AAV ($\Delta \approx +0.009$ on both metrics) and essentially neutral on CreiLOV ($\Delta \approx +0.001$–$0.002$), but negative on PAB1 ($\Delta$ max $-0.014$). Because finetuning did not consistently help on multi-site, it was not adopted there and is used only on four-site, where the prior is initialized in-run.

**Summary of ablation studies**

The two benchmark regimes are governed by different load-bearing components: four-site performance rests on prior finetuning and the policy-gradient step, whereas multi-site performance rests on the manifold-constraining mutation cap. AlphaVariant's robustness across regimes therefore arises from applying the appropriate constraint where the landscape demands it. A landscape specific configuration can further improve the performance of AlphaVariant, but aiming for a critical benchmark, we didn't implement it on our work. This finding also highlights the heterogenicity of protein fitness landscapes, and landscape-specific implementation will be useful in real world engineering projects.

### Supplementary Note 3: IP conflict matching

**Variant representation and residue numbering**

Each candidate variant was first converted into a set of point substitutions relative to the wild-type savinase. To keep residue positions consistent with the numbering used across all analyzed patents, the wild-type reference sequence was stored in a positionally aligned form in which gap characters mark reference positions that are absent from the wild type.

A variant was scored only if it was full-length (269 residues) and contained no ambiguous or unknown residues. The variant was then compared residue-by-residue against the aligned wild type, skipping gap positions when reading the variant residue but assigning each position its index in the unified (alignment-based) coordinate system. This produced, for each variant, a mutation string such as ‘A21V/N76D/.../L262E’, in which every token encodes a substitution at a unified position.

Importantly, matching is indexed only on the (position, substituted residue) pair; the identity of the original wild-type residue in a token is treated as informational, so a patent token written with a wild-card source residue (e.g., ‘X61E’ meaning "any-to-E at position 61") matches a variant carrying that substitution regardless of the source residue.

**Patent rule definition**

The matching rules from 71 unique patents were retrieved. For each patent, the relevant claims were read manually and translated into one or more machine-readable matching rules expressed in a compact, regex-like grammar (summarized in **Supplementary Table 2**), and every rule was verified by an IP expert.

The grammar encodes the logical structure of patent claims: ‘&’ denotes conjunction (all listed substitutions must co-occur in the variant); ‘;’ separates independent claimed variants of the same patent (a top-level disjunction); ‘{…}’ lists the alternative target residues allowed at a single position (a position-level disjunction, e.g., ‘N261{D,C,E,L,M,R,V,W,Y,F}’); ‘^’ encodes a negation proviso (the variant must not carry the indicated substitution); a single parenthesized group combined by ‘&’ with a set of fixed substitutions captures the common claim pattern "the listed mutations and a further substitution selected from the group" (e.g., ‘X101E&(S3T|R45{E,D,Q}|P55N|…)’); the prefix ‘n+’ inside a group requires at least n of the listed alternatives to be present simultaneously; bracketed alternatives ‘[…]’ allow a multi-substitution combination to appear as a single option within a disjunction (e.g., ‘…|[N238E&L262E]|…’); and insertions and deletions are supported through dedicated tokens (e.g., ‘G97GN’ for an insertion and ‘X131-‘ for a deletion).

**Matching procedure**

At evaluation time, each patent rule was first expanded into disjunctive normal form: meta-rules were split on ‘;’, and every grouped, ‘n+’, ‘|’, or ‘{…}’ construct was enumerated into the full set of equivalent pure-conjunctive (AND-only) sub-rules.

A variant was deemed to conflict with a conjunctive sub-rule if its mutation set contained all of the required (position, residue) substitutions and none of the negated ones;

additional substitutions beyond those claimed did not prevent a match, so the rule recognizes any variant that subsumes a claimed combination. A variant was flagged as an IP conflict if it satisfied at least one expanded sub-rule of at least one patent, and was otherwise considered IP-free.

### Supplementary Tables

#### Supplementary Table 1 | Ablation studies of AlphaVariant on four-site datasets.

Leave-one-out ablation on the three four-site (combinatorial lookup) landscapes. Each configuration removes one component from the default configuration and re-runs the full 480-query campaign (96 × 5 rounds); n = 30 seeds per configuration. Fitness is min–max normalized to [0, 1] per landscape; Δ is the change in the summary statistic versus the full configuration on the same landscape.

| Dataset | Configuration | N | Max fitness, mean ± s.d. | Δ mean | Top-128 fitness, mean ± s.d. | Δ mean |
|---|---|---|---|---|---|---|
| GB1 | AlphaVariant (bare + finetune) | 30 | 0.896 ± 0.126 | | 0.454 ± 0.063 | |
| | − finetune prior | 30 | 0.889 ± 0.129 | -0.008 | 0.423 ± 0.064 | -0.031 |
| | − ensemble scoring (single RF) | 30 | **0.943 ± 0.107** | 0.047 | **0.462 ± 0.052** | 0.008 |
| | − RL step (generate + prioritize) | 30 | 0.876 ± 0.113 | -0.02 | 0.449 ± 0.043 | -0.005 |
| PhoQ | AlphaVariant (bare + finetune) | 30 | 0.484 ± 0.220 | | **0.116 ± 0.020** | |
| | − finetune prior | 30 | 0.485 ± 0.201 | 0.001 | 0.112 ± 0.022 | -0.004 |
| | − ensemble scoring (single RF) | 30 | 0.502 ± 0.237 | 0.018 | 0.110 ± 0.018 | -0.006 |
| | − RL step (generate + prioritize) | 30 | **0.525 ± 0.210** | 0.042 | 0.109 ± 0.020 | -0.007 |
| TrpB | AlphaVariant (bare + finetune) | 30 | 0.871 ± 0.083 | | **0.579 ± 0.046** | |
| | − finetune prior | 30 | 0.867 ± 0.094 | -0.005 | 0.535 ± 0.085 | -0.044 |
| | − ensemble scoring (single RF) | 30 | 0.889 ± 0.098 | 0.017 | 0.525 ± 0.066 | -0.054 |
| | − RL step (generate + prioritize) | 30 | **0.891 ± 0.096** | 0.019 | 0.538 ± 0.054 | -0.041 |

#### Supplementary Table 2 | The grammar of patent matching rules.

| Operator | Meaning | Example |
|---|---|---|
| <src><pos><tgt> | Substitution token; matched on (pos,tgt) only, src informational (X = wild-type) | X61E, N76D |

| Operator | Meaning | Example |
|---|---|---|
| & | Conjunction — all substitutions must co-occur | X61E&X194P&X261D |
| ; | Separates independent claimed variants (top-level OR) | …S130A;I72V&…&S164I |
| {…} | Alternative target residues at one position (OR) | X9{C,D,E,Q} |
| ^ | Negation proviso (must not be present) | X9R&X19L&X62D&^X14H |
| (…\|…) | Single grouped OR alternatives combined with fixed mutations | X101E&(S3T\|R45{E,D,Q}\|P55N) |
| n+ | At least n of the listed alternatives required | N76D&(2+S259D\|T260{A,E,P}\|…) |
| […] | Multi-substitution combination as one alternative inside an OR | …\|[N238E&L262E]\|… |
| insertion / deletion | Insertion G97GN; deletion X131- | X99D&(1+G97GN\|A98AA\|…) |

**Supplementary Table 3 | Primers for constructing fixed site savinase variants in the first round.**

| Primer | Sequence (5' → 3') |
|---|---|
| L1-203,212-F | GGTTCATACTATGCCAGCTTAAGCGGTACATCGATGGCTAC |
| L1-203,212-R | TACCGCTTAAGCTGGCATAGTATGAACCTGGGTATGTGCTC |
| L1-250-F | GCAACGAACTTAGGAAGCACGAACTTGTATGG |
| L1-250-R | CGTGCTTCCTAAGTTCGTTGCCGTATTCTTTAG |
| L2-S3T-F | AATGGCGCAAACAGTGCCATGGGGAATTAGCC |
| L2-S3T-R | GCACTGTTTGCGCCATTGTCGTTACTTCTGCA |
| L2-212,218-F | AGCTTAAGCGGTACATCGATGGCTGCTCCTCATGTTGCAGGTGCAGCAG |
| L2-212,218-R | GATGTACCGCTTAAGCTGGCATACGCTGAACCTGGGTATG |
| L3-96-F | GTTAAAGTATTAGGGTCGAGCGGTTCAGGTTCGG |
| L3-96-R | CCGAACCTGAACCGCTCGACCCTAATACTTTAAC |
| L3-207-F | CAGGTTCAAGGTATGCCAGCTTAAG |
| L3-207-R | CATACCTTGAACCTGGGTATGTGCTC |
| L4-207-F | GGTTCAGAGTATGCCAGCTTAAGC |
| L4-207-R | GGCATACTCTGAACCTGGGTATG |
| L4-250-F | ATACGGCAACGAACTTAGGAAGCACGAAC |
| L4-250-R | TTCGTGCTTCCTAAGTTCGTTGCCGTAT |

**Supplementary Table 4 | Primers for constructing the libraries in the first round.**

| Primer | Sequence (5' → 3') |
|---|---|
| L1-sav1-1 | GAAGTAACGACAATGGCGCAARCCGTGCCATGGGGAATTAGCCGTGTGCAAGCCCCAGC |
| L1-sav1-2 | GAAGTAACGACAATGGCGCAAWGCGTGCCATGGGGAATTAGCCGTGTGCAAGCCCCAGC |
| L1-sav1-3 | GAAGTAACGACAATGGCGCAAGTGGTGCCATGGGGAATTAGCCGTGTGCAAGCCCCAGC |
| L1-sav2 | AGCAACTTTTACACCAGAACCTGTCAATCCACGGTTATGGGCAGCTGGGGCTTGCACAC |
| L1-sav3 | CAGGTTCTGGTGTAAAAGTTGCTGTCCTCGATACAGGTATTTCCACTCATCCAGACTTA |
| L1-sav4-1 | CCCTGGTACAAAGCTAGCGCCACCACGAATKTTTAAGTCTGGATGAGTGGAAATACCTG |
| L1-sav4-2 | CCCTGGTACAAAGCTAGCGCCACCACGAATCTCTAAGTCTGGATGAGTGGAAATACCTG |
| L1-sav4-3 | CCCTGGTACAAAGCTAGCGCCACCACGAATACGTAAGTCTGGATGAGTGGAAATACCTG |
| L1-sav5 | CGCTAGCTTTGTACCAGGGGAACCATCCACTCAAGATGGGAATGGGCATGGCACGCATG |
| L1-sav6 | AGAACGCCAATCGAATTGTTTAAAGCAGCAATCGTCCCGGCCACATGCGTGCCATGCCC |
| L1-sav7 | CTTTAAACAATTCGATTGGCGTTCTTGGCGTAGCGCCGAGCGCGGAACTATACGCTGTT |
| L1-sav8-1 | TCGAGCTGACCGAACCTGAACCGCTGCYCCCTAATACTTTAACAGCGTATAGTTCCGCG |
| L1-sav8-2 | TCGAGCTGACCGAACCTGAACCGCTTTTCCCTAATACTTTAACAGCGTATAGTTCCGCG |
| L1-sav8-3 | TCGAGCTGACCGAACCTGAACCGCTCGCCCCTAATACTTTAACAGCGTATAGTTCCGCG |
| L1-sav9 | GTGCATGCCATTGTTCCCTGCCCATTCCAATCCTTGGGCAATCGAGCTGACCGAACCTG |
| L1-sav10 | GGCACTGCYCGAAGGGCTTCCTAAACTCAAATTAGCAACGTGCATGCCATTGTTCCCTG |
| L1-sav11 | CCCTTCGRGCAGTGCCACACTTGAGCAAGCTGTTAATAGCGCGACTTCTAGAGGGGTTC |

| Primer | Sequence (5' → 3') |
|---|---|
| L1-sav12 | TTGAGCCTGCACCTGAATTCCCAGATGCCGCTACAACAAGAACCCCTCTAGAAGTCGCG |
| L1-sav13 | GAATTCAGGTGCAGGCTCAATCAGCTATCCGGCCCGTTATGCGAACGCAATGGCAGTCG |
| L1-sav14 | CTGTGAAAAGCTGGCGCGGTTGTTGTTTTGGTCAGTCGCTCCGACTGCCATTGCGTTCG |
| L1-sav15 | CGCGCCAGCTTTTCACAGTATGGCGCAGGGCTTGACATTGTCGCACCAGGTGTAAACGT |
| L1-sav16 | CGCTTAAGCTGGCATAATATGAACCTGGWAATGTGCTCWGCACGTTTACACCTGGTGCG |
| L1-sav17-1 | GGTTCATATTATGCCAGCTTAAGCGGTACATCGATGKCCTSTCCTCATGTTGCAGGTGC |
| L1-sav17-2 | GGTTCATATTATGCCAGCTTAAGCGGTACATCGATGKCCGCGCCTCATGTTGCAGGTGC |
| L1-sav18 | ACATTGGACCAAGATGGGTTCTTTTGTTTAACAAGGGCTGCTGCACCTGCAACATGAGG |
| L1-sav19 | AGAACCCATCTTGGTCCAATGTACAAATCCGCAATCATCTAAAGAATACGGCAACGAAC |
| L1-sav20 | CAAGTCCGCTTCCATACAAGTTCGTGCTTCCTAAGTTCGTTGCCGTATTCTTTAGATGA |
| L1-sav21 | AACTTGTATGGAAGCGGACTTGTCAATGCAGAAGCGGCAACACGCTAACTTCAGCACAA |
| L1-sav22 | TCGTGTCTTTCTTGGAATTGTGCTGAAGTTAGCGTGTTGCC |
| L2-sav1 | GATGCAGAAGTAACGACAATGGCGCAAACAGTGCCATGGGGAATTAGCCGTGTGCAAGC |
| L2-sav2 | TACACCGCWACCTGTCAATCCACGGTTATGGGCAGCTGGGGCTTGCACACGGCTAATTC |
| L2-sav3 | GGATTGACAGGTWGCGGTGTAAAAGTTGCTGTCCTCGATACAGGTATTTCCACTCATCC |
| L2-sav4-1 | GGTACAAAGCTAGCGCCACCACGAATWTTTAAGTCTGGATGAGTGGAAATACCTGTATC |
| L2-sav4-2 | GGTACAAAGCTAGCGCCACCACGAATACGTAAGTCTGGATGAGTGGAAATACCTGTATC |
| L2-sav5 | GGCGCTAGCTTTGTACCAGGGGAACCATCCACTCAAGATGGGAATGGGCATGGCACGCA |

| Primer | Sequence (5' → 3') |
|---|---|
| L2-sav6 | ACGCCAATCGAATTGTTTAAAGCAGCAATCGTCCCGGCCACATGCGTGCCATGCCCATT |
| L2-sav7 | GCTTTAAACAATTCGATTGGCGTTCTTGGCGTAGCGCCGAGCGCGGAACTATACGCTGT |
| L2-sav8 | TCGAGCTGACCGAACCTGAACCGCTGGMCCCTAATACTTTAACAGCGTATAGTTCCGCG |
| L2-sav9 | AGGTTCGGTCAGCTCGATTGCCCAAGGATTGGAATGGGCAGGGAACAATGGCATGCACG |
| L2-sav10-1 | TGTGGCACTCGTCGAAGGGCTTCCTAAACTCAAATTAGCAACGTGCATGCCATTGTTCC |
| L2-sav10-2 | TGTGGCACTAYCCGAAGGGCTTCCTAAACTCAAATTAGCAACGTGCATGCCATTGTTCC |
| L2-sav11-1 | CTTCGACGAGTGCCACACTTGAGCAAGCTGTTAATAGCGCGACTTCTAGAGGGGTTCTT |
| L2-sav11-2 | CTTCGGRTAGTGCCACACTTGAGCAAGCTGTTAATAGCGCGACTTCTAGAGGGGTTCTT |
| L2-sav12 | GATTGAGCCTGCACCTGAATTCCCAGAGGMCGCTACAACAAGAACCCCTCTAGAAGTCG |
| L2-sav13 | CAGGTGCAGGCTCAATCAGCTATCCGGSCCGTTATGCGAACGCAATGGCAGTCGGAGCG |
| L2-sav14 | GCCATACTGTGAAAAGCTGGCGCGGTTGTTGTTTTGGTCAGTCGCTCCGACTGCCATTG |
| L2-sav15 | CCAGCTTTTCACAGTATGGCGCAGGGCTTGACATTGTCGCACCAGGTGTAAACGTGCAG |
| L2-sav16 | GTACCGCTTAAGCTGGCATACGCTGAACCTGGAWGTGTGCTCTGCACGTTTACACCTGG |
| L2-sav17 | GTATGCCAGCTTAAGCGGTACATCGATGGCTGCTCCTCATGTTGCAGGTG |
| L2-sav18 | GGACCAAGATGGGTTCTTTTGTTTAACAAGGGCTGCTGCACCTGCAACATGAGGAG |
| L2-sav19 | AAAGAACCCATCTTGGTCCAATGTACAAATCCGCAATCATCTAAAGAATACG |
| L2-sav20-1 | CGCTTCCATACAAGTTCGTGCTTCCTAATTTCGTTGCCGTATTCTTTAGATGATTGCGG |
| L2-sav20-2 | CGCTTCCATACAAGTTCGTGCTTCCTAAGCCCGTTGCCGTATTCTTTAGATGATTGCGG |
| L2-sav20-3 | CGCTTCCATACAAGTTCGTGCTTCCTAAATWCGTTGCCGTATTCTTTAGATGATTGCGG |

| Primer | Sequence (5' → 3') |
|---|---|
| L2-sav21 | ACGAACTTGTATGGAAGCGGACTTGTCAATGCASAAGCGGCAACACGCTAACTTCAGCA |
| L2-sav22 | TCGTGTCTTTCTTGGAATTGTGCTGAAGTTAGCGTGTTGC |
| L3-sav1-1 | GATGCAGAAGTAACGACAATGGCGCAAASCGTGCCATGGGGAATTAGCCGTGTGCAAGC |
| L3-sav1-2 | GATGCAGAAGTAACGACAATGGCGCAATATGTGCCATGGGGAATTAGCCGTGTGCAAGC |
| L3-sav1-3 | GATGCAGAAGTAACGACAATGGCGCAAKTGGTGCCATGGGGAATTAGCCGTGTGCAAGC |
| L3-sav2 | TACACCAGRACCTGTCAATCCACGGTTATGGGCAGCTGGGGCTTGCACACGGCTAATTC |
| L3-sav3 | GGATTGACAGGTYCTGGTGTAAAAGTTGCTGTCCTCGATACAGGTATTTCCACTCATCC |
| L3-sav6 | ACGCCAATCGAATTATYTAAAGCAGCAATCGTCCCGGCCACATGCGTGCCATGCCCATT |
| L3-sav7 | GCTTTARATAATTCGATTGGCGTTCTTGGCGTAGCGCCGAGCGCGGAACTATACGCTGT |
| L3-sav8 | TCGAGCTGACCGAACCTGAACCGCTCGACCCTAATACTTTAACAGCGTATAGTTCCGCG |
| L3-sav10 | TGTGGCACTTGGCGAAGGGCTTCCTAAACTCAAATTAGCAACGTGCATGCCATTGTTCC |
| L3-sav11 | CTTCGCCAAGTGCCACACTTGAGCAAGCTGTTAATAGCGCGACTTCTAGAGGGGTTCTT |
| L3-sav12 | GATTGAGCCTGCACCTGAATTCCCAGATGCCGCTACAACAAGAACCCCTCTAGAAGTCG |
| L3-sav13 | CAGGTGCAGGCTCAATCAGCTATCCGGSCCGTTATGCGAACGCAATGGCAGTCGGAGC |
| L3-sav15 | CCAGCTTTTCACAGTATGGCGCAGGGCTTGACATTGTCGCACCAGGTGTAAACGTGCHG |
| L3-sav16 | TACCTTGAACCTGGAWGTGTGCTCDGCACGTTTACACCTGGTGCGACAATGTCAAGCCC |
| L4-sav1-2 | GAAGTAACGACAATGGCGCAAGTGGTGCCATGGGGAATTAGCCGTGTGCAAGCCCCAGC |
| L4-sav1-3 | GAAGTAACGACAATGGCGCAATMTGTGCCATGGGGAATTAGCCGTGTGCAAGCCCCAGC |

| Primer | Sequence (5' → 3') |
|---|---|
| L4-sav6 | AGAACGCCAATCGAATTATYTAAAGCAGCAATCGTCCCGGCCACATGCGTGCCATGCCC |
| L4-sav7 | CTTTARATAATTCGATTGGCGTTCTTGGCGTAGCGCCGAGCGCGGAACTATACGCTGTT |
| L4-sav8 | TCGAGCTGACCGAACCTGAACCGCTGGMCCCTAATACTTTAACAGCGTATAGTTCCGCG |
| L4-sav11 | CCCTTCGCCAAGTGCCACACTTGAGCAAGCTGTTAATAGCGCGACTTCTAGAGGCGTTC |
| L4-sav15-1 | CGCGCCAGCTTTTCACAGTATGGCGCAGGGCTTGACATTGTCGCACCAGGTGTAGACGTGCAGAGCACATTWCCAGGTTCAGAGTATGCCAGC |
| L4-sav15-2 | CGCGCCAGCTTTTCACAGTATGGCGCAGGGCTTGACATTGTCGCACCAGGTGTAAGCGTGCAGAGCACATTWCCAGGTTCAGAGTATGCCAGC |
| L4-sav15-3 | CGCGCCAGCTTTTCACAGTATGGCGCAGGGCTTGACATTGTCGCACCAGGTGTAAGCGTGCAGAGCACACATCCAGGTTCAGAGTATGCCAGC |
| L4-sav15-4 | CGCGCCAGCTTTTCACAGTATGGCGCAGGGCTTGACATTGTCGCACCAGGTGTAGACGTGCAGAGCACACATCCAGGTTCAGAGTATGCCAGC |
| L4-sav16-1 | CCTGCAACATGAGGAGCAGCCATCGATGTACCGCTTTTGCTGGCATACTCTGAACCTGG |
| L4-sav16-2 | CCTGCAACATGAGGAGCAGCCATCGATGTACCGCTGBCGCTGGCATACTCTGAACCTGG |
| L4-sav16-3 | CCTGCAACATGAGGAGCAGCCATCGATGTACCGCTATAGCTGGCATACTCTGAACCTGG |
| L4-sav16-4 | CCTGCAACATGAGGAGCAGCCATCGATGTACCGCTCAKGCTGGCATACTCTGAACCTGG |
| L4-sav17 | CTGCTCCTCATGTTGCAGGTGCAGCAGCCCTTGTTAAACAAAAGAACCCATCTTGGTCC |
| L4-sav18 | GCCGTATTCTTTAGATGATTGCGGATTTGTACATTGGACCAAGATGGGTTCTTTTGTTT |
| L4-sav19 | CCGCAATCATCTAAAGAATACGGCAACGAACTTAGGAAGCACGAACTTGTATGGAAGCG |
| L4-sav20 | TCGTGTCTTTCTTGGAATTGTGCTGAAGTTAGCGTGTTGCCGCTTSTGCATTGACAAGTCCGCTTCCATACAAGTTCGTGC |
| MC-PCR-F | GATGCAGAAGTAACGACAATGGC |
| MC-PCR-R | TCGTGTCTTTCTTGGAATTGTGC |
| pWB980(-mp)-F | AGCACAATTCCAAGAAAGACACGATTT |
| pWB980(-mp)-R | CCATTGTCGTTACTTCTGCATCCTCTT |

| Primer | Sequence (5' → 3') |
|---|---|
| Sav-ep-pcr-F | CGCTTGAACTCGATCCAGCGATTTCTTATATTGAAGAGGATGCAGAAGTAACGACAATG |
| Sav-ep-pcr-R | GTTATGAGTTAGTTCAAATTCGTTCTTTTTAGGTTCTTCTTGGAATTGTGCTGAAGTTA |
| 980-ep-pcr-f | GAACTAACTCATAACCGAGAGG |
| 980-ep-pcr-R | GCTGGATCGAGTTCAAGCG |
| Sav-SSM-F | GTTCAACGTATGCCAGCNNKNNKGGTACATCGNNKGCTACTCCTCATGTTG |
| Sav-SSM-R | GCTGGCATACGTTGAACCTGGGTATGTGCT |

**Supplementary Table 5 | Primers for constructing fixed-site variants in the second round.**

| Primer | Sequence (5' → 3') |
|---|---|
| R2-FP-1 | GTTAAAGTATTAGGGAGCAACGGTTCAGGTTCGGTCAGCTCGATTGCCCAAGGATTGGAATGGGCAGGGAACAATGGCATGCACGTTGCT |
| R2-FP-2 | CTAGAAGTCGCGCTATTAACAGCTTGCTCAAGTGTGGCACTGCTCGAAGGGCTTCCTAAACTCAAATTAGCAACGTGCATGCCATTGTTC |
| R2-FP-3 | CAAGCTGTTAATAGCGCGACTTCTAGAGGGGTTCTTGTTGTAGCGGCATCTGGGAATTCAGGTGCAGGCTCAATCAGCTATCCGGCCCGT |
| R2-FP-4 | CCTGCGCCATACTGTGAAAAGCTGGCGCGGTTGTTGTTTTGGTCAGTCGCTCCGACTGCCATTGCGTTCGCATAACGGGCCGGATAGCTG |
| R2-FP-5 | GCTTTTCACAGTATGGCGCAGGGCTTGACATTGTCGCACCAGGTGTAAACGTGCAGAGCACACTGCCAGGTTCAGCGTATGCCAGCTTAA |
| R2-FP-6 | TTGTTTAACAAGGGCTGCTGCACCTGCAACATGAGGCGCAGCCATCGATGTACCGCTTAAGCTGGCATACGCTGAACCTG |
| R2-FP230-F | GCCCTTGTTAAACAAAAGAACCCATC |
| R2-FP-95-R | CCCTAATACTTTAACAGCGTATAGTTCCGCG |

**Supplementary Table 6 | Primers for constructing the library in the second round.**

| Primer | Sequence (5' → 3') |
|---|---|
| R2-1-1 | GAAGTAACGACAATGGCGCAAASCGTGCCATGGGGAATTAGCCGTGTGCAAGCCCCAGCTGCCCATAACCGTGGATTGAC |
| R2-1-2 | GAAGTAACGACAATGGCGCAAGTGGTGCCATGGGGAATTAGCCGTGTGCAAGCCCCAGCTGCCCATAACCGTGGATTGAC |
| R2-2 | TAAGTCTGGATGAGTGGAAATACCTGTATCGAGGACAGCAACTTTTACACCAGAACCTGTCAATCCACGGTTATGGGCAG |
| R2-3-1 | ACAGGTATTTCCACTCATCCAGACTTAAACATTCGTGGTGGCGCTAGCTTTGTACCAGGGGAACCATCCACTCAAGATGGGAATGGGCAT |

| Primer | Sequence (5' → 3') |
|---|---|
| R2-3-2 | ACAGGTATTTCCACTCATCCAGACTTARAAATTCGTGGTGGCGCTAGCTTTGTACCAGGGGAACCATCCACTCAAGATGGGAATGGGCAT |
| R2-3-3 | ACAGGTATTTCCACTCATCCAGACTTACGCATTCGTGGTGGCGCTAGCTTTGTACCAGGGGAACCATCCACTCAAGATGGGAATGGGCAT |
| R2-4 | CTCGGCGCTACGCCAAGAACGCCAATCGAATTGTYTAAAGCAGCAATCGTCCCGGCCACATGCGTGCCATGCCCATTCCCATCTTGAGTG |
| R2-5-1 | CTTGGCGTAGCGCCGAGCGCCGAACTATACGCTGTTAAAGTATTAGGGAGCAACGGTTCAGGTTCGGTCAGCTCGATTGCCCAAGGATTG |
| R2-5-2 | CTTGGCGTAGCGCCGAGCAGCGAACTATACGCTGTTAAAGTATTAGGGAGCAACGGTTCAGGTTCGGTCAGCTCGATTGCCCAAGGATTG |
| R2-6 | GTGGCACTGCTCGAAGGGCTTCCTAAACTCAAATTAGCAACGTGCATGCCATTGTTCCCTGCCCATTCCAATCCTTGGGCAATCGAGCTG |
| R2-7 | CCCTTCGAGCAGTGCCACACTTGAGCAAGCTGTTAATAGCGCGACTTCTAGAGGGGTTCTTGTTGTAGCGGCATCTGGGAATTCAGGTGC |
| R2-8 | GGTCAGTCGCTCCGACTGCCATTGCGTTCGCATAACGGGCCGGATAGCTGATTGAGCCTGCACCTGAATTCCCAGATGCC |
| R2-9-1 | GCAGTCGGAGCGACTGACCAAAACAACRACCGCGCCAGCTTTTCACAGTATGGCGCCGGGCTTGACATTGTCGCACCAGGTGT |
| R2-9-2 | GCAGTCGGAGCGACTGACCAAAACAACRACCGCGCCAGCTTTTCACAGTATGGCAGCGGGCTTGACATTGTCGCACCAGGTGT |
| R2-9-3 | GCAGTCGGAGCGACTGACCAAAACAACCAGCGCGCCAGCTTTTCACAGTATGGCGCCGGGCTTGACATTGTCGCACCAGGTGT |
| R2-9-4 | GCAGTCGGAGCGACTGACCAAAACAACCAGCGCGCCAGCTTTTCACAGTATGGCAGCGGGCTTGACATTGTCGCACCAGGTGT |
| R2-10 | ATCGATGTACCGCTTAAGCTGGCATACGCTGAACCTGGCAGTGTGCTCTGCACGTTTACACCTGGTGCGACAATGTCAAG |
| R2-11 | CCAGCTTAAGCGGTACATCGATGGCTGCGCCTCATGTTGCAGGTGYGGCAGCCCTTGTTAAACAAAAGAACCCATCTTGGTCCAAT |
| R2-12-1 | GTACAGGTTCGTGCTCCCCAGTTTCGTTGCCGTATTCTTTAGATGATTGCGGATTTGTACATTGGACCAAGATGGGTTCTTTTGTTTAAC |
| R2-12-2 | GTACAGGTTCGTGCTCCCCAGGCYCGTTGCCGTATTCTTTAGATGATTGCGGATTTGTACATTGGACCAAGATGGGTTCTTTTGTTTAAC |
| R2-12-3 | GTACAGGTTCGTGCTCCCCAGATWCGTTGCCGTATTCTTTAGATGATTGCGGATTTGTACATTGGACCAAGATGGGTTCTTTTGTTTAAC |
| R2-13-1 | CTGGGGAGCACGAACCTGTACGGAAGCGGACTTGTCAATGCASAGGCGGCAACACGCTAACTTCAGCACAATTCCAAGAAAGACACGA |

| Primer | Sequence (5' → 3') |
|---|---|
| R2-13-2 | CTGGGGAGCACGAACCTGTACGGARCCGGACTTGTCAATGCASAGGCG GCAACACGCTAACTTCAGCACAATTCCAAGAAAGACACGA |
| mp-R | TCGTGTCTTTCTTGGAATTGTGCTGAAG |
| R2-ProD-R | CATTGTCGTTACTTCTGCATCCTCTTC |
| R2-(-mp)-F | CCAAGAAAGACACGATTTAGAACCTAAAAAGAAC |

**Supplementary Table 7 | Primers for constructing the savinase libraries in the third round.**

| Primer / promoter | Sequence (5' → 3') |
|---|---|
| amyE-5'-F | TCGGTACCTCGCGAATGCATCTAGATATCGGATCCGCGGCATTA TGTTTGAATTTCCGTTTAAAG |
| amyE-5'-R | CTTGAGACCTTTGAGCTTCCGAGACTGGTCTCATCTTGACACTC CTTATTTGATTTTTTGAAGAC |
| amyE-3'-F | AAGATGAGACCAGTCTCGGAAGCTCAAAGGTCTCAAGCAAGTG CCCGGTCAGAATC |
| amyE-3'-R | ACAGGAAACAGCTATGACCATGATTACGCCAAGCTTCTTCACTA ACGATGCCTTTGAAAATCTTC |
| Sav-F | CGAAGTAAGTCTTCAAAAAATCAAATAAGGAGTGTCAAGAATGAA GAAACCGTTGGGGAAAATTG |
| Sav-R | AGTTGGGTAACGCCAGGGTTTTCCCAGTCACGACGTTATACACA AAAAAAC GCTGTGCCC |
| Cm-F | AACGTCGTGACTGGGAAAACC |
| Cm-R | AGCCAGGCTGATTCTGACCGGGCACTTGGGCGCTTTATTGGTAT GACTGGTTTTAAGCGC |
| Promoter | GTAAGCGTTAACAAAATTCTCCAGTCTTCACATCGGTTTGAAAGG AGGAAGCGGAAGAATGAAGTAAGAGGGATTTTTGACTCCGAAGT AAGTCTTCAAAAAATCAAATAAGGAGTGTCAAGA |
| Terminator | TAATCAATAAAAAAACGCTGTGCGGTTAAAGGGCACAGCGTTTT TTTGTGTAT |
| Signaling peptide (MKKPLGKIVASTALLI SVAFSSSIASA) | ATGAAGAAACCGTTGGGGAAAATTGTCGCAAGCACCGCACTACT CATTTCTGTTGCTTTTAGTTCATCGATCGCATCGGCT |

### The DNA sequence of savinase plasmid in the first and second rounds

PWB980ori-p43-SacB-Savinase

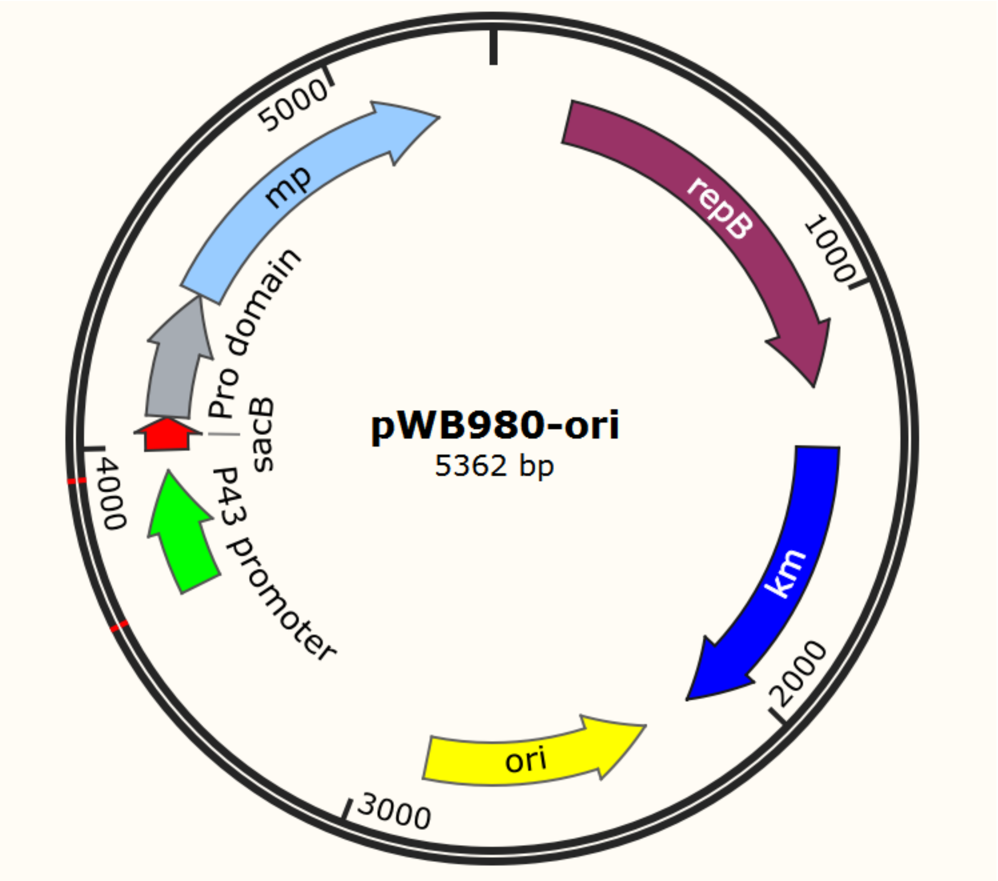


> P43 promoter

GAGCTCAGCATTATTGAGTGGATGATTATATTCCTTTTGATAGGTGGTATGTTTTCG
CTTGAACTTTTAAATACAGCCATTGAACATACGGTTGATTTAATAACTGACAAACAT
CACCCTCTTGCTAAAGCGGCCAAGGACGCTGCCGCCGGGGCTGTTTGCGTTTTTG
CCGTGATTTCGTGTATCATTGGTTTACTTATTTTTTTGCCAAAGCTGTAATGGCTGA
AAATTCTTACATTTATTTTACATTTTTAGAAATGGGCGTGAAAAAAAGCGCGCGATT
ATGTAAAATATAAAGTGATAGCGGTACC

> SacB

ATGAACATCAAAAAGTTTGCAAAACAAGCAACAGTATTAACCTTTACTACCGCACTG
CTGGCAGGAGGCGCAACTCAAGCTTTTGCC

> Prodomain

ATGAAGAAACCGTTGGGGAAAATTGTCGCAAGCACCGCACTACTCATTTCTGTTGC
TTTTAGTTCATCGATCGCATCGGCTGCTGAAGAAGCAAAAGAAAAATATTTAATTGG
CTTTAATGAGCAGGAAGCTGTCAGTGAGTTTGTAGAACAAGTAGAGGCAAATGACG
AGGTCGCCATTCTCTCTGAGGAAGAGGAAGTCGAAATTGAATTGCTTCATGAATTT

GAAACGATTCCTGTTTTATCCGTTGAGTTAAGCCCAGAAGATGTGGACGCGCTTGA
ACTCGATCCAGCGATTTCTTATATTGAAGAGGATGCAGAAGTAACGACAATG

> Mp

GCGCAATCAGTGCCATGGGGAATTAGCCGTGTGCAAGCCCCAGCTGCCCATAACC
GTGGATTGACAGGTTCTGGTGTAAAAGTTGCTGTCCTCGATACAGGTATTTCCACT
CATCCAGACTTAAATATTCGTGGTGGCGCTAGCTTTGTACCAGGGGAACCATCCAC
TCAAGATGGGAATGGGCATGGCACGCATGTGGCCGGGACGATTGCTGCTTTAAAC
AATTCGATTGGCGTTCTTGGCGTAGCGCCGAGCGCGGAACTATACGCTGTTAAAG
TATTAGGGGCGAGCGGTTCAGGTTCGGTCAGCTCGATTGCCCAAGGATTGGAATG
GGCAGGGAACAATGGCATGCACGTTGCTAATTTGAGTTTAGGAAGCCCTTCGCCA
AGTGCCACACTTGAGCAAGCTGTTAATAGCGCGACTTCTAGAGGGGTTCTTGTTGT
AGCGGCATCTGGGAATTCAGGTGCAGGCTCAATCAGCTATCCGGCCCGTTATGCG
AACGCAATGGCAGTCGGAGCGACTGACCAAAACAACAACCGCGCCAGCTTTTCAC
AGTATGGCGCAGGGCTTGACATTGTCGCACCAGGTGTAAACGTGCAGAGCACATA
CCCAGGTTCAACGTATGCCAGCTTAAACGGTACATCGATGGCTACTCCTCATGTTG
CAGGTGCAGCAGCCCTTGTTAAACAAAAGAACCCATCTTGGTCCAATGTACAAATC
CGCAATCATCTAAAGAATACGGCAACGAGCTTAGGAAGCACGAACTTGTATGGAAG
CGGACTTGTCAATGCAGAAGCGGCAACACGCTAA

**The DNA sequence of savinase plasmid in the third round**

The CDS region of the protease is marked in blue.

> pUC57-Sav-Cm-amyE

TCGCGCGTTTCGGTGATGACGGTGAAAACCTCTGACACATGCAGCTCCCGGAGAC
GGTCACAGCTTGTCTGTAAGCGGATGCCGGGAGCAGACAAGCCCGTCAGGGCGC
GTCAGCGGGTGTTGGCGGGTGTCGGGGCTGGCTTAACTATGCGGCATCAGAGCA
GATTGTACTGAGAGTGCACCATATGCGGTGTGAAATACCGCACAGATGCGTAAGG
AGAAAATACCGCATCAGGCGCCATTCGCCATTCAGGCTGCGCAACTGTTGGGAAG
GGCGATCGGTGCGGGCCTCTTCGCTATTACGCCAGCTGGCGAAAGGGGGATGTG
CTGCAAGGCGATTAAGTTGGGTAACGCCAGGGTTTTCCCAGTCACGACGTTGTAA
AACGACGGCCAGTGAATTCGAGCTCGGTACCTCGCGAATGCATCTAGATATCGGA

TCCGCGGCATTATGTTTGAATTTCCGTTTAAAGAATGGGCTGCAAGCCTTGTGTTTT
TGTTCATCATTATCTTATATTACTGCATCAGGGCTGCGGCATCCGGAATGCTCATG
CCGAGAATAGACACCAAAGAAGAACTGCAAAAACGGGTGAAGCAGCAGCGAATAG
AATCAATTGCGGTCGCCTTTGCGGTAGTGGTGCTTACGATGTACGACAGGGGGAT
TCCCCATACATTCTTCGCTTGGCTGAAAATGATTCTTCTTTTTATCGTCTGCGGCGG
CGTTCTGTTTCTGCTTCGGTATGTGATTGTGAAGCTGGCTTACAGAAGAGCGGTAA
AAGAAGAAATAAAAAAGAAATCATCTTTTTTGTTTGGAAAGCGAGGGAAGCGTTCA
CAGTTTCGGGCAGCTTTTTTTATAGGAACATTGATTTGTATTCACTCTGCCAAGTTG
TTTTGATAGAGTGATTGTGATAATTTTAAATGTAAGCGTTAACAAAATTCTCCAGTCT
TCACATCGGTTTGAAAGGAGGAAGCGGAAGAATGAAGTAAGAGGGATTTTTGACTC
CGAAGTAAGTCTTCAAAAAATCAAATAAGGAGTGTCAAGAATGAAGAAACCGTTGG
GGAAAATTGTCGCAAGCACCGCACTACTCATTTCTGTTGCTTTTAGTTCATCGATCG
CATCGGCTGCTGAAGAAGCAAAAGAAAAATATTTAATTGGCTTTAATGAGCAGGAA
GCTGTCAGTGAGTTTGTAGAACAAGTAGAGGCAAATGACGAGGTCGCCATTCTCTC
TGAGGAAGAGGAAGTCGAAATTGAATTGCTTCATGAATTTGAAACGATTCCTGTTTT
ATCCGTTGAGTTAAGCCCAGAAGATGTGGACGCGCTTGAACTCGATCCAGCGATTT
CTTATATTGAAGAGGATGCAGAAGTAACGACAATG**GCGCAATCAGTGCCATGGGG**
**AATTAGCCGTGTGCAAGCCCCAGCTGCCCATAACCGTGGATTGACAGGTTCTGG**
**TGTAAAAGTTGCTGTCCTCGATACAGGTATTTCCACTCATCCAGACTTAAATATTC**
**GTGGTGGCGCTAGCTTTGTACCAGGGGAACCATCCACTCAAGATGGGAATGGGC**
**ATGGCACGCATGTGGCCGGGACGATTGCTGCTTTAAACAATTCGATTGGCGTTCT**
**TGGCGTAGCGCCGAGCGCGGAACTATACGCTGTTAAAGTATTAGGGGCGAGCG**
**GTTCAGGTTCGGTCAGCTCGATTGCCCAAGGATTGGAATGGGCAGGGAACAATG**
**GCATGCACGTTGCTAATTTGAGTTTAGGAAGCCCTTCGCCAAGTGCCACACTTGA**
**GCAAGCTGTTAATAGCGCGACTTCTAGAGGCGTTCTTGTTGTAGCGGCATCTGGG**
**AATTCAGGTGCAGGCTCAATCAGCTATCCGGCCCGTTATGCGAACGCAATGGCA**
**GTCGGAGCTACTGACCAAAACAACAACCGCGCCAGCTTTTCACAGTATGGCGCA**
**GGGCTTGACATTGTCGCACCAGGTGTAAACGTGCAGAGCACATACCCAGGTTCA**
**ACGTATGCCAGCTTAAACGGTACATCGATGGCTACTCCTCATGTTGCAGGTGCAG**
**CAGCCCTTGTTAAACAAAAGAACCCATCTTGGTCCAATGTACAAATCCGCAATCA**
**TCTAAAGAATACGGCAACGAGCTTAGGAAGCACGAACTTGTATGGAAGCGGACT**

**TGTCAATGCAGAAGCGGCAACACGCTAA**TAATCAATAAAAAAACGCTGTGCGGTT
AAAGGGCACAGCGTTTTTTTGTGTATAACGTCGTGACTGGGAAAACCCTGGCGTTA
CCCAACTTAATCGCCTTGCAGCACATCCCCCTTTCGCCAGCTGGCGTAATAGCGAA
GAGGCCCGCACCGATCGCCCTTCCCAACAGTTGCGCAGCCTGAATGGCGAATGG
CGCTAGCAGCACGCCATAGTGACTGGCGATGCTGTCGGAATGGACGACGGCAATA
GTTACCCTTATTATCAAGATAAGAAAGAAAAGGATTTTTCGCTACGCTCAAATCCTT
TAAAAAAACACAAAAGACCACATTTTTTAATGTGGTCTTTTATTCTTCAACTAAAGCA
CCCATTAGTTCAACAAACGAAAATTGGATAAAGTGGGATATTTTTAAAATATATATTT
ATGTTACAGTAATATTGACTTTTAAAAAAGGATTGATTCTAATGAAGAAAGCAGACA
AGTAAGCCTCCTAAATTCACTTTAGATAAAAATTTAGGAGGCATATCAAATGAACTT
TAATAAAATTGATTTAGACAATTGGAAGAGAAAAGAGATATTTAATCATTATTTGAAC
CAACAAACGACTTTTAGTATAACCACAGAAATTGATATTAGTGTTTTATACCGAAAC
ATAAAACAAGAAGGATATAAATTTTACCCTGCATTTATTTTCTTAGTGACAAGGGTG
ATAAACTCAAATACAGCTTTTAGAACTGGTTACAATAGCGACGGAGAGTTAGGTTAT
TGGGATAAGTTAGAGCCACTTTATACAATTTTTGATGGTGTATCTAAAACATTCTCT
GGTATTTGGACTCCTGTAAAGAATGACTTCAAAGAGTTTTATGATTTATACCTTTCT
GATGTAGAGAAATATAATGGTTCGGGGAAATTGTTTCCCAAAACACCTATACCTGA
AAATGCTTTTTCTCTTTCTATTATTCCATGGACTTCATTTACTGGGTTTAACTTAAATA
TCAATAATAATAGTAATTACCTTCTACCCATTATTACAGCAGGAAAATTCATTAATAA
AGGTAATTCAATATATTTACCGCTATCTTTACAGGTACATCATTCTGTTTGTGATGGT
TATCATGCAGGATTGTTTATGAACTCTATTCAGGAATTGTCAGATAGGCCTAATGAC
TGGCTTTTATAATATGAGATAATGCCGACTGTACTTTTTACAGTCGGTTTTCTAACG
ATACATTAATAGGTACGAAAAAGCAACTTTTTTTGCGCTTAAAACCAGTCATACCAA
TAAAGCGCCCAAGTGCCCGGTCAGAATCAGCCTGGCTTTGATTACGTGCTAAATG
GTTTATATAATGACTCGGGCTTAAGCGGTTCTCTTCCCCATTGAGGGCAAGGCTAG
ACGGGACTTACCGAAAGAAACCATCAATGATGGTTTCTTTTTTGTTCATAAATCAGA
CAAAACTTTTCTCTTGCAAAAGTTTGTGAAGTGTTGCACAATATAAATGTGAAATAC
TTCACAAACAAAAAGACATCAAAGAGAAACATACCCTGGAAGGATGATTAATGATG
AACAAACATGTAAATAAAGTAGCTTTAATCGGAGCGGGTTTTGTTGGAAGCAGTTAT
GCATTTGCGTTAATTAACCAAGGAATCACAGATGAGCTTGTGGTCATTGATGTAAAT
AAAGAAAAAGCAATGGGCGATGTGATGGATTTAAACCACGGAAAGGCGTTTGCGC

CACAACCGGTCAAAACATCTTACGGAACATATGAAGACTGCAAGGATGCTGATATT
GTCTGCATTTGCGCCGGAGCAAACCAAAAACCTGGTGAGACACGCCTTGAATTAG
TAGAAAAGAACTTGAAGATTTTCAAAGGCATCGTTAGTGAAGAAGCTTGGCGTAAT
CATGGTCATAGCTGTTTCCTGTGTGAAATTGTTATCCGCTCACAATTCCACACAACA
TACGAGCCGGAAGCATAAAGTGTAAAGCCTGGGGTGCCTAATGAGTGAGCTAACT
CACATTAATTGCGTTGCGCTCACTGCCCGCTTTCCAGTCGGGAAACCTGTCGTGCC
AGCTGCATTAATGAATCGGCCAACGCGCGGGGAGAGGCGGTTTGCGTATTGGGC
GCTCTTCCGCTTCCTCGCTCACTGACTCGCTGCGCTCGGTCGTTCGGCTGCGGCG
AGCGGTATCAGCTCACTCAAAGGCGGTAATACGGTTATCCACAGAATCAGGGGAT
AACGCAGGAAAGAACATGTGAGCAAAAGGCCAGCAAAAGGCCAGGAACCGTAAAA
AGGCCGCGTTGCTGGCGTTTTTCCATAGGCTCCGCCCCCCTGACGAGCATCACAA
AAATCGACGCTCAAGTCAGAGGTGGCGAAACCCGACAGGACTATAAAGATACCAG
GCGTTTCCCCCTGGAAGCTCCCTCGTGCGCTCTCCTGTTCCGACCCTGCCGCTTA
CCGGATACCTGTCCGCCTTTCTCCCTTCGGGAAGCGTGGCGCTTTCTCATAGCTCA
CGCTGTAGGTATCTCAGTTCGGTGTAGGTCGTTCGCTCCAAGCTGGGCTGTGTGC
ACGAACCCCCCGTTCAGCCCGACCGCTGCGCCTTATCCGGTAACTATCGTCTTGA
GTCCAACCCGGTAAGACACGACTTATCGCCACTGGCAGCAGCCACTGGTAACAGG
ATTAGCAGAGCGAGGTATGTAGGCGGTGCTACAGAGTTCTTGAAGTGGTGGCCTA
ACTACGGCTACACTAGAAGAACAGTATTTGGTATCTGCGCTCTGCTGAAGCCAGTT
ACCTTCGGAAAAAGAGTTGGTAGCTCTTGATCCGGCAAACAAACCACCGCTGGTA
GCGGTGGTTTTTTTGTTTGCAAGCAGCAGATTACGCGCAGAAAAAAAGGATCTCAA
GAAGATCCTTTGATCTTTTCTACGGGGTCTGACGCTCAGTGGAACGAAAACTCACG
TTAAGGGATTTTGGTCATGAGATTATCAAAAAGGATCTTCACCTAGATCCTTTTAAA
TTAAAAATGAAGTTTTAAATCAATCTAAAGTATATATGAGTAAACTTGGTCTGACAGT
TACCAATGCTTAATCAGTGAGGCACCTATCTCAGCGATCTGTCTATTTCGTTCATCC
ATAGTTGCCTGACTCCCCGTCGTGTAGATAACTACGATACGGGAGGGCTTACCATC
TGGCCCCAGTGCTGCAATGATACCGCGAGATCCACGCTCACCGGCTCCAGATTTA
TCAGCAATAAACCAGCCAGCCGGAAGGGCCGAGCGCAGAAGTGGTCCTGCAACTT
TATCCGCCTCCATCCAGTCTATTAATTGTTGCCGGGAAGCTAGAGTAAGTAGTTCG
CCAGTTAATAGTTTGCGCAACGTTGTTGCCATTGCTACAGGCATCGTGGTGTCACG
CTCGTCGTTTGGTATGGCTTCATTCAGCTCCGGTTCCCAACGATCAAGGCGAGTTA

CATGATCCCCCATGTTGTGCAAAAAAGCGGTTAGCTCCTTCGGTCCTCCGATCGTT
GTCAGAAGTAAGTTGGCCGCAGTGTTATCACTCATGGTTATGGCAGCACTGCATAA
TTCTCTTACTGTCATGCCATCCGTAAGATGCTTTTCTGTGACTGGTGAGTACTCAAC
CAAGTCATTCTGAGAATAGTGTATGCGGCGACCGAGTTGCTCTTGCCCGGCGTCA
ATACGGGATAATACCGCGCCACATAGCAGAACTTTAAAAGTGCTCATCATTGGAAA
ACGTTCTTCGGGGCGAAAACTCTCAAGGATCTTACCGCTGTTGAGATCCAGTTCGA
TGTAACCCACTCGTGCACCCAACTGATCTTCAGCATCTTTTACTTTCACCAGCGTTT
CTGGGTGAGCAAAAACAGGAAGGCAAAATGCCGCAAAAAAGGGAATAAGGGCGAC
ACGGAAATGTTGAATACTCATACTCTTCCTTTTTCAATATTATTGAAGCATTTATCAG
GGTTATTGTCTCATGAGCGGATACATATTTGAATGTATTTAGAAAAATAAACAAATA
GGGGTTCCGCGCACATTTCCCCGAAAAGTGCCACCTGACGTCTAAGAAACCATTAT
TATCATGACATTAACCTATAAAAATAGGCGTATCACGAGGCCCTTTCGTC
